# From diet to defense: serpin duplication and gene-expression evolution as putative contributors to poison-frog alkaloid sequestration

**DOI:** 10.64898/2026.07.13.737953

**Authors:** Jeffrey L. Coleman, Vincent S. Le, Genrietta Yagudayeva Rozenberg, Md Abu Bakar Siddique, Mónica I. Páez-Vacas, David Salazar-Valenzuela, Matthew H. Dixon, Ian M. Riddington, Nicolás Del Castillo-Aguilera, Malki Bustos, Juan C. Santos, Rebecca L. Young, David C. Cannatella

## Abstract

Diverse organisms use toxins as antipredator defenses. Although many taxa synthesize their toxins, dietary toxin acquisition is less documented, particularly in vertebrates. In poison frogs (Dendrobatidae), efficient sequestration of dietary alkaloids evolved at least three times from ancestors that bore at most trace concentrations of skin alkaloids. Yet molecular mechanisms underlying poison-frog sequestration remain poorly understood. We used two approaches to address this. First, we performed a broad phylogenetic analysis of the serpinA gene family, which includes *serpina1*-like/alkaloid-binding globulin (ABG), the putative dendrobatid alkaloid transporter, and biliverdin-binding serpins (BBSs), which bind and spectrally tune biliverdin in blue-green arboreal frogs. Second, we used an evolutionarily narrower but discovery-oriented liver gene-expression analysis, contrasting seven populations from two sequestering *Epipedobates* species, the most recent origin of sequestration among dendrobatids (<15 mya), with two trace-accumulating lineages, *Silverstoneia* and *Hyloxalus*. Phylogenetic analyses revealed extensive diversification of ABGs and BBS-like genes in Dendrobatidae, forming two and five major clades, respectively. These expansions reveal broader ligand-binding serpin diversity than previously recognized and indicate that alkaloid sequestration may involve multiple ABG paralogs. Gene-expression analyses revealed 23 candidate genes associated with small-molecule transport, xenobiotic metabolism, and immune response, including *serpina1*-like, which was overexpressed in *Epipedobates*. A co-expression network analysis independently placed 20 of the 23 candidates into four key modules (enriched for differentially-expressed genes, containing ≥1 candidate, and correlated with sequestration). Our findings point to ligand-binding-protein diversification and coordinated gene-expression changes as major contributors to the evolution of alkaloid defenses in poison frogs.

## Introduction

Plants, animals, fungi, and many microorganisms have evolved an astounding array of compounds that render them distasteful, harmful, or toxic to organisms that attempt to consume them. In many cases, defended organisms synthesize their toxins, whereas others acquire defensive compounds from environmental sources such as the diet or symbiotic microbes (Vaelli et al. 2020). After acquisition, these compounds are often stored unchanged, though some organisms chemically modify them for enhanced potency (Daly et al. 2003). Studies of diet-acquired defensive compounds in animals have focused largely on invertebrates (e.g., Pawlik 1993; Zvereva and Kozlov 2016; Dean and Prinsep 2017), including phytophagous insects (Opitz and Müller 2009) and nudibranch mollusks (McPhail et al. 2001).

Examples of dietary toxin acquisition in land vertebrates are comparatively sparse and taxonomically widespread. These include a polyphyletic assemblage of New Guinean birds from the families Ifritidae, Oriolidae, Oreoicidae, and Pachycephalidae (Bodawatta et al. 2024); the Asian natricine snake *Rhabdophis tigrinus* (Hutchinson et al. 2007); the North American garter snake *Thamnophis sirtalis* (Williams et al. 2004); the Japanese fire-bellied newt *Cynops pyrrhogaster* (Kudo et al. 2015, 2017; Nakazawa et al. 2026); putatively, two species of *Lyciasalamandra* salamanders (Eleftherakos et al. 2024); and members from five families of frogs (Daly et al. 1994, 1997; Smith et al. 2002; Hantak et al. 2013; Rodríguez et al. 2013). In vertebrates, dietary toxin acquisition often evolves alongside antipredator signaling, including warning coloration (aposematism) and mimicry (Darst and Cummings 2006; Saporito et al. 2007a), and specialized defensive behaviors such as death-feigning (Fukada 1961; Mutoh 1983).

Of the vertebrates in which dietary toxin acquisition is documented, frogs are the best studied in terms of toxin diversity (Daly et al. 2005). Diet-derived skin alkaloids have been documented in diverse anuran families, namely members of Dendrobatidae (Daly et al. 1994), Mantellidae (*Mantella*) (Daly et al. 1997), Eleutherodactylidae (*Eleutherodactylus limbatus* group) (Rodríguez et al. 2013), Bufonidae (*Melanophryniscus*) (Hantak et al. 2013), and Myobatrachidae (*Pseudophryne*) (Smith et al. 2002). More than 1,400 alkaloids representing 24 structural classes (e.g., decahydroquinolines, pyridines, pumiliotoxins) have been identified in anurans (Daly et al. 2005; Tarvin et al. 2024). Most of these alkaloids appear to be derived from small leaf-litter arthropods, such as ants (Saporito et al. 2004), oribatid soil mites (Saporito et al. 2007b), millipedes (Saporito et al. 2003), and possibly melyrid beetles (Dumbacher et al. 2004).

Dendrobatids (Dendrobatoidea *sensu* Grant et al. [2006]) are an exceptional system for evolutionary studies given their recurrent origins of diet-acquired chemical defenses. In particular, members of three dendrobatid clades—*Epipedobates*, *Ameerega*, and Dendrobatinae (i.e., *Dendrobates sensu lato*, “*Colostethus*” *ruthveni*, and *Phyllobates*)—have independently evolved high diversities and concentrations of diet-derived alkaloids in the skin, producing exceptionally strong chemical defenses (Fig. 1a; Santos et al. 2003; Vences et al. 2003; Santos and Cannatella 2011; Santos et al. 2014). We use “sequestration” (Tarvin et al. 2024) to describe the ability to efficiently concentrate high levels of alkaloids in the skin, a phenotype suspected to be underpinned by active transport mechanisms (e.g., transporter proteins). Of the 359 dendrobatid species (AmphibiaWeb 2026), 107 belong to the three sequestering clades and are largely aposematic, whereas the remaining 252 species are, in almost all cases, inconspicuously colored and generally presumed to retain an ancestral phenotype characterized by low-level accumulation of alkaloids in the skin (Fig. 1a). Although we use “sequestering” for the derived phenotype and “trace-accumulating” for the ancestral phenotype for simplicity, this classification is mechanistically ambiguous. It is unknown whether trace-accumulating frogs acquire skin alkaloids from arthropod prey through passive diffusion coupled with reduced clearance, or through an uncharacterized incipient active transport system (see Daly 1998; Jeckel et al. 2026).

**Figure 1.**
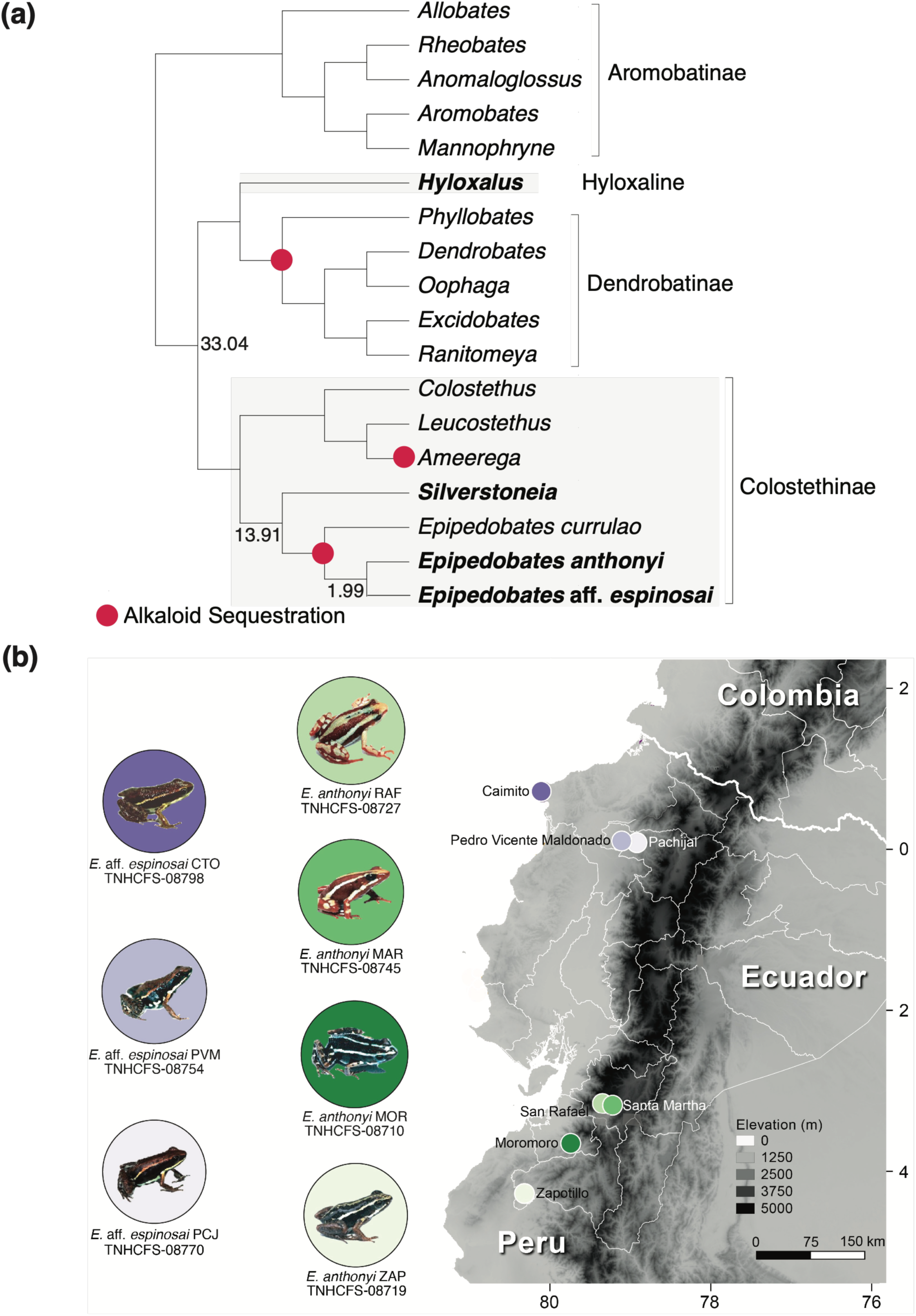
Panel (a); a cladogram of Dendrobatidae showing three independent appearances of alkaloid sequestration—once in Dendrobatinae and twice in Colostethinae (in *Epipedobates* and *Ameerega*) (Santos and Cannatella 2011; Santos et al. 2014; Tarvin et al. 2024). Genera sampled in this study have tip labels bolded, and divergence times among these genera are displayed, based on López-Hervas et al. (2024). Topology used is that of Grant et al. (2017). Panel (b); a topographic map of Ecuador showing locations from which *Epipedobates* sampled in this study were collected. *Hyloxalus awa* and *S. flotator* (both not shown), respectively, were collected near Pedro Vicente Maldonado and near Gamboa, Panamá (see Coleman et al. [2025] for map) (supplementary tables S3a, S3b). The abbreviations for localities, following the museum numbers, are used across figures and are as follows: CTO, Caimito; PVM, Pedro Vicente Maldonado; PCJ, Pachijal; RAF, San Rafael de Sharug; MAR, Santa Martha; MOR, Moromoro; ZAP, Zapotillo. For ease, future uses of abbreviations include a hyphenated species suffix (A for *E. anthonyi*, E for *E.* aff. *espinosai*). Photos are of *Epipedobates* field-collected in 2023 and were taken by J.L.C., N.D.C., and M.B. Alt text: Two-panel figure showing the phylogenetic and geographic context of the study. A dendrobatid cladogram marks three independent origins of alkaloid sequestration, and a topographic map shows sampled *Epipedobates* localities in Ecuador with representative frog photographs.

Regardless of mechanism, chemical surveys show that trace-accumulating dendrobatids in the wild typically harbor only a handful of alkaloids, whereas sympatric and syntopic species from sequestering lineages often contain tens to hundreds of alkaloids, with total concentrations of up to three orders of magnitude greater (Tarvin et al. 2024). Differences in foraging behavior or access to alkaloid-rich arthropods may contribute to this disparity and remain an important ecological axis of variation. However, experimental dosing studies indicate that dietary variation does not explain the difference. Under controlled exposure to the same alkaloids, sequestering frogs accumulate dietary alkaloids orders of magnitude more efficiently than trace-accumulating relatives (Jeckel et al. 2026). Earlier experiments using less sensitive analytical methods found little or no evidence of dietary alkaloid accumulation in trace-accumulating species (Daly et al. 1994; Saporito et al. 2009).

Identifying the molecular mechanisms of dendrobatid chemical defense is essential for understanding how the trait has evolved repeatedly across Dendrobatidae (e.g., Tarvin et al. 2016). More broadly, pinpointing these mechanisms can reveal whether independent origins of diet-acquired chemical defense in vertebrates arise through shared (parallel) or distinct (convergent) routes. Indeed, alkaloid sequestration is expected to be physiologically complex, requiring frogs to absorb alkaloids through the gut lining, bind and mobilize them into circulation within hours of consuming prey (O’Connell et al. 2021), and deliver them into cutaneous granular glands for storage (Neuwirth et al. 1979; Daly et al. 1980). Sequestration also requires mechanisms for tolerating or mitigating physiological impairment, such as reduced binding of alkaloids to sensitive target sites (Tarvin et al. 2016, 2017a), and can be accompanied by additional adaptations, including safe provisioning of alkaloids to larvae (Stynoski et al. 2014; Fischer et al. 2019) and, in some species, enzymatic potentiation of compounds for enhanced deterrence (Daly et al. 2003; Alvarez-Buylla et al. 2022). Chemical, genetic, physiological, and biochemical studies over the last decade have begun to reveal how dendrobatids acquire, transport, store, and tolerate defensive alkaloids. This work has uncovered new alkaloids in both aposematic and inconspicuously-colored dendrobatids (Tarvin et al. 2024), proteins involved in alkaloid transport (Alvarez-Buylla et al. 2023), and diverse mechanisms of autotoxicity avoidance (e.g., Tarvin et al. 2017a; Yeager et al. 2024).

Given the physiological breadth of alkaloid sequestration, it is unsurprising that previous studies have recovered large suites of proteins whose expression varies between sequestering and trace-accumulating species (Sanchez et al. 2019) and within sequestering species after experimental alkaloid exposure (e.g., Caty et al. 2019; O’Connell et al. 2021; Alvarez-Buylla et al. 2023). Some of these proteins appear to contribute to alkaloid bioconversion (Alvarez-Buylla et al. 2022), resistance (e.g., Tarvin et al. 2016, 2017a; Abderemane-Ali et al. 2021), and clearance (Sanchez et al. 2019). However, mechanisms that rapidly degrade or eliminate xenobiotics may also need to be reduced or bypassed in sequestering dendrobatids, enabling prolonged retention of alkaloids for defense (e.g., Caty et al. 2019; O’Connell et al. 2021; Alvarez-Buylla et al. 2023). For example, enzymes and transporter proteins in the vertebrate gut, including CYP3A and P-glycoprotein, often limit the passive diffusion of potentially toxic substances into the bloodstream (Zhang and Benet 2001). The modulation of such barriers may therefore be one route by which sequestering frogs increase alkaloid uptake while avoiding rapid detoxification or clearance.

In the sequestering dendrobatid species *Oophaga sylvatica*, Alvarez-Buylla et al. (2023) identified a liver-derived plasma protein central to alkaloid binding and transport. Plasma assays revealed that the protein, which the authors named the alkaloid-binding globulin (ABG), preferentially bound a photoprobe structurally similar to the common dendrobatid alkaloid pumiliotoxin (PTX) **251D**. ABG was found to be encoded by a *serpina1*-like gene (serpinA gene family). The putative homolog of the *O. sylvatica* ABG-encoding *serpina1*-like gene was recovered in two additional sequestering dendrobatid species, *Dendrobates tinctorius* and *Epipedobates tricolor*. *Oophaga sylvatica* and *D. tinctorius* belong to Dendrobatinae, the oldest origin of sequestration among dendrobatids (∼25 mya), whereas *Epipedobates* represents the most recent origin of this complex phenotype (∼12–15 mya) (Fig. 1a; Guillory et al. 2019; López-Hervas et al. 2024).

Recombinant-expression assays performed by the same authors showed that the putative ABG homologs from *O. sylvatica*, *D. tinctorius*, and *E. tricolor* can bind structurally distinct alkaloid classes and alkaloid-like ligands. However, because ABG was discovered using a PTX-like photoprobe, this approach may have preferentially recovered only one of multiple potential ABGs with affinity for PTX-like structures rather than the full repertoire of ABG paralogs that dendrobatids use to bind alkaloids. In addition, given the large numbers of proteins whose expression is correlated with sequestration in dendrobatids (e.g., Caty et al. 2019; O’Connell et al. 2021; Alvarez-Buylla et al. 2023), the sequestering phenotype likely depends not only on ABGs but also on additional molecular pathways involved in alkaloid uptake, transport, metabolism, and resistance, which have largely not been characterized outside of Dendrobatinae.

We performed two distinct yet complementary analyses: (1) we reconstructed the evolutionary history of serpinA, the gene family that includes ABG; (2) we performed a discovery-oriented differential gene expression (DGE) analysis to identify genes and pathways that may act alongside or independently of ABG, focusing on members of the *Epipedobates* origin of sequestration. For the second analysis, we contrasted seven populations from two *Epipedobates* species with two trace-accumulating lineages, *Silverstoneia flotator* and *Hyloxalus awa* (Fig. 1a,b). *Epipedobates* is a rapidly radiating clade with evidence of introgression and incomplete reproductive isolation (Tarvin et al. 2017b; López-Hervas et al. 2024), and all species examined possess moderate to high quantities of diverse skin alkaloids (Cipriani and Rivera 2009; Tarvin et al. 2024; Caty et al. 2025). Because the two *Epipedobates* species sampled for the DGE analysis diverged only ∼2 mya, we do not attempt to reconstruct ancestral expression states at the origin of sequestration. Instead, we use this comparative design to identify genes and pathways whose liver expression is associated with the sequestering phenotype.

## Results

### Evolutionary Reconstruction of the SerpinA Gene Family

Because one-to-one orthology between sequences from the serpinA1 subfamily in frogs and the canonical vertebrate *serpina1* ortholog (i.e., the human alpha-1-antitrypsin, or A1AT) remains unresolved, we use conservative terminology for members of the serpinA1 radiation in amphibians. Among neobatrachians (“modern” frogs), we refer to two broader serpinA1 gene groups: (1) *serpina1*-like, which contains the monophyletic, functionally-validated dendrobatid subgroup “ABGs” and (2) biliverdin-binding-serpin-like (BBS-like), which contains the validated, monophyletic hylid and centrolenid subgroup “BBSs.” We use “ancestral A1AT” for serpinA1 genes recovered from non-neobatrachian amphibians, namely archaeobatrachian frogs, salamanders, and caecilians (Figs. 2, 3).

**Figure 2.**
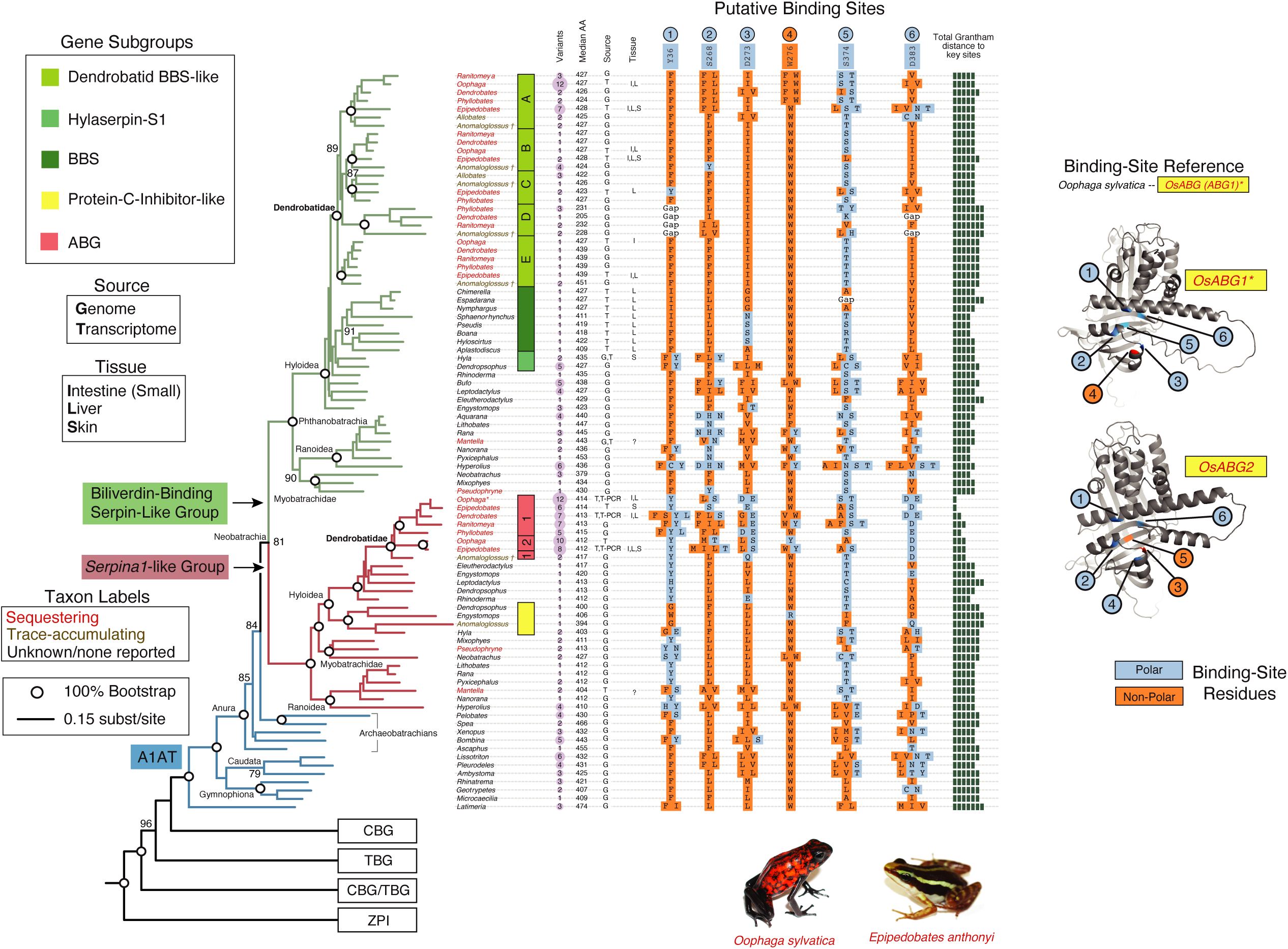
Maximum-likelihood phylogeny of amphibian serpinA genes inferred from a nucleotide alignment. Branch lengths are proportional to substitutions per site. Bootstrap support is shown only for selected deeper nodes to emphasize support for major serpinA relationships and focal gene groups; white circles indicate 100% bootstrap support, while other selected nodes with >75% support are shown numerically. Comprehensive support values are provided in supplementary fig. S2. Branch colors denote broad gene groups: biliverdin-binding serpin/biliverdin-binding-serpin-like genes (BBS/BBS-like; green), alkaloid-binding globulins (ABGs; red), and the ancestral alpha-1-antitrypsin-like lineage from which these groups diverged (blue). Colored boxes denote gene subgroups within the *serpina1*-like and BBS-like radiation. The additional canonical serpinA subfamilies collapsed at the bottom of the phylogeny are serpinA6/corticosteroid-binding globulin-like (CBG-like), serpinA6/7-like corticosteroid/thyroxine-binding globulin-like sequences (CBG/TBG-like), and serpinA10/ZPI-like sequences (see supplementary fig. S1 for the expanded tree). Five dendrobatid BBS-like clades (A–E) and two duplicated ABG clades (ABG1 and ABG2) were recovered. Tip-label colors indicate alkaloid phenotype/status, where red denotes alkaloid-sequestering taxa, brown denotes trace-accumulating taxa, and black denotes taxa for which alkaloid accumulation is unknown or no alkaloids have been confirmed (supplementary table S1b). AlphaFold models of *O. sylvatica* ABG1 and ABG2 show the structural positions of six residues near the predicted alkaloid-binding site and implicated by molecular docking in binding PTX **251D** in *O. sylvatica* (Alvarez-Buylla et al. 2023). Sequence columns summarize amino acid states at these six positions relative to *O. sylvatica* ABG1 (OsABG1) and classify residues by polarity. Total Grantham distance summarizes cumulative physicochemical divergence from OsABG1 across the six key sites. Additional columns indicate sequence source, transcriptomic tissue of origin (question mark indicates unknown or mixed), median protein length, and the maximum number of variants recovered per sampled species within each collapsed tip, rather than summed sequence counts across species. For sequence source, T-PCR denotes ABG-targeted PCR sequences generated by Alvarez-Buylla et al. (2023) from total RNA-derived cDNA. Variant numbers are backdropped with arbitrarily-colored circles (light magenta) whose sizes correspond to the numbers. Detailed criteria for variant designation are provided in supplementary table S2. * indicates the *O. sylvatica* ABG1 reference sequence used for residue comparisons. † *Anomaloglossus* has not been screened for alkaloids but is coded as trace-accumulating based on phylogenetic position within Aromobatinae (Tarvin et al. 2024) and consistent ecological traits, including inconspicuous coloration and a non-ant-and-mite-specialized diet reported for the congener *An. stepheni* (Lima and Moreira 1993; Juncá and Eterovick 2007). Alt text: A maximum-likelihood phylogeny distinguishing ancestral A1AT-like genes, dendrobatid ABG, and BBS-like genes, with residue, sequence-source, tissue, length, and variant annotations.

**Figure 3.**
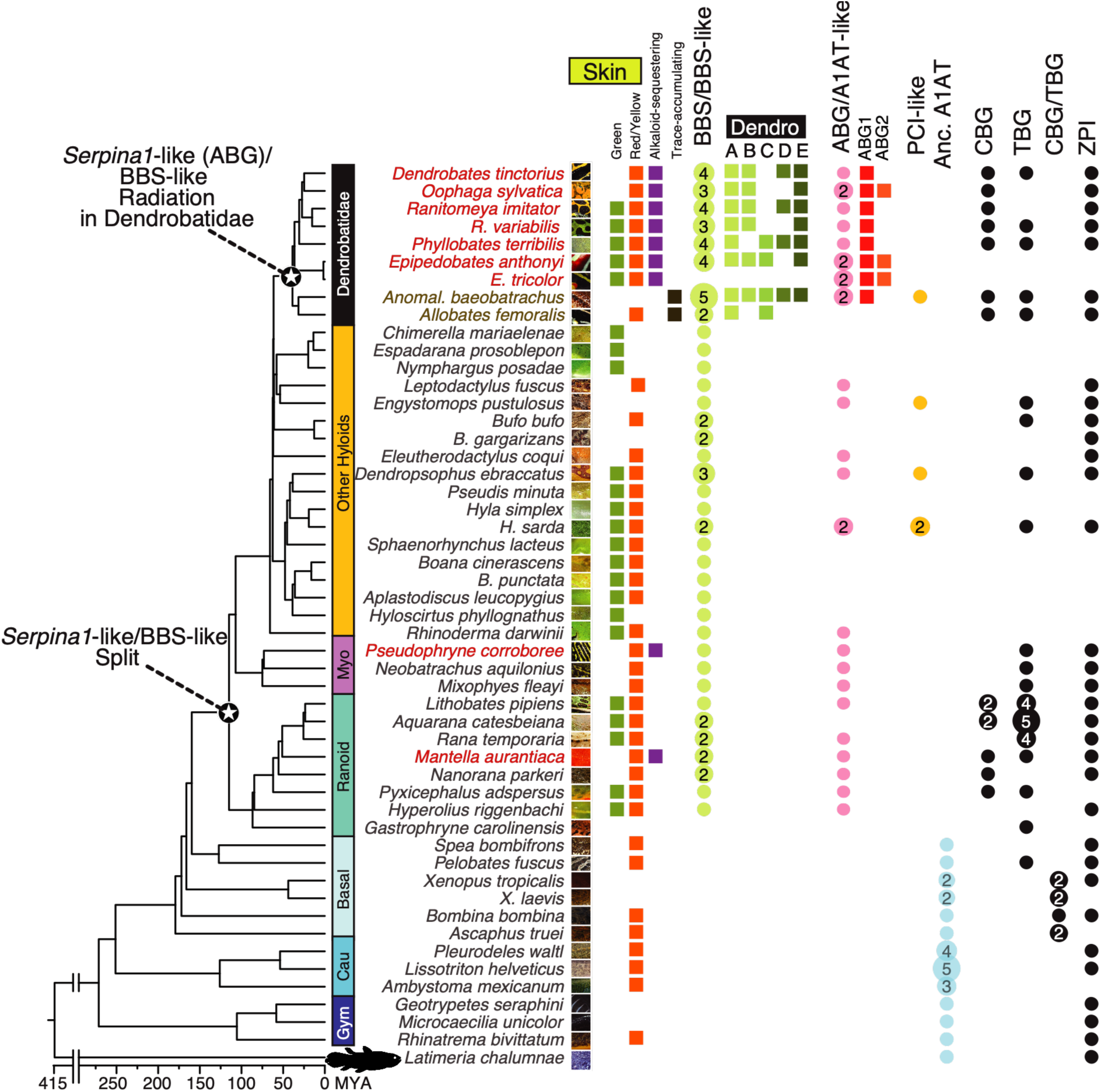
A time-calibrated amphibian species chronogram (after Portik et al. 2023) summarizing the distribution of recovered serpinA gene groups and subfamilies and selected phenotypes. Black stars indicate the inferred origin of the *serpina1*-like/BBS-like split near the base of modern frogs (Neobatrachia) and subsequent diversification of these groups within Dendrobatidae. Tip-label colors denote alkaloid phenotype: red, alkaloid-sequestering; brown, trace-accumulating; black/gray, unknown or no alkaloids confirmed (supplementary table S1b). Trait columns summarize green coloration, red/yellow coloration, alkaloid sequestration, and trace alkaloid accumulation. Gene columns indicate recovered sequence placements in named serpinA gene groups or subfamilies. For the dendrobatid ABG and BBS-like radiations, colored shapes denote named paralogous clades: ABG1–2 and BBS-like A–E. For other serpinA groups, numbers denote multiple distinct recovered phylogenetic lineages within a gene group for a given taxon; absence of a number indicates one recovered lineage. Dendrobatids exhibit exceptional recovered diversity across the ABG and BBS-like radiations relative to other sampled frogs. Alt text: A time-calibrated amphibian chronogram with trait and gene-presence columns, highlighting exceptional diversification of ABG and BBS-like genes in dendrobatids.

The phylogenetic reconstruction of the serpinA gene family from genomes and transcriptomes, both publicly available and newly generated (for *E. anthonyi* and *E. tricolor*), revealed that within Anura, the serpinA1 subfamily formed the two well-supported sister gene groups BBS-like and *serpina1*-like (Figs. 2, 3; supplementary figs. S1, S2; supplementary tables S1, S2). This split, which was followed by extensive gene diversification, appears to coincide with the major radiation of modern frogs (near the base of Neobatrachia, between the Neobatrachia and Phthanobatrachia nodes at ∼142–130 mya; Moen et al. 2016; Feng et al. 2017). At the origin of Dendrobatidae (∼38 mya), both BBS-like and *serpina1*-like experienced substantial diversification (Santos et al. 2009; Guillory et al. 2019). In contrast, the diversity of other serpin subfamilies (e.g., serpinA6, serpinA7, serpinA10) in dendrobatids was similar to that of non-dendrobatid frog lineages (Fig. 3).

Dendrobatid BBS-like genes we found were exceptionally diverse, forming at least five clades (designated BBS-like A–E) (Figs. 2, 3). Conservatively, we use “paralog” for BBS-like or *serpina1*-like sequences that were resolved in distinct clades in the gene tree, and “variant” for all distinct sequence records, or duplicated copies, of BBS-like or *serpina1*-like within a species. We recovered variant diversity within dendrobatid BBS-like clades that tended to be moderate and comparable to that of non-dendrobatids; only in a few genera did we find large numbers (i.e., *Oophaga* and *Epipedobates* BBS-like A) (Fig. 2). Interestingly, a trace-accumulating species (*Anomaloglossus baeobatrachus*) was the only dendrobatid from which we recovered paralogs from all five BBS-like clades. We found moderate diversity of BBS-like genes outside of Dendrobatidae, including three BBS-like lineages in *Dendropsophus* and one to six variants among hylids, leptodactylids, bufonids, Ranoidea, and Myobatrachidae (Figs. 2, 3). For hylids, putative defensive serpins (the hylaserpin-S1 group) (Wu et al. 2011) of *Hyla* and *Dendropsophus* were nested within the BBS-like gene group.

*Serpina1*-like genes showed limited recovered diversity across sampled Neobatrachia, with only one or two variants in most sampled lineages except dendrobatids and *Hyperolius* (Figs. 2, 3). Among dendrobatids, we recovered 5–12 variants from alkaloid-sequestering species, compared with 0 and 2 from the trace-accumulating species *Allobates femoralis* and *Anomaloglossus baeobatrachus*, respectively. These recovered variants fell into two distinct clades, which we designated ABG1 and ABG2; three alkaloid-sequestering species (*E. anthonyi*, *E. tricolor*, and *O. sylvatica*) were represented in both clades.

To validate transcript recovery and read support for these variants, we evaluated read-mapping metrics for all reconstructed serpinA sequences from the three sequestering dendrobatids represented only by transcriptome data (*E. anthonyi*, *E. tricolor*, and *O. sylvatica*) and compared these patterns against 11 housekeeping genes used as references (Supplementary Methods). In brief, the main limitation was not transcript recovery, but confident read assignment among closely related ABG/BBS-like sequences; this was consistent with expectations, given that we would predict multi-mapping of reads among closely related paralogs and transcript variants (supplementary tables S2a–c). High-quality read mapping to many reconstructed variants supports that these sequences are unlikely to represent simple assembly artifacts. However, these data do not by themselves distinguish among alternative isoforms, allelic variants, and recently duplicated gene copies. Closely related variants often differed at multiple amino acid sites in homologous aligned regions (supplementary table S1), a pattern more consistent with allelic variation or recent gene duplication followed by sequence divergence than with simple assembly redundancy (Pegueroles et al. 2013; Escorcia-Rodríguez et al. 2022). Interestingly, within both ABG1 and ABG2, individual sequence variants were often not species-specific: variants from different dendrobatid taxa were interspersed across subclades rather than forming fully species-monophyletic clusters. This pattern may reflect limited phylogenetic resolution among closely related, rapidly evolving variants, incomplete lineage sorting or shared ancestral polymorphism, or convergent sequence evolution at sites associated with alkaloid binding (supplementary fig. S2).

We found that the alkaloid-binding globulins identified by Alvarez-Buylla et al. (2023) in *O. sylvatica* (OsABG) and *D. tinctorius* (DtABG) are members of the ABG1 clade; we henceforth refer to ABGs by the clade to which they belong (i.e., OsABG1 and DtABG1). Variants of OsABG1 were recovered from liver and intestine transcriptomes, but not skin, of *O. sylvatica*. In contrast, skin-expressed sequences were overrepresented within the ABG2 clade, including multiple OsABG2 variants all from skin transcriptomes. This clade also includes EtABG2, previously identified as the *E. tricolor* ortholog of OsABG1; our reconstruction instead suggests that a second *E. tricolor* paralog, EtABG1, which nests within ABG1, is the more likely one-to-one ortholog of OsABG1 (supplementary fig. S2). Given these relationships, we suspect that EtABG1 is probably the primary liver-expressed serpin in *E. tricolor*, though sequence data from this species derive only from skin samples, so expression of EtABG1 in other tissues cannot yet be confirmed.

We also present AlphaFold structural predictions for OsABG1 and the newly identified OsABG2 (Fig. 2), which are broadly similar in overall structure but differ in polarity at residues corresponding to predicted alkaloid-binding sites in OsABG1, potentially favoring binding by distinct alkaloid classes (Supplementary Methods). Because OsABG1 uses a small-molecule-binding pocket whose architecture is conserved among Serpin A proteins, we compared six homologous residues across the family corresponding to OsABG1 positions predicted to contribute to PTX **251D** binding (based on molecular docking and subsequent site-directed mutagenesis; Alvarez-Buylla et al. 2023). We calculated the total Grantham distance (a measure of physicochemical dissimilarity) between each sequence and OsABG1 (Supplementary Methods). These analyses indicated pronounced sequence divergence between dendrobatid ABGs and non-dendrobatid *serpina1*-like, as well as between dendrobatid ABGs and non-dendrobatid members of other serpinA gene subfamilies (Fig. 2; supplementary table S1). We also observed substantial amino-acid variation across ABG1 and ABG2 at the six aligned positions corresponding to these OsABG1 putative binding sites (Fig. 2; supplementary table S1). Within dendrobatids, substitutions at site 6 (D383) uniformly retained polarity, whereas the other five sites included both polar and nonpolar residues. Polymorphism within these six homologous positions may alter binding-pocket chemistry, geometry, or ligand specificity.

In other amphibians (archaeobatrachian frogs, salamanders, and caecilians), serpinA1 sequences were not nested within either the *serpina1*-like or BBS-like groups. This pattern suggests that the *serpina1*-like and BBS-like clades recovered here are restricted to Neobatrachia (or possibly Phthanobatrachia). In contrast, serpinA1 variants corresponding to ancestral A1AT were recovered from archaeobatrachian anurans (*Spea*, *Pelobates*, *Bombina*, *Xenopus*, and *Ascaphus*), Caudata, and Gymnophiona (Figs. 2, 3). In the ancestral A1AT group we found modest diversity, with a single subgroup recovered in each of archaeobatrachians, salamanders, and caecilians, and low-to-moderate (1–6) copy numbers recovered among taxa (Figs. 2, 3).

More distant serpinA subfamilies sampled included serpinA6/corticosteroid-binding globulin (CBG), serpinA7/thyroxine-binding globulin (TBG), and serpinA10/protein Z-dependent protease inhibitor (ZPI) (Fig. 2). These subfamilies showed relatively low recovered diversity, with mostly one sequence per subfamily, notwithstanding the multiple TBG-like sequences discovered in the three sampled ranid species (*Lithobates pipiens*, *Aquarana catesbeiana*, and *Rana temporaria*) (Fig. 3; supplementary fig. S2). Some sequences from *Bombina*, *Xenopus*, and *Ascaphus* formed a monophyletic clade that could not be confidently assigned to either CBG or TBG; we therefore designated this clade CBG/TBG (Figs. 2, 3; supplementary figs. S1, S2; supplementary table S1). A small clade annotated as protein-C inhibitor (PCI; *serpina5* in humans) was nested within the ABG/BBS-containing *serpina1*-like radiation rather than with canonical vertebrate PCI sequences, so we refer to these sequences as “PCI-like”; this clade included one dendrobatid sequence from *Anomaloglossus baeobatrachus* and sequences from other hyloid frogs (Fig. 2; supplementary figs. S1, S2).

### Phenotype Verification, Annotation, and Mapping for the Gene-Expression Analysis

We next moved beyond the phylogenetic analysis to ask about broader molecular correlates of alkaloid sequestration, including resistance and metabolism. We carried out an untargeted gene-expression study comparing two *Epipedobates* species from the most recent origin of dendrobatid sequestration (<15 mya) with two trace-accumulating dendrobatid lineages outside *Epipedobates*: *Silverstoneia* and *Hyloxalus*. Because our gene-expression analyses relied on phenotypic contrasts between alkaloid-sequestering and trace-accumulating lineages, we first quantified alkaloid abundance and diversity in our samples of wild-caught frogs, to validate that alkaloid levels were consistent with the sequestering versus trace-accumulating categories. For *Epipedobates*, suffixes to population abbreviations denote species identity: A for *E. anthonyi* and E for *E.* aff. *espinosai*. The sequestering set included MOR-A, ZAP-A, RAF-A, MAR-A, PVM-E, PCJ-E, and CTO-E (N = 3–5 per population). Because only one population was sampled for both trace-accumulating species, we refer to those samples by species name rather than population abbreviation: *H. awa* (N = 4) and *S. flotator* (N = 3) (Fig. 1b; supplementary table S3a). We used the same individuals whenever possible for alkaloid profiling and transcriptomics; exceptions are noted in the Supplementary Methods, with further details in supplementary tables S3a and S3b. Stringent screening recovered 117 alkaloids representing 18 structural classes in *Epipedobates* but none in *S. flotator* or *H. awa*, validating the expected phenotypic categories (Fig. 4); however, low per-population sample sizes (mean = 3.9 ± 0.7 SD; N = 7 populations), substantial overlap in alkaloid classes, and modest among-population differences in total abundance prohibited well-powered tests of associations between individual alkaloid classes and gene expression (Supplementary Methods and Results; supplementary tables S4a–g; supplementary figs. S3, S4).

**Figure 4.**
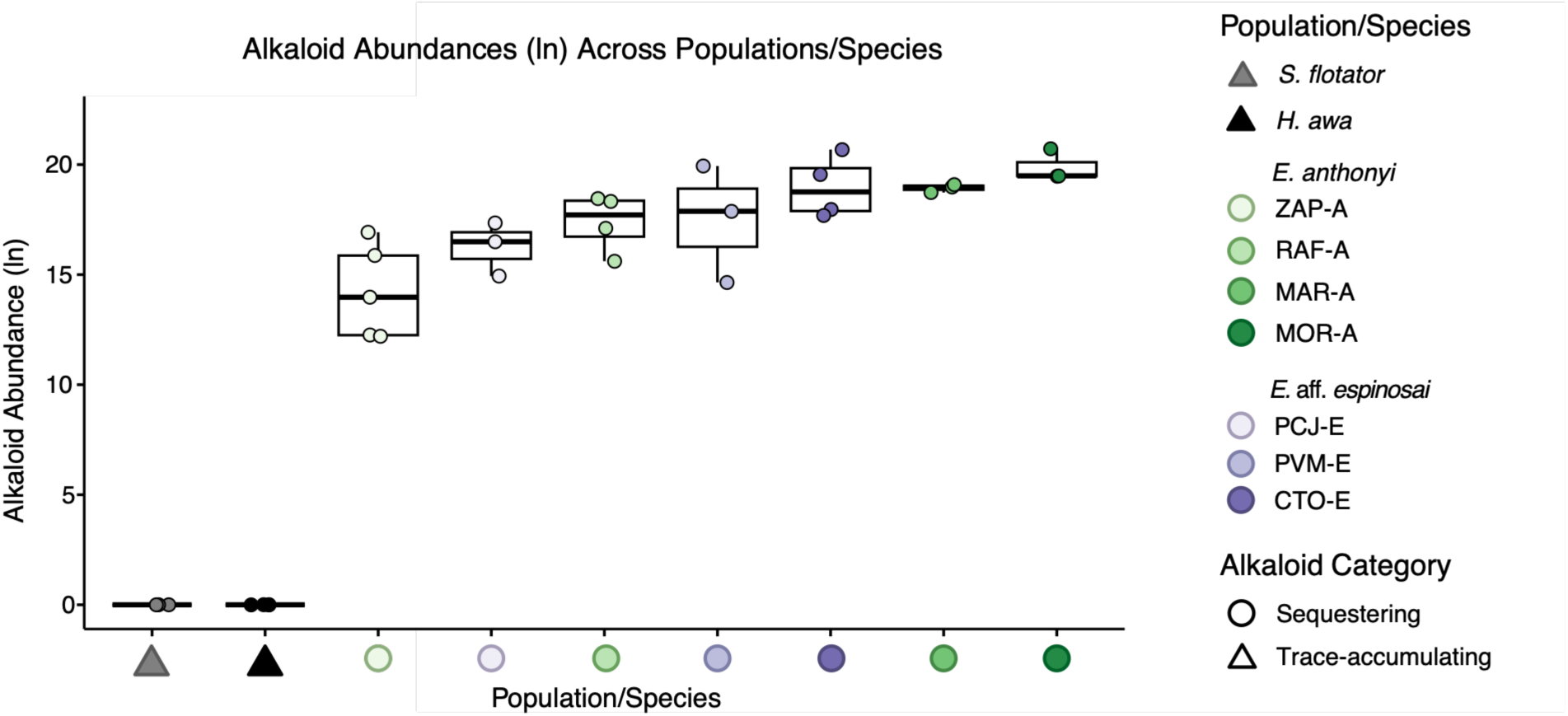
A boxplot showing alkaloid-abundance data (ln-transformed integrated areas) for all individuals (shown as jittered points and color-coded by population) profiled in this study, grouped by the two phenotypic categories into which the data naturally cluster (trace-accumulating and sequestering). We did not recover alkaloids in trace-accumulating species, so non-transformed values of 0 are plotted for all low-alkaloid individuals (see Supplementary Methods and Results). Samples are colored by population, with circles indicating sequestering species and triangles indicating trace-accumulating species. Alt text: Boxplot showing alkaloid abundance across trace-accumulating and sequestering frogs. Trace-accumulating species have zero detected alkaloids, whereas all *Epipedobates* populations show elevated alkaloid abundance with population-level variation.

We next reconstructed liver gene-expression patterns and evaluated their association with alkaloid sequestration. For this analysis, we sampled liver tissue from the four dendrobatid species. Gene expression profiling was completed using Tag-Seq, a high-throughput alternative to RNA-seq that concentrates sequencing effort on the 3′ end of mRNA (Meyer et al. 2011), thereby lowering the per-sample sequencing depth required (Lohman et al. 2016). We quantified all samples against a reference liver transcriptome from the sequestering dendrobatid *O. sylvatica* rather than using a single focal-species reference for all taxa, which could bias read recovery toward the reference lineage (Meyer et al. 2011; Lohman et al. 2016). We annotated the Caty et al. (2019) *O. sylvatica* liver transcriptome (file: *Osylvatica_transcriptome_Catyetal.fasta*) using Pincho (Ortiz et al. 2021) and standardized transcript/gene identifiers using the annotated *R. imitator* genome (aRanImi1; GCF_032444005.1) (Rhie et al. 2021) as an auxiliary reference, providing gene-level identifiers for downstream expression analyses. We then used the serpinA phylogenetic analysis to refine the gene identifiers for serpinA transcripts in the *O. sylvatica* reference, which had been included in the phylogenetic analysis. Transcripts corresponding to *serpina6*-like, *serpina10*-like, and *serpina1*-like were found in the *O. sylvatica* liver reference transcriptome. All *serpina1*-like isoforms were partial sequences identical to three ABG1 variants across the aligned region: two OsABG1-like sequences and one DtABG1-like sequence. *Serpina10*-like isoforms were partial or complete sequences identical to the single *O. sylvatica serpina10*-like variant across the aligned region. The *serpina6*-like isoforms were partial or complete sequences identical to one of the two *O. sylvatica serpina6*-like variants across the aligned region (supplementary fig. S2).

We then mapped Tag-Seq reads from the four focal species (*E. anthonyi*, *E.* aff. *espinosai*, *S. flotator*, and *H. awa*) to the *O. sylvatica* reference transcriptome. Because Tag-Seq quantification depends on consistent read mapping across samples, we calculated statistics to ensure uniform alignment percentages; an average of 5.57 million reads (±0.78 SD) were generated per sample, of which 39.09% (±4.33% SD) mapped to the reference, consistent with strong alignment rates for Tag-Seq studies (Meyer et al. 2011; Lohman et al. 2016). A one-way ANOVA detected a marginally significant effect of species on mapping percentage (F = 3.042, p = 0.0425), but Tukey HSD tests revealed no significant pairwise differences, indicating that mapping variation was not driven by any one species (supplementary fig. S5).

We first evaluated how consistently annotated genes in the *O. sylvatica* reference transcriptome received mapped reads across the focal species. Of the 12,763 uniquely annotated genes in the reference, 12,472 had ≥1 mapped read in at least one sample. The number of genes detected across all individuals declined modestly as more phylogenetically distant lineages were included: 3,733 genes were detected in all *Epipedobates* individuals, 3,516 of these were also detected in *S. flotator*, and 3,261 were detected across all samples (including *Epipedobates*, *S. flotator*, and *H. awa*). After applying expression filtering with filterByExpr in edgeR (Chen et al. 2025a), the 940 genes present in all focal species were retained for downstream analyses.

### Differentially Expressed Genes (DEGs) Associated with the Sequestering Phenotype Category

Before performing the DGE analyses, we conducted several principal component analyses (PCAs) to explore broad gene-expression patterns. PCA and k*-*means clustering revealed broad gene-expression structure corresponding to the three sampled genera while also distinguishing the two *Epipedobates* species. Phylogenetic PCAs revealed no disproportionate expression structure at the individual or population level but suggested substantial expression clustering by species that did not track phylogenetic relatedness (Supplementary Methods and Results; supplementary fig. S7a–k).

Because the sequestering samples represent a single origin of sequestration (*Epipedobates*), shared expression differences in *E*. aff. *espinosai* and *E. anthonyi* cannot be fully distinguished from lineage-specific changes unrelated to sequestration. We therefore used a conservative prioritization strategy to identify category-level expression differences most plausibly related to sequestration. First, we contrasted sequestering *Epipedobates* with a combined trace-accumulating category consisting of *S. flotator* and *H. awa*, retaining this combined category despite the broad global expression divergence between these two species evident in the PCA and pPCA analyses. Second, we screened the resulting DEGs for overlap with an *a priori* list of gene superfamilies implicated in toxin metabolism in poison-frog livers (supplementary table S5) and vertebrates more broadly (supplementary table S6). Genes meeting both criteria are hereafter designated candidate DEGs (Supplementary Methods).

We identified 401 DEGs between the trace-accumulating and sequestering categories (supplementary table S7), 23 of which (5.74%) were candidate DEGs (Supplementary Methods). Eleven candidate DEGs were broadly underexpressed in *Epipedobates* relative to the trace-accumulating species, whereas the remaining 12 were broadly overexpressed (Fig. 5; Table 2; Supplementary Methods). Although *c7* is not itself a member of the thioester-containing protein (TEP) superfamily, we retained it as a candidate DEG because of its functional interdependence and co-expression with complement genes *c3* and *c5* in the pathway leading to formation of the membrane attack complex (Table 1). Notably, two candidate DEGs (*apoh* and *hspa8*) were among the top 20 loadings contributing to PC1 from the non-phylogenetic PCA (supplementary fig. S8). No significant correlations were detected between expression of any of the 23 candidate DEGs and alkaloid abundance among *Epipedobates* individuals (supplementary fig. S9).

**Figure 5.**
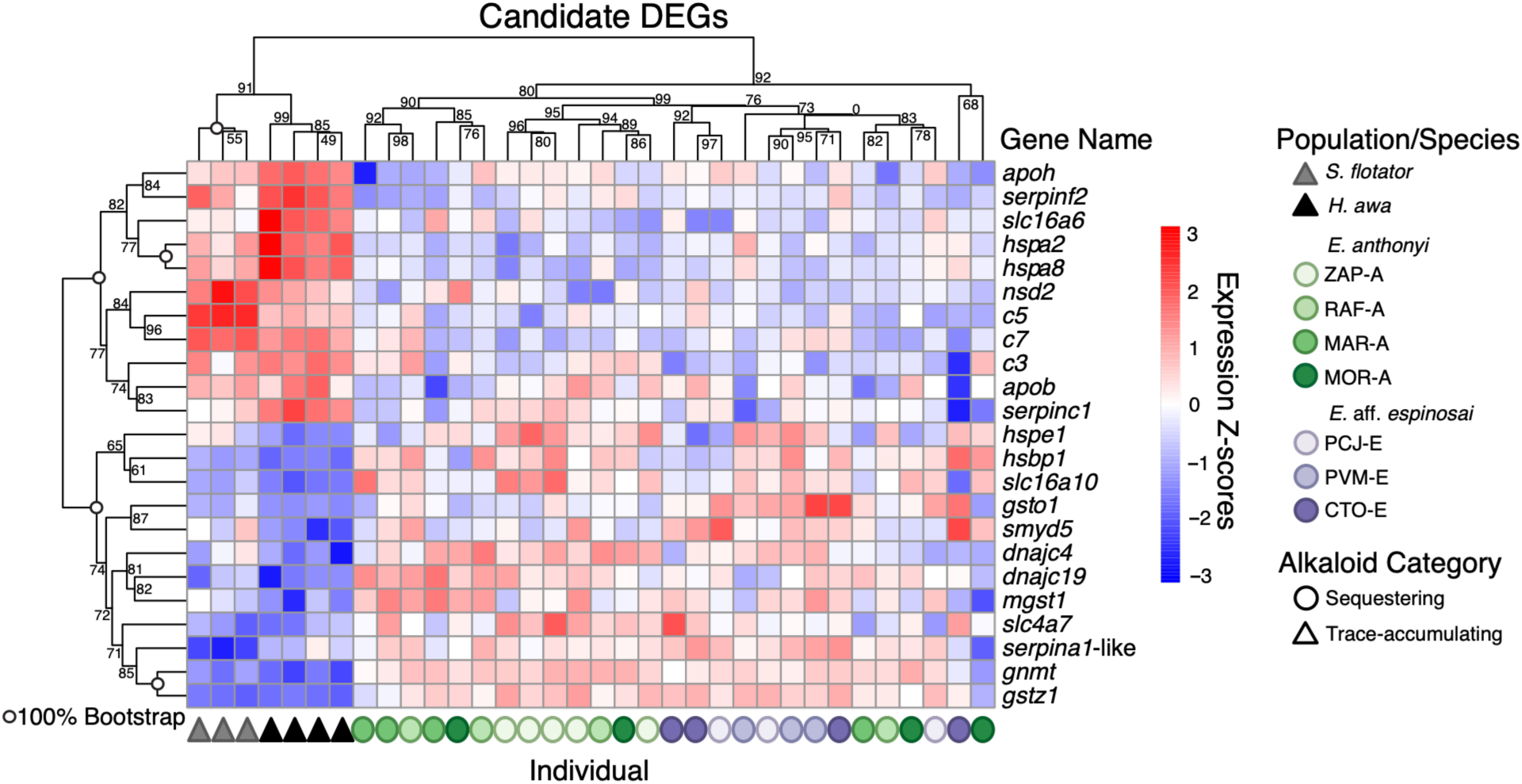
Heatmap of candidate genes differentially expressed (adjusted p-value ≤ 0.05) between sequestering and trace-accumulating lineages. Values represent gene expression centered and scaled per gene (row Z-scores), with color indicating relative expression (blue = lower, red = higher). Dendrograms for both rows (genes) and columns (individuals) are hierarchically clustered based on expression similarity. Node values show AU support from *pvclust* version 2.2-0 multiscale bootstrap resampling (Suzuki and Shimodaira 2006). Samples are colored by population, with circles indicating sequestering species and triangles indicating trace-accumulating species. See details on heatmap construction in Supplementary Methods. Alt text: Heatmap of differentially-expressed candidate genes showing expression clustering across individuals. Trace-accumulating species cluster together and show distinct expression patterns from *Epipedobates* populations for small-molecule transport and metabolism genes (including *serpina1*-like/ABG) and stress-response genes.

**Table 1.**
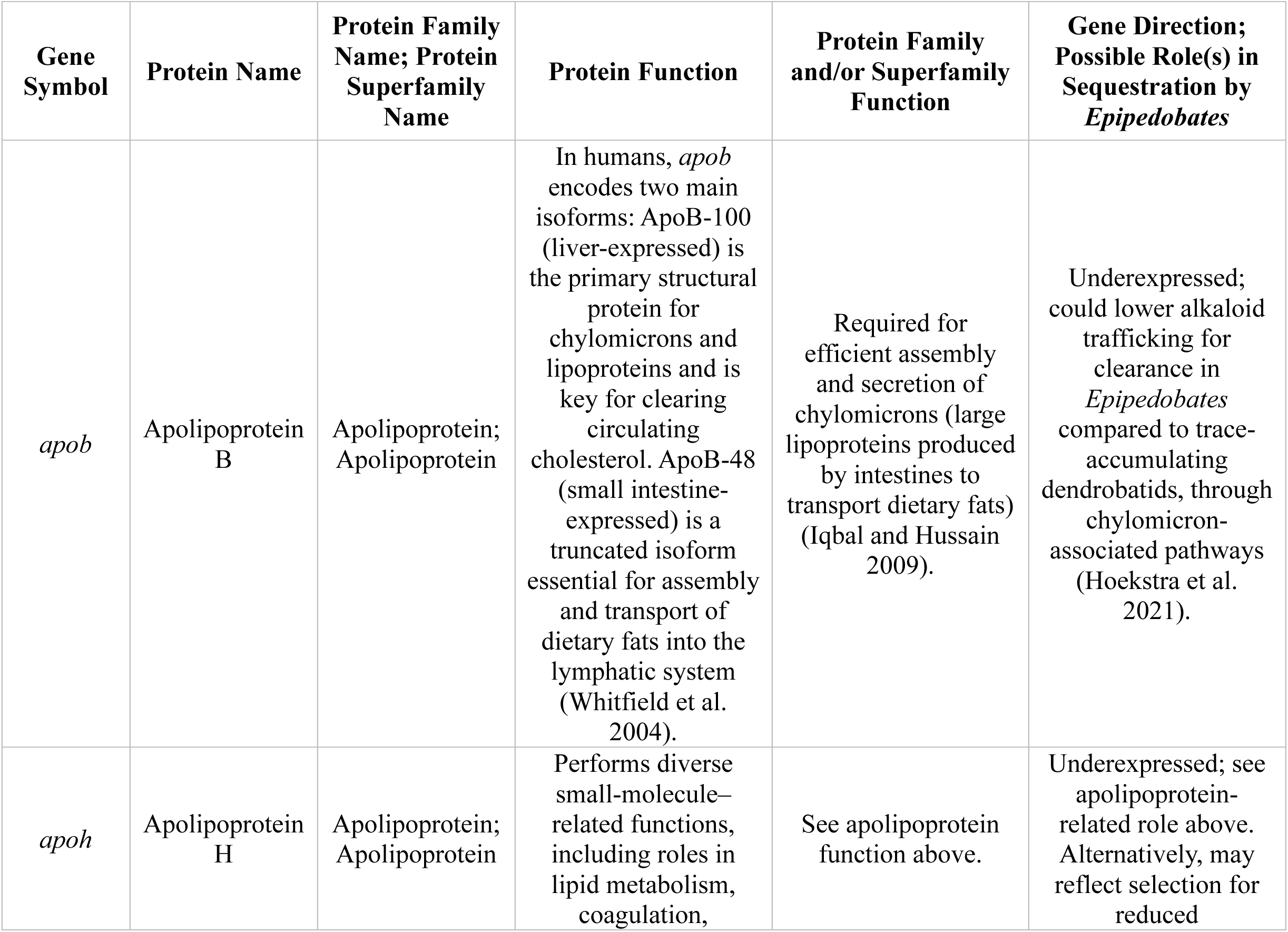
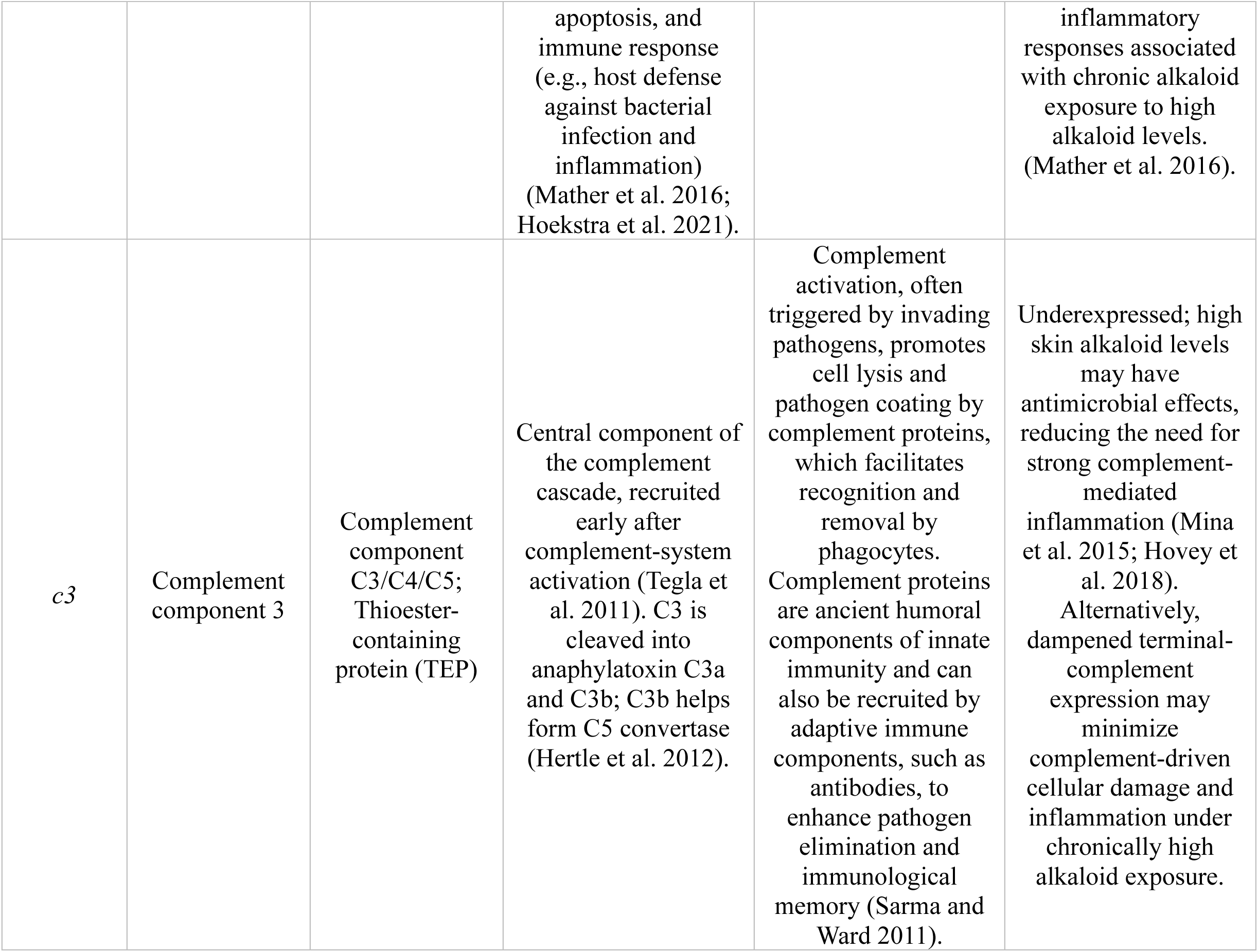
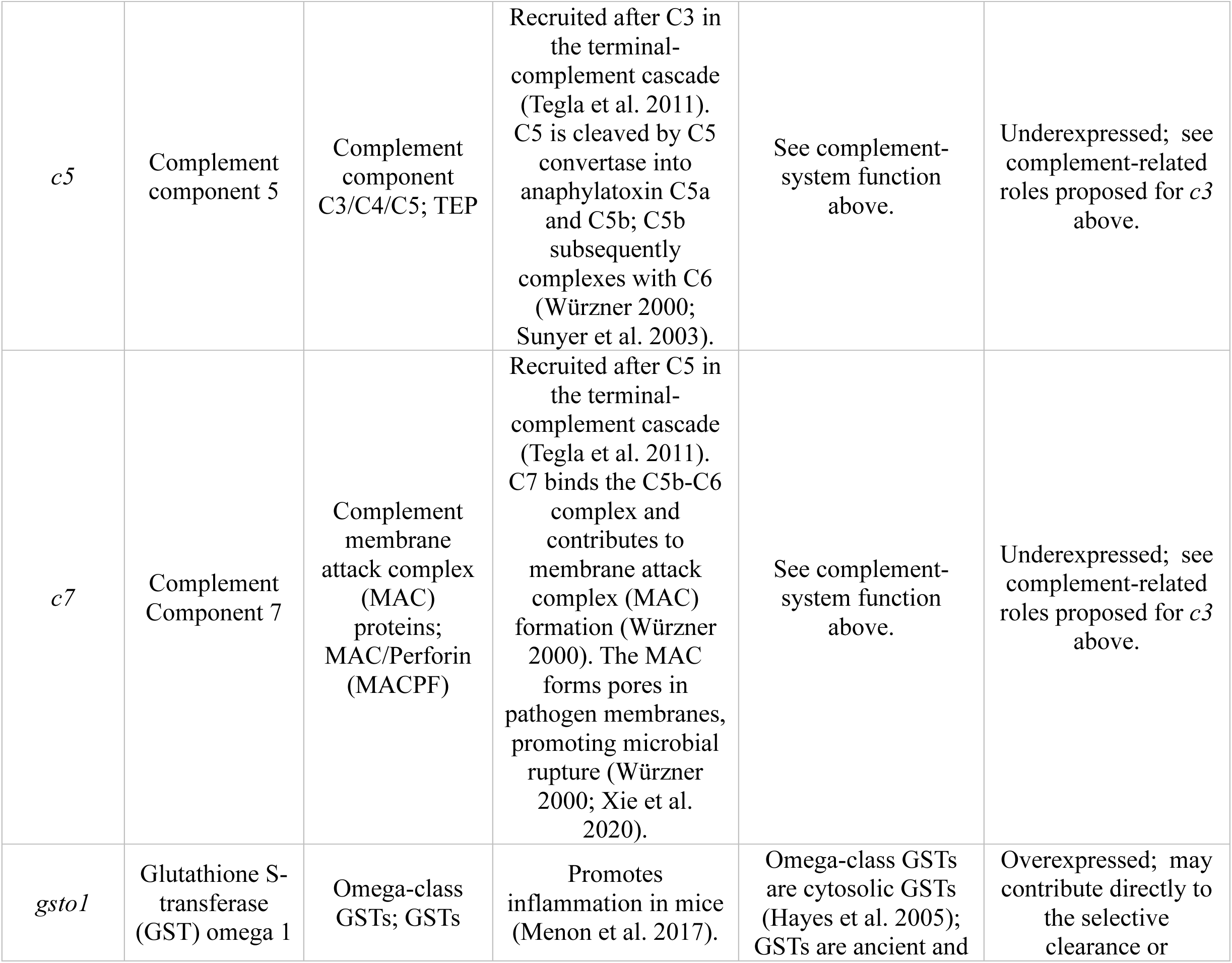
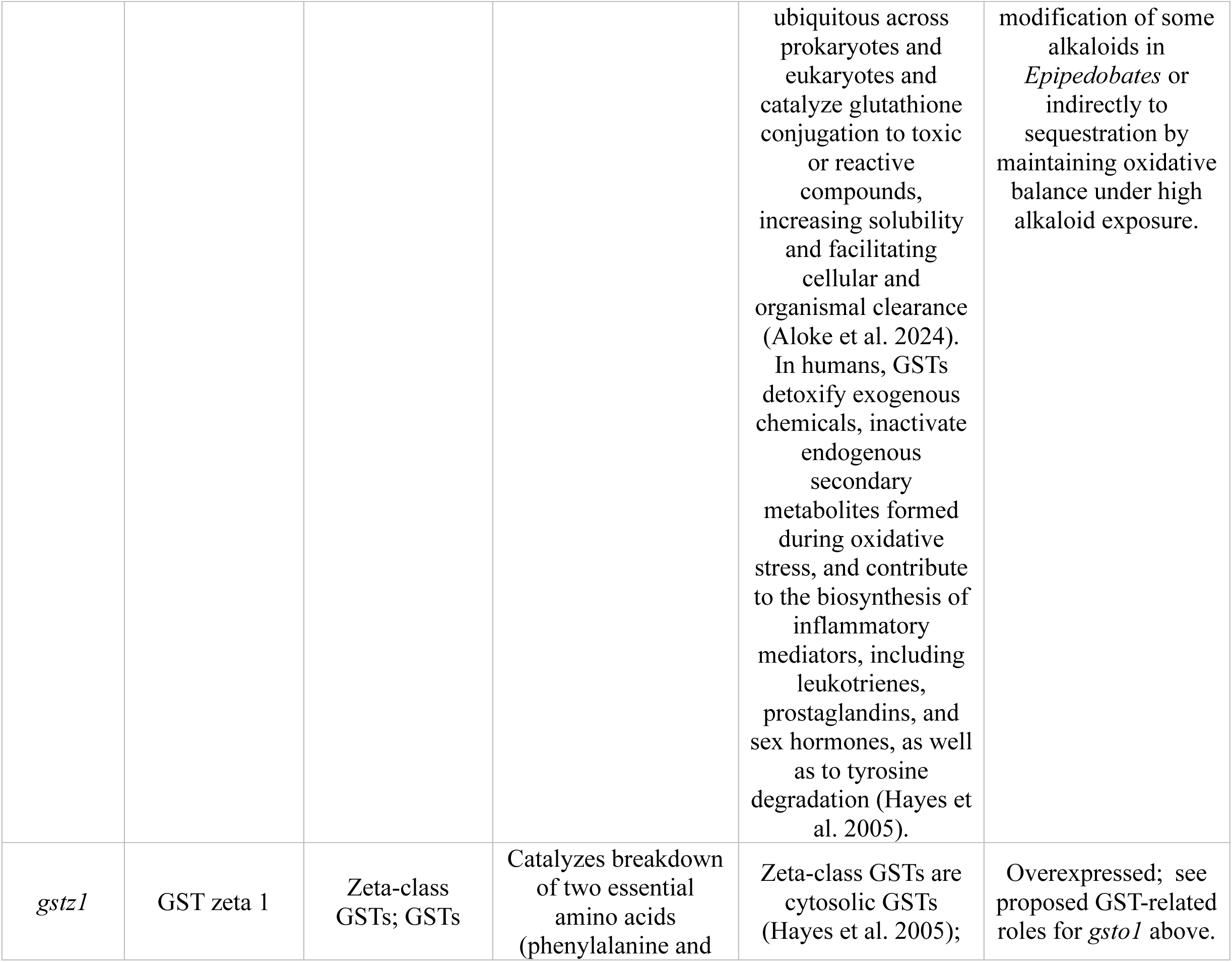
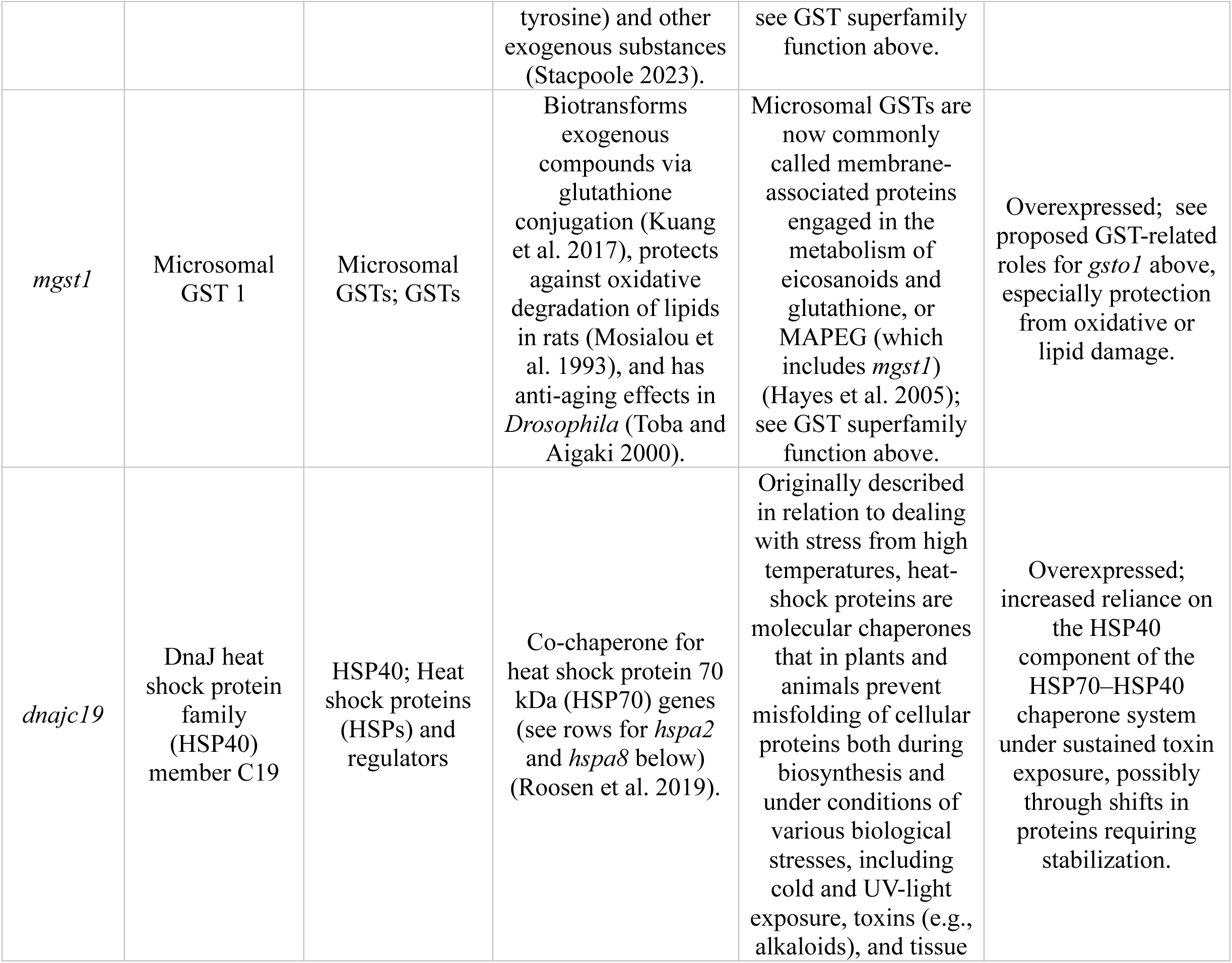
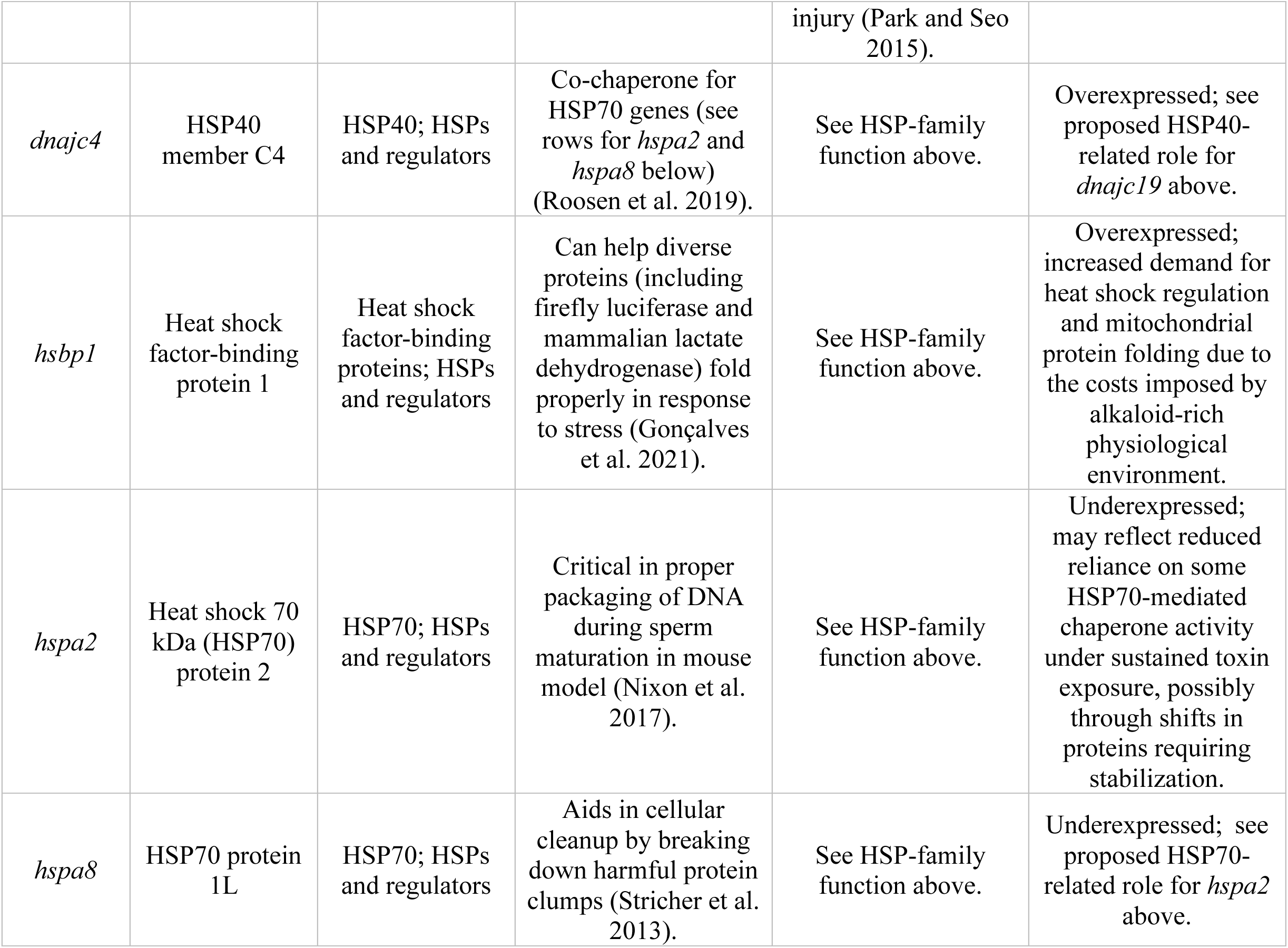
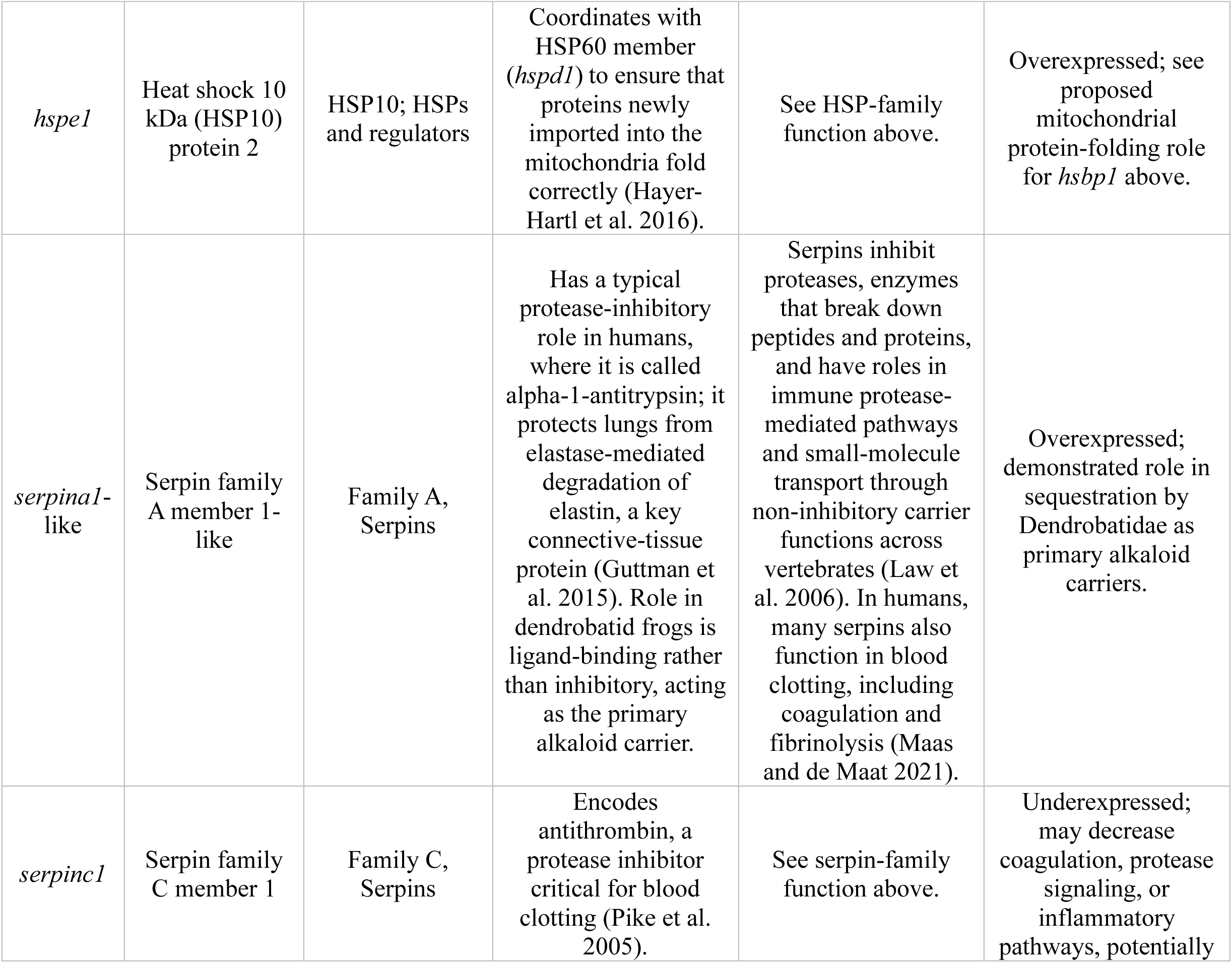
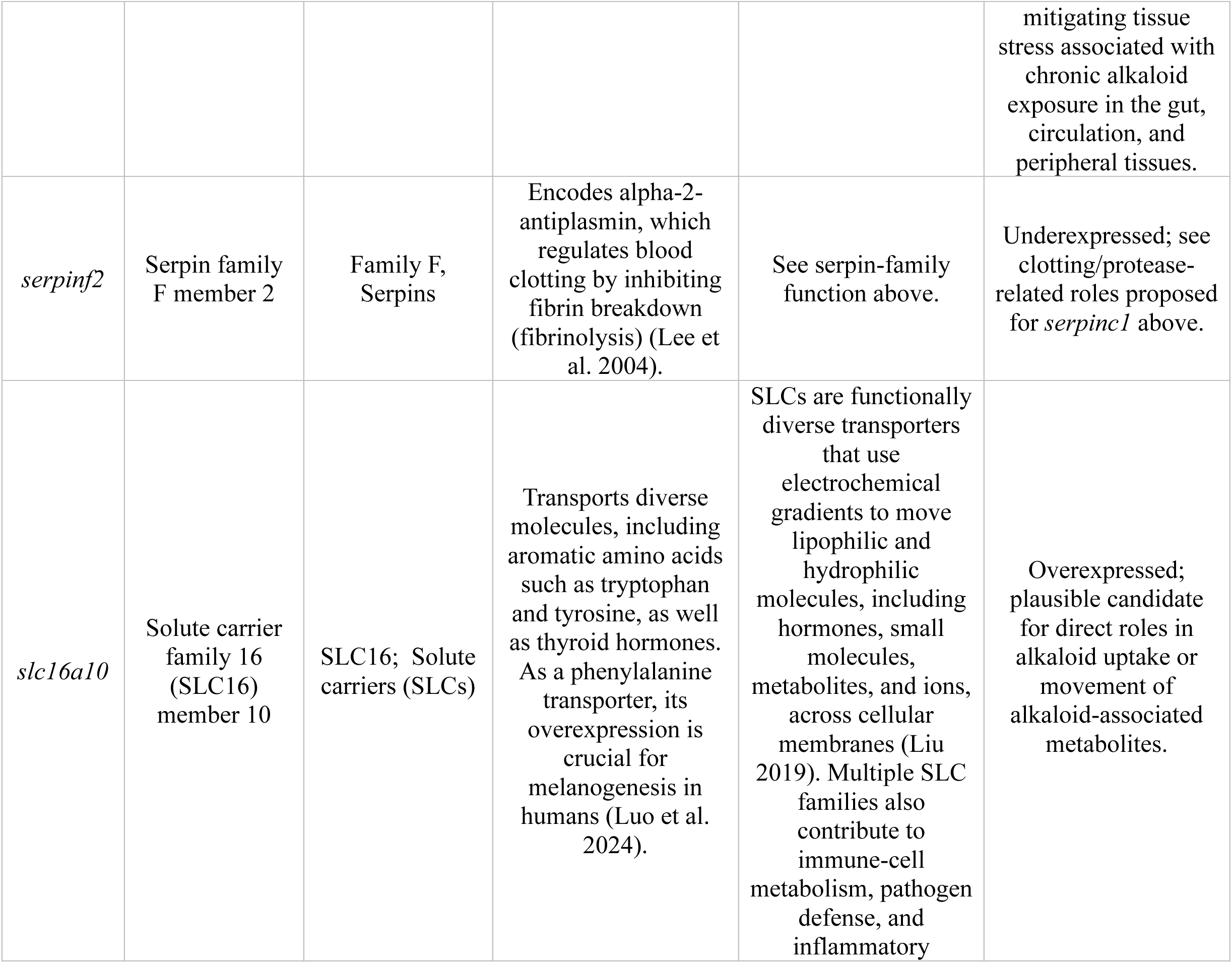
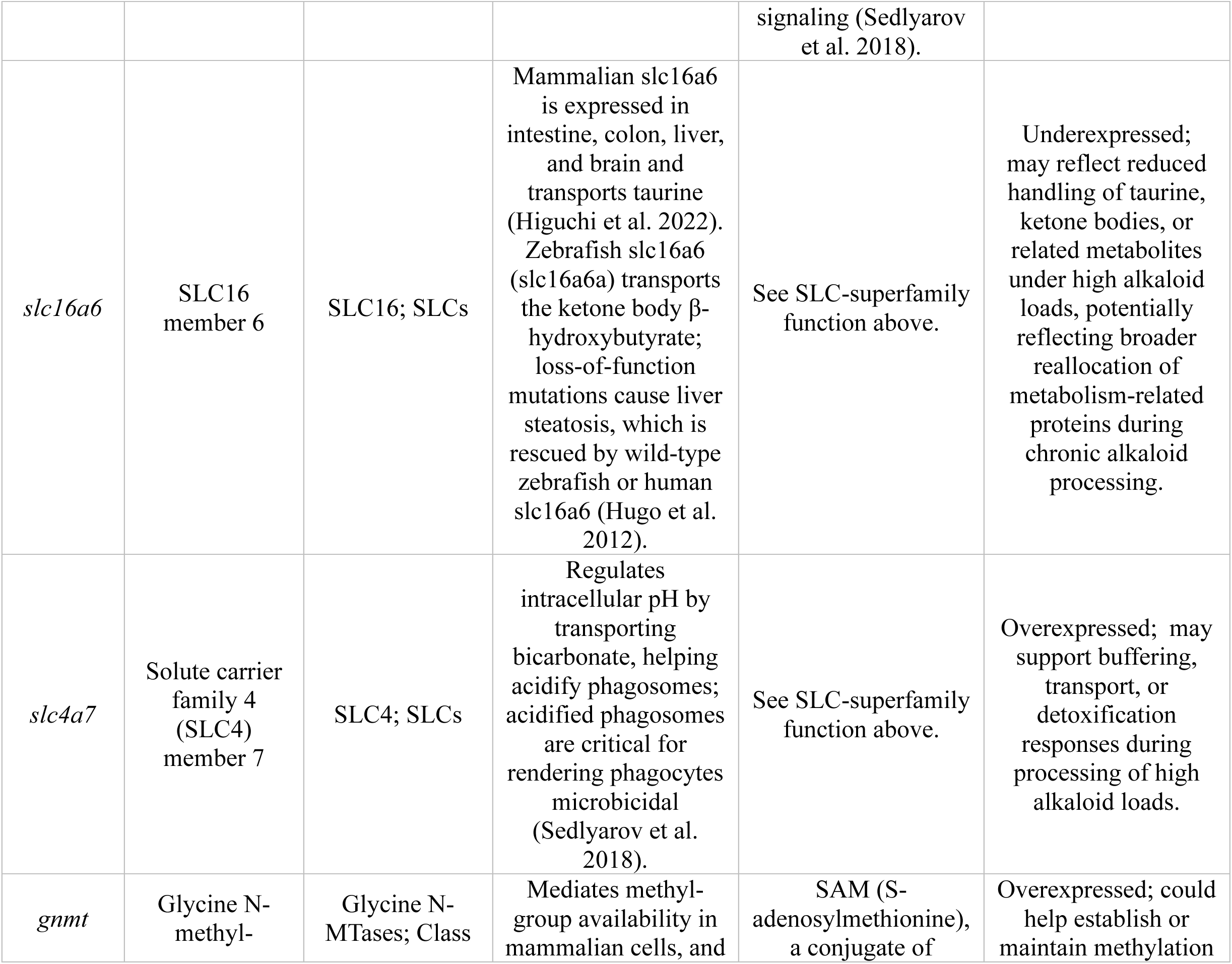
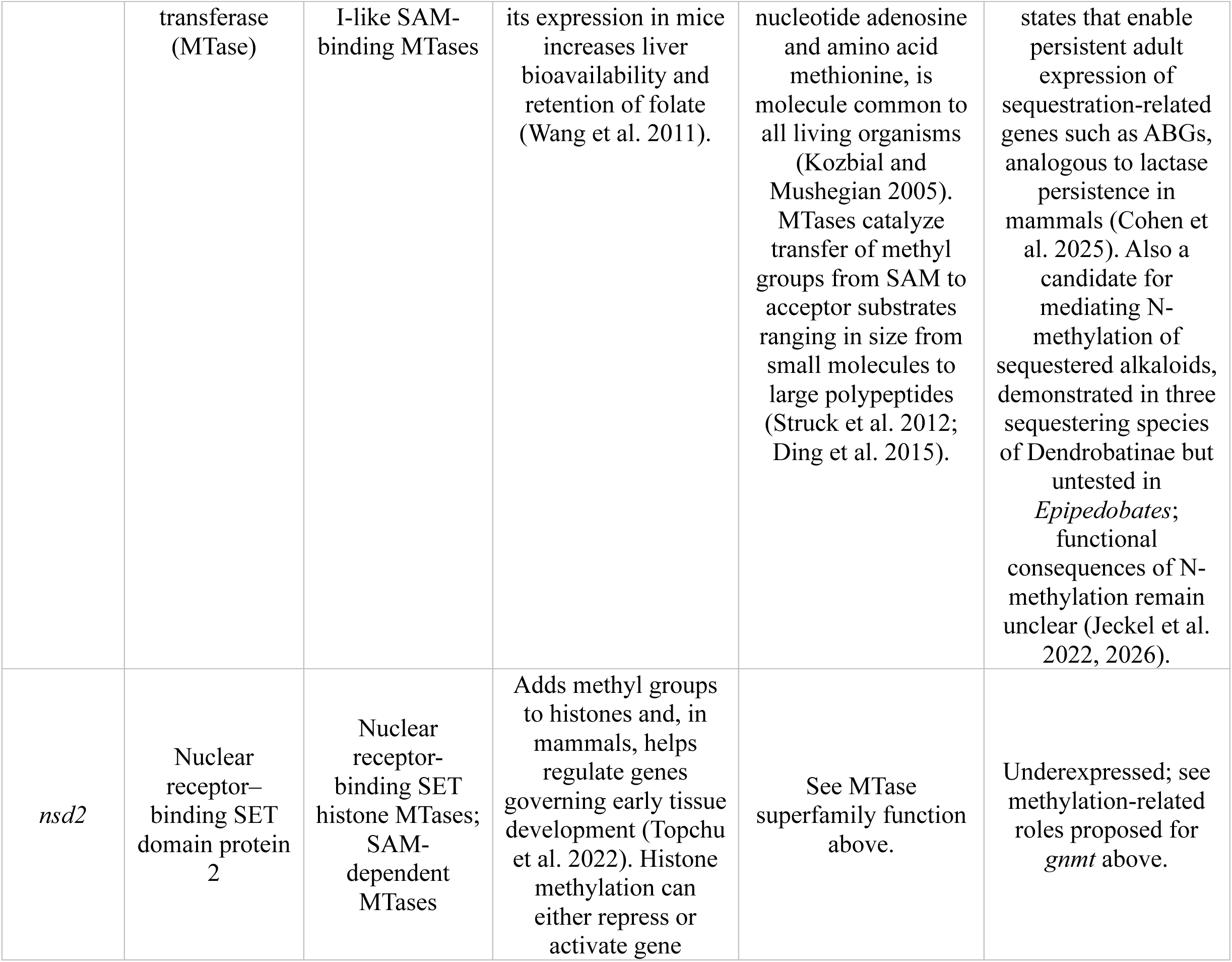
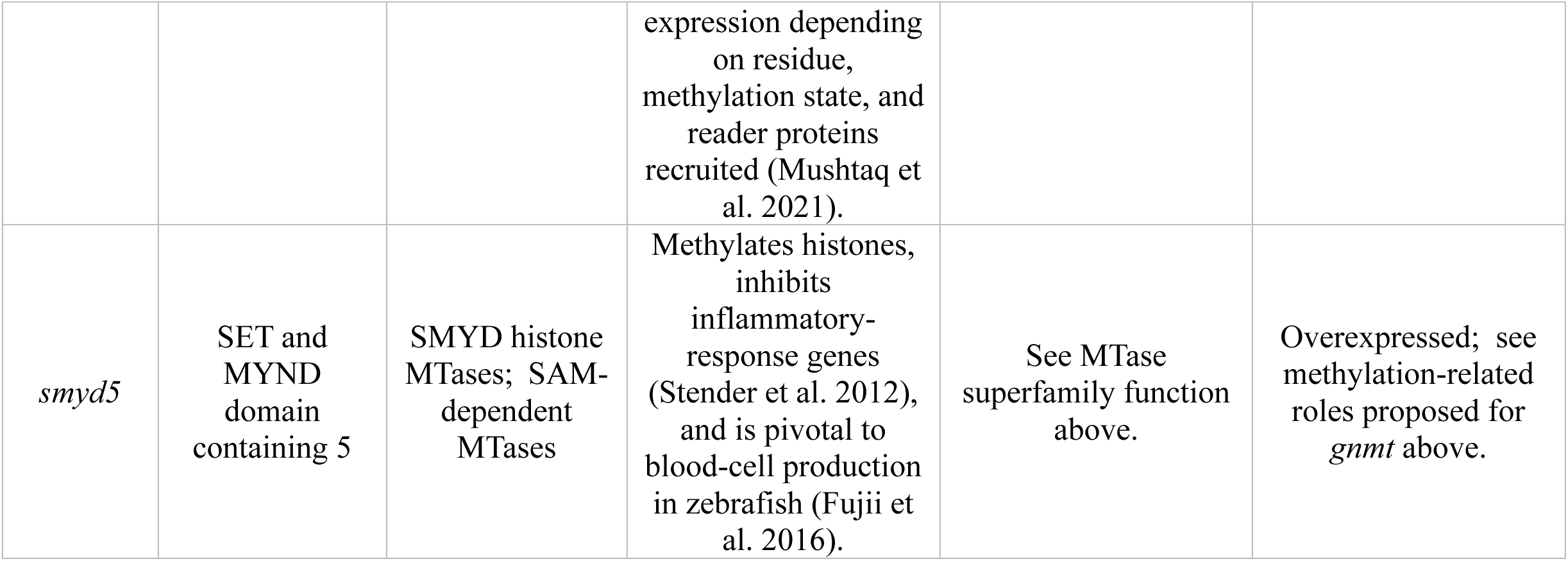
Candidate genes identified among DEGs in our study. For each gene, columns list the encoded protein; gene/protein family and superfamily; main molecular function; expression direction in *Epipedobates*; and hypothesized role in sequestration.

**Table 2.**
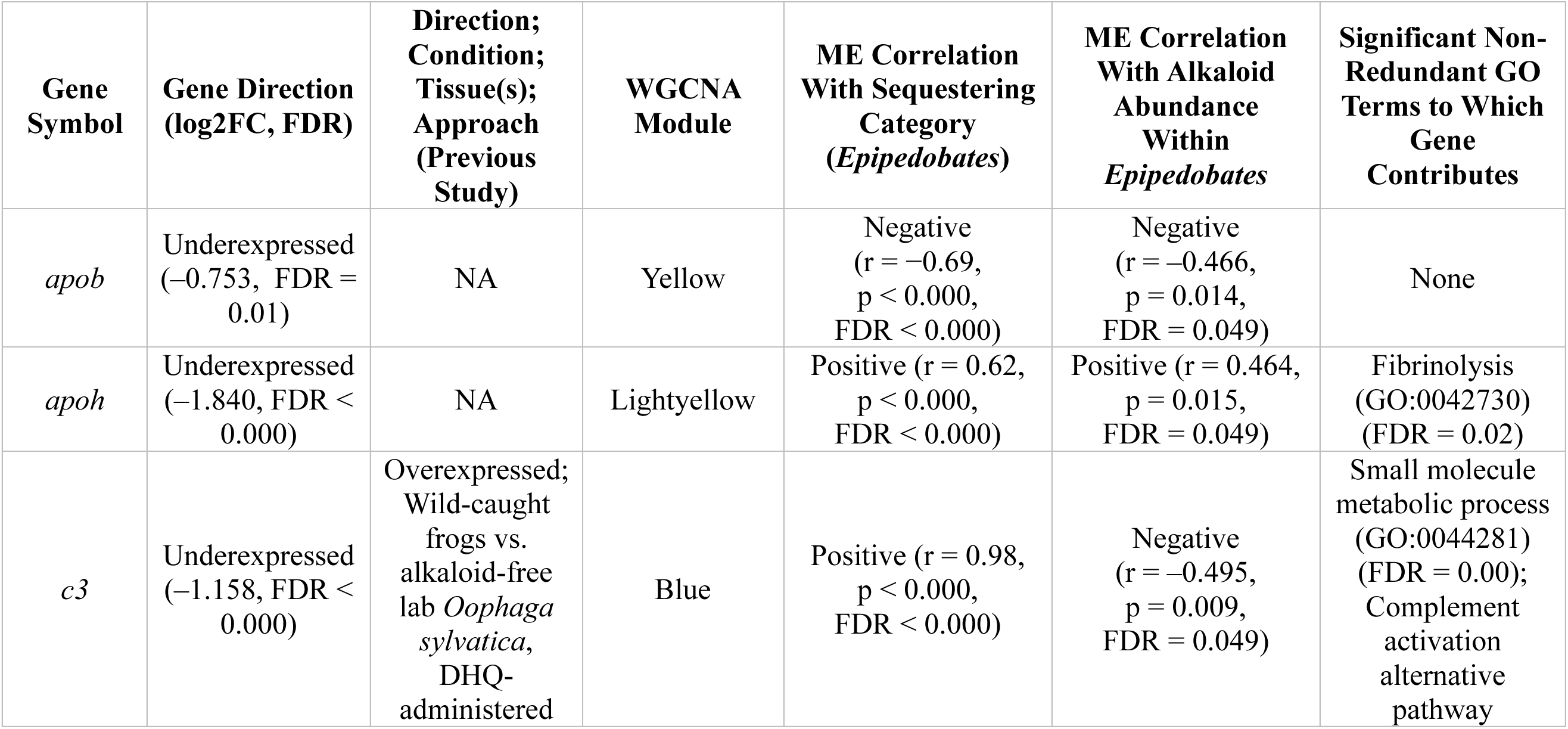
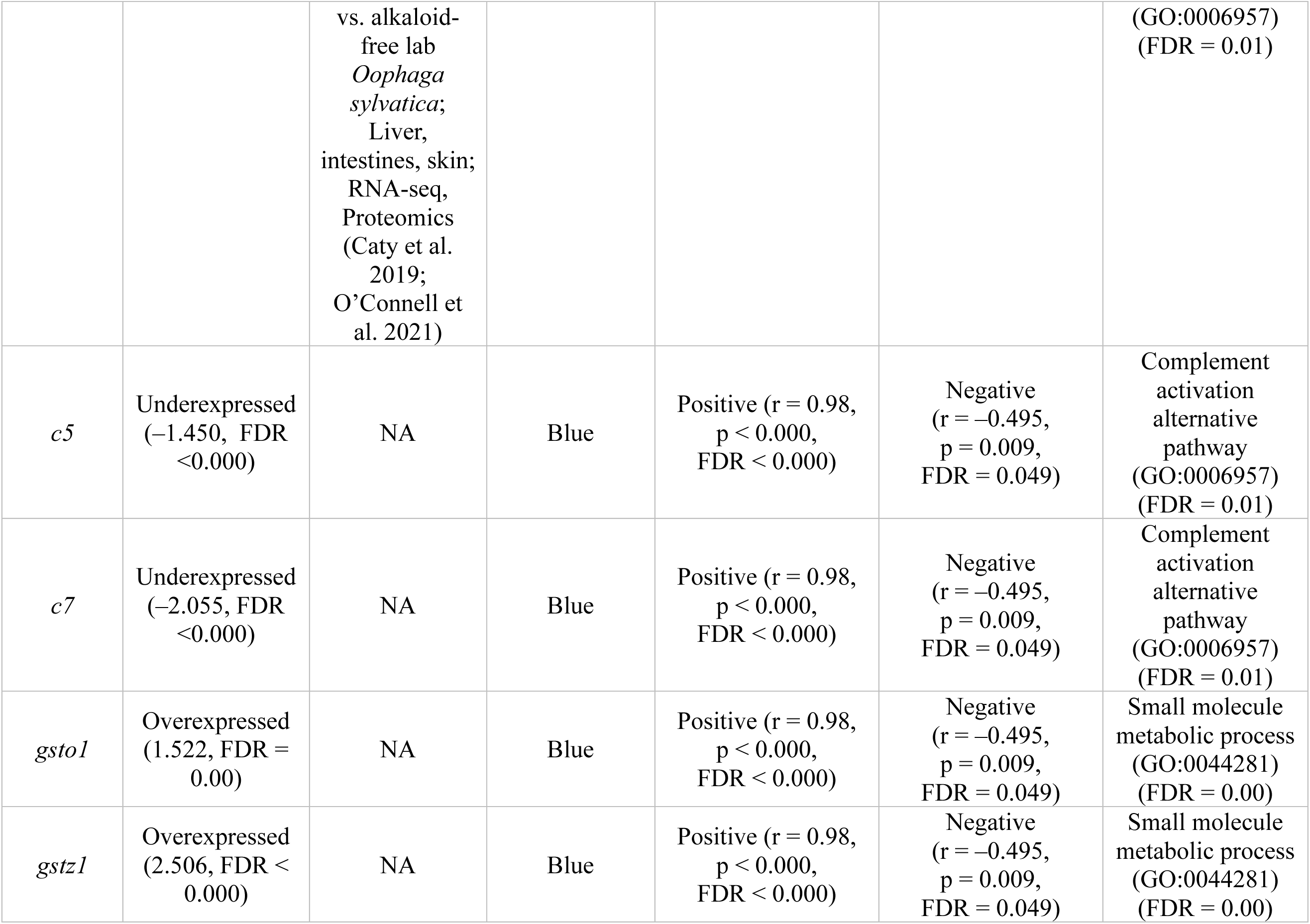
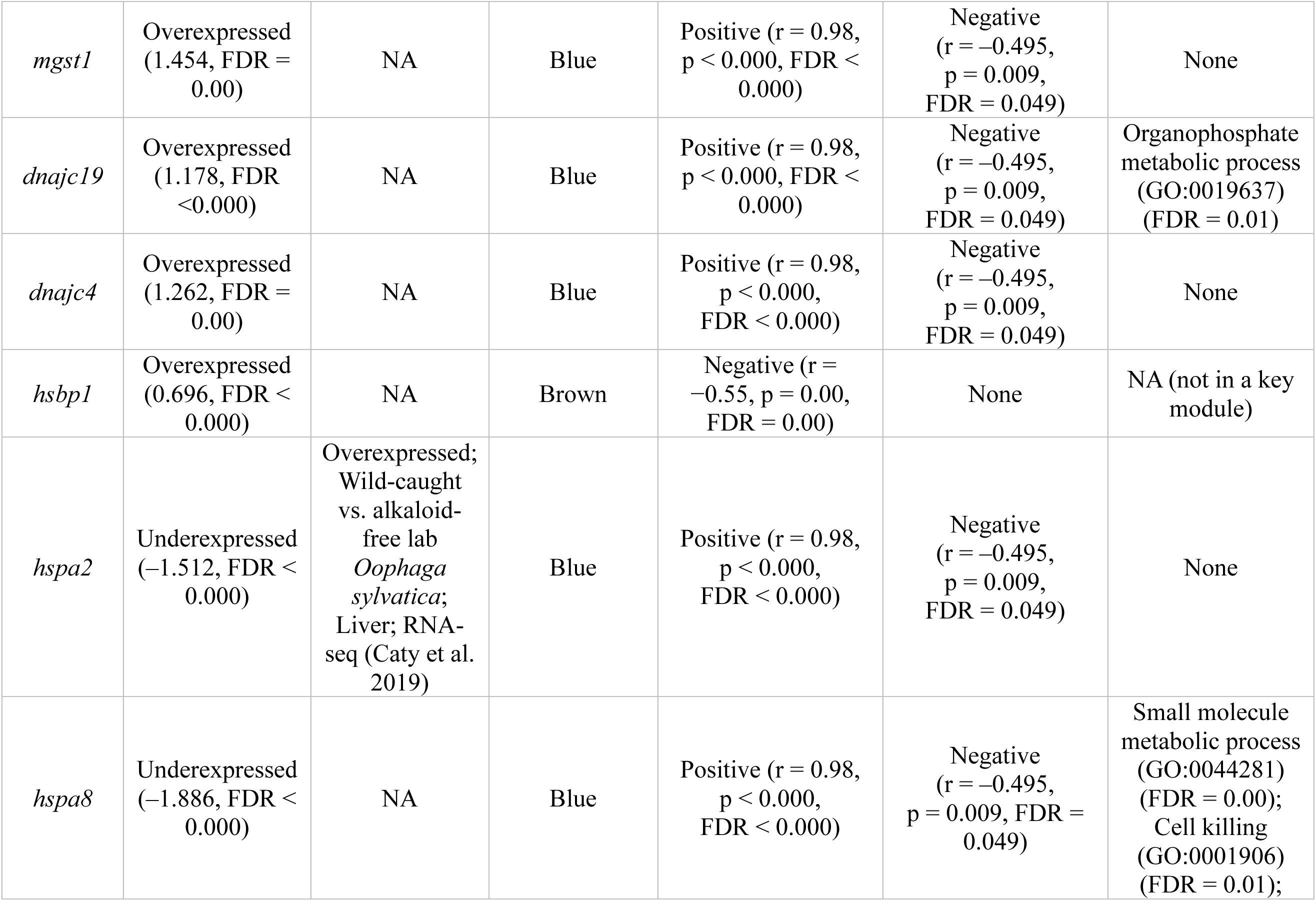
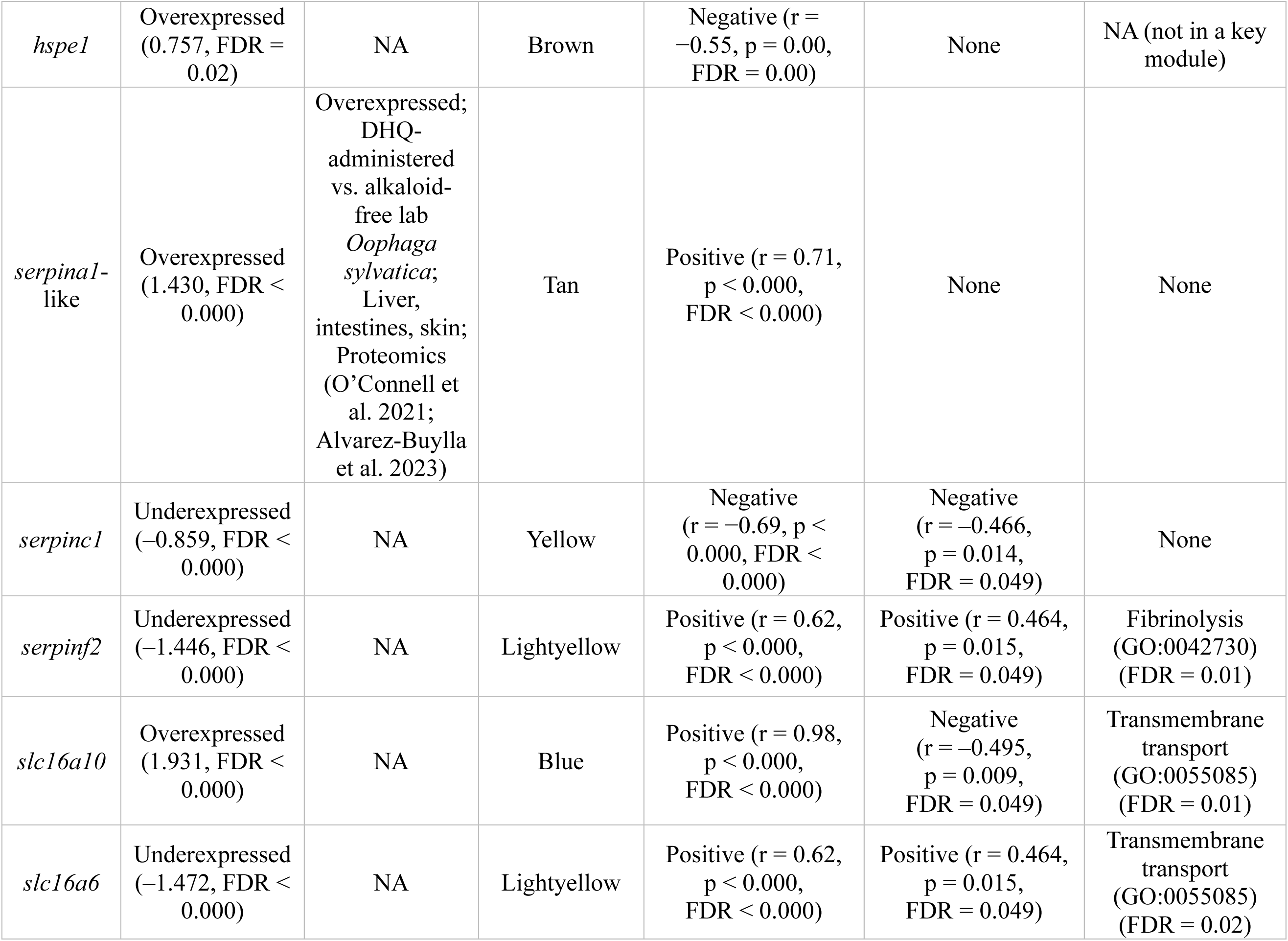
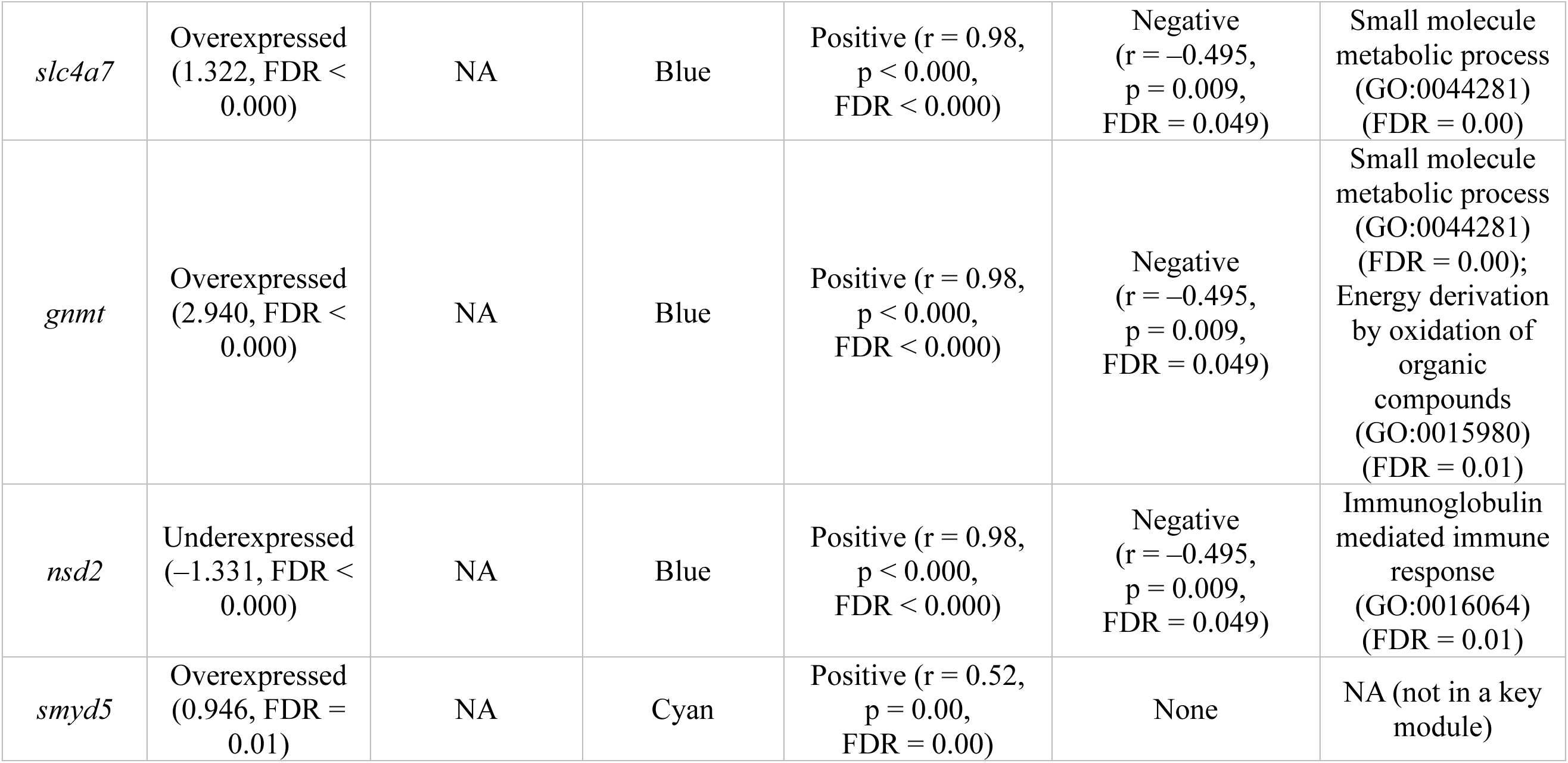
Candidate DEGs, their WGCNA module assignments, expression direction in sequestering category (*Epipedobates*), log2FC values, and BHFDR-corrected p-values. For genes identified in previous dendrobatid liver studies of alkaloid exposure, we report the prior direction of differential expression, contrast condition, additional tissues in which differential expression was detected, and genomic approach. We also report ME correlations with sequestration and marginally significant (FDR = 0.049) associations between ME and alkaloid abundance within *Epipedobates*; because these are module-level statistics, module-eigengene correlation values are repeated for candidate DEGs assigned to the same WGCNA module. For genes in key modules, we list the most significant non-redundant GO terms to which each gene contributes, with BHFDR-corrected enrichment p-values. PTX = pumiliotoxin; DHQ = decahydroquinoline (Caty et al. 2019; O’Connell et al. 2021; Alvarez-Buylla et al. 2023).

### Weighted Gene Co-expression Network Analysis (WGCNA)

WGCNA identifies groups of genes with correlated expression across samples; co-expression likely reflects shared regulation, pathway participation, or related biological functions. To determine functions of broader co-expression neighborhoods for candidate DEGs, we conducted a WGCNA using the 940 filtered and normalized genes retained for the DGE analysis. The analysis recovered 20 co-expression modules. For each module, we calculated a module eigengene (ME), defined as the first principal component of the expression matrix for genes assigned to that module. Thus, each module is represented by an ME profile with one score per sample, where ME scores describe each sample’s position along the dominant shared expression pattern of that module across samples (supplementary fig. S10; supplementary tables S8, S9; Langfelder and Horvath 2008). Modules are named using arbitrary and analysis-specific colors. The MEs of ten modules were significantly associated with the sequestering category (supplementary table S10). These modules contained 379 of the 401 (94.5%) DEGs identified with limma (Ritchie et al. 2015), indicating broad concordance between the DGE and co-expression analyses (supplementary table S9).

For each module, we compared ME scores among species using pairwise Welch two-sample t-tests, with p-values adjusted for multiple comparisons within each module-trait combination using the Benjamini-Hochberg false-discovery-rate (BHFDR) procedure. Pairwise differences with adjusted p ≤ 0.05 were considered statistically significant. MEs from six modules (supplementary tables S9, S10; blue, tan, yellow, brown, cyan, and midnightblue) did not differ between the two sequestering species but differed from one (brown, cyan, midnightblue) or both (blue, tan, yellow) trace-accumulating species (Fig. 6a; supplementary fig. S11; supplementary table S11). Five modules (blue, tan, yellow, lightyellow, and pink) were both significantly enriched for DEGs (FDR ≤ 0.05) and significantly associated with the sequestration category (Table 2; supplementary tables S9, S10). These five modules contained 263 of the 401 DEGs (65.6%): 133 of the 139 genes in the blue module were DEGs (95.7%), 28 of the 38 genes in tan (73.7%), 19 of the 26 genes in yellow (73.1%), 79 of the 153 genes in lightyellow (51.6%), and four of the 21 genes in pink (19.0%) (supplementary table S9).

**Figure 6.**
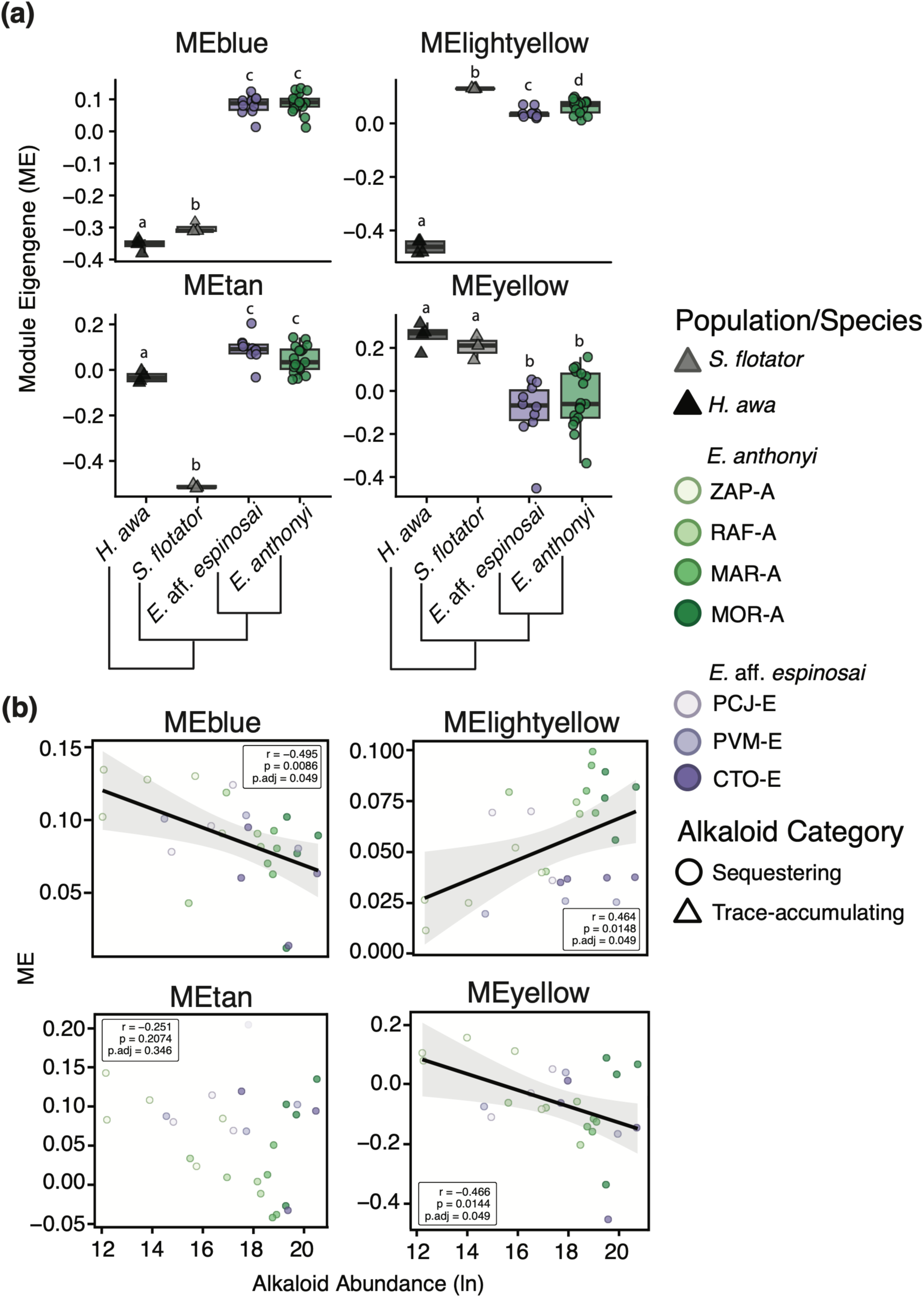
Panel (a); module eigengene values for four WGCNA modules associated with sequestration. Boxplots show module eigengene distributions by species, with individual points representing sampled individuals. Letters indicate significant differences among species based on BHFDR-corrected pairwise Welch two-sample t-tests. Panel (b); scatterplots showing relationships between module eigengene values and ln-transformed alkaloid abundance within *Epipedobates* for the same four WGCNA modules. Pearson correlation coefficients, raw p-values, and BHFDR-adjusted p-values are shown within each panel. Regression lines are ordinary least-squares fits, and shaded bands indicate 95% confidence intervals. The MEtan scatterplot lacks a regression line and confidence interval because the correlation was non-significant. Samples are colored by population, with circles indicating sequestering species and triangles indicating trace-accumulating species. Alt text: Two-panel figure showing WGCNA module associations. Four sequestration-associated modules differ among species and correlate variably with alkaloid abundance in *Epipedobates*.

### Alkaloid Categories and Biological Functions of Key Modules

Twenty of the 23 candidate DEGs were concentrated in four sequestration-associated modules: blue, tan, yellow, and lightyellow (supplementary figs. S12–S15). We therefore treated these as key modules for downstream visualization and interpretation. Fourteen candidate DEGs (*c3*, *c5*, *c7*, *gsto1*, *gstz1*, *mgst1*, *dnajc19*, *dnajc4*, *hspa2*, *hspa8*, *slc16a10*, *slc4a7*, *gnmt*, *nsd2*) were assigned to the blue module; *serpina1*-like was assigned to tan; *apob* and *serpinc1* were assigned to yellow; and *apoh*, *serpinf2*, and *slc16a6* were assigned to lightyellow (Table 2). We visualized these four key modules as co-expression networks, with node color intensity indicating log-fold expression (see Methods; supplementary figs. S12–S15). The remaining three candidate genes were assigned to brown (*hspe1* and *hsbp1*) or cyan (*smyd5*) and were not included among the key-module network visualizations (supplementary table S9).

Of the 10 modules with significant ME-category associations, three key modules (blue, yellow, and lightyellow) also showed FDR-significant correlations with alkaloid abundance among *Epipedobates* individuals, although these associations were close to the significance threshold (Fig. 6b; Table 2; supplementary figs. S12, S14–S16). These modules therefore represent co-expression networks strongly associated with sequestration category, with marginal statistical support for correlations with alkaloid abundance within *Epipedobates*. Gene ontology (GO) enrichment analysis of the full module gene sets recovered significantly overrepresented, non-redundant biological-process terms for the blue and lightyellow modules (Table 2; supplementary figs. S17, S18; supplementary tables S12, S13). Among these enriched terms, we identified those that included candidate DEGs, linking module-level functional enrichment to specific genes of interest (Table 2). No significant GO terms were recovered for the tan or yellow modules. Given the low number of genes in both modules (supplementary table S9), this absence of enriched terms likely reflects limited power.

The MEblue values were positively associated with the sequestering category and showed marginal (near-threshold) FDR-significant support for a negative correlation with alkaloid abundance among *Epipedobates* individuals (Fig. 6a,b). GO terms significantly enriched in the blue module were related to small-molecule transport and metabolism, as well as immune functions, including complement activation and immunoglobulin-mediated immune responses (Table 2; supplementary fig. S17; supplementary table S12). The lightyellow ME values were also positively associated with sequestration and showed near-threshold FDR-significant support for a positive correlation with alkaloid abundance within *Epipedobates* (Fig. 6a,b). Lightyellow was enriched for processes related to blood clot formation and dissolution (Table 2; supplementary fig. S18; supplementary table S13). Notably, MElightyellow values were highest in the trace-accumulating species *S. flotator*, intermediate in the sequestering *Epipedobates*, and lowest in the other trace-accumulating species, *H. awa* (Fig. 6a).

Although the yellow module lacked significant GO enrichment, its two candidate DEGs, *apob* and *serpinc1*, had functionally related genes in the lightyellow module (*apoh* and *serpinf2*, respectively) that contributed to significantly enriched GO terms related to blood clotting (Table 2). MEyellow values were negatively associated with the sequestering category and showed near-threshold FDR-significant support for a negative correlation with alkaloid abundance within *Epipedobates*, in contrast to the positive module-abundance association observed for lightyellow (Fig. 6a,b; Table 2).

Finally, the tan module contained *serpina1*-like (ABG). Given its possible role in alkaloid transport and binding, this module unsurprisingly was positively correlated with sequestration and had higher ME values in sequestering than trace-accumulating species, though the module may not track finer-scale variation in toxin load given that we did not find statistical support for an association with alkaloid abundance among *Epipedobates* individuals (Fig. 6b; Table 2). MEtan values for the two trace-accumulating species differed significantly from each other; values for *H. awa* were higher than those for *S. flotator* (Fig. 6a).

## Discussion

High-throughput approaches are illuminating the molecular basis of chemical defenses in non-model organisms, including amphibians (e.g., Lan et al. 2022; Alvarez-Buylla et al. 2023; Chen et al. 2024). Neotropical poison frogs (Dendrobatidae) provide an exceptional system, having evolved the ability to sequester alkaloids in high diversity and abundance at least three times from a trace-accumulating ancestral state characterized by low alkaloid diversity and abundance (Santos et al. 2014; Tarvin et al. 2024). Mechanistic studies of dendrobatid chemical defense have yielded important insights into physiological responses to alkaloid exposure but have focused largely on single species within Dendrobatinae, which represents the oldest of the three origins of alkaloid sequestration (>25 mya) (e.g., Caty et al. 2019; O’Connell et al. 2021; Alvarez-Buylla et al. 2022, 2023), with comparatively few incorporating trace-accumulating taxa (e.g., Sanchez et al. 2019).

Here, we combined a targeted analysis of gene-family evolution with an untargeted but evolutionarily finer-scale gene-expression comparison to investigate the molecular basis of sequestration in dendrobatids. Given the demonstrated mechanistic importance of ABG proteins, we reconstructed the evolutionary history of the relevant gene family (serpinA) across amphibians, including sequences from both trace-accumulating and sequestering dendrobatids. We then compared liver gene expression between two species of *Epipedobates*, the youngest origin of sequestration in Dendrobatidae (∼12–15 mya), and two trace-accumulating lineages (*Silverstoneia* and *Hyloxalus*), to identify genes and gene networks associated with the sequestering phenotype.

In Dendrobatidae, we identified patterns consistent with extensive duplication and diversification in both *serpina1*-like genes (ABGs) and BBS-like genes, which formed two and five major clades, respectively, each containing multiple tissue- and species-specific gene variants (Figs. 2, 3). *Serpina1*-like was also identified among our candidate DEGs and was overexpressed in *Epipedobates* relative to trace-accumulating lineages.

We next examined other candidate DEGs (aside from *serpina1*-like) that we identified in *Epipedobates*. Because the functional roles of these genes in sequestration remain unresolved, we focused our DGE analyses on curated candidate superfamilies previously implicated in dendrobatid responses to alkaloid exposure, specifically in *O. sylvatica* and *D. tinctorius* (supplementary table S5), or broadly associated with xenobiotic handling in the vertebrate liver (supplementary table S6). Because transcriptomic studies in non-model systems often yield large numbers of candidates with uncertain functions (Conesa et al. 2016), integrating gene-level hypotheses with pathway-level approaches such as co-expression network analysis can strengthen biological inference. Accordingly, we combined DGE, WGCNA, and GO analyses. We evaluated expression in the liver, a major site of xenobiotic metabolism in vertebrates (Trefts et al. 2017) and a key tissue for processing dietary alkaloids in dendrobatids (Caty et al. 2019; Jeckel et al. 2022; Minder et al. 2026).

Candidate DEGs distinguishing sequestering and trace-accumulating categories were involved in small-molecule transport and metabolism, as well as immune responses to xenobiotics (Caty et al. 2019; O’Connell et al. 2021; Alvarez-Buylla et al. 2022, 2023). Interpreting these genes within their co-regulated WGCNA modules allowed us to infer broader biological processes associated with the sequestering phenotype. Our results chiefly implicated pathways related to small-molecule binding and transport, detoxification, and immune function. WGCNA also enabled the identification of genes with potential as exploratory targets in future studies beyond predefined candidate sets. For example, *rab1a* appears in the blue module and *rab10* in the lightyellow module, with both overexpressed in *Epipedobates* (supplementary figs. S12, S15). These genes encode Rab GTPases, key regulators of intracellular membrane trafficking and vesicle transport (Hutagalung and Novick 2011), suggesting that vesicle-mediated trafficking may warrant further investigation as a potential component of sequestration.

We also examined whether candidate genes were correlated with alkaloid abundance within *Epipedobates*. We found no significant gene-level correlations. However, three of the ten phenotype-associated co-expression modules, all belonging to the subset of four modules enriched for DEGs, showed marginal support for correlations with *Epipedobates* alkaloid abundance (Table 2).

The serpinA phylogeny and DGE analyses together suggest that diversification of alkaloid-binding proteins and regulatory evolution in pathways involved in alkaloid transport, metabolism, and resistance have enabled the evolution of abundant and structurally diverse diet-acquired toxins in poison frogs. However, the binding activities of ABG and BBS-like paralogs and variants (besides OsABG1, DtABG1, EtABG1) and the roles of the candidate DEGs require functional validation. Our results therefore supply research directions for future work and provide a framework for exploratory studies of the molecular mechanisms underlying dietary toxin acquisition in diverse non-model taxa.

### SerpinA Genes

The serpin superfamily is the largest and most ubiquitous group of protease inhibitors (Law et al. 2006; Seixas 2015). Although many vertebrate serpins function in this way, others have diversified into non-inhibitory roles, including hormone transport (Pemberton et al. 1988), molecular chaperoning (Nagata 1996), and tumor suppression (Zou et al. 1994). In mammals, serpins have also been implicated in toxin resistance through inhibition of serine proteases common in venoms (Gibbs et al. 2020; Cavalcante et al. 2022; Ward et al. 2025). In frogs, two serpinA1 lineages relevant to small-molecule binding and transport—biliverdin-binding serpins (BBSs) and alkaloid-binding globulins (ABGs)—appear more similar functionally to non-inhibitory transporter genes from the serpinA family, such as the canonical protein C inhibitor (*serpina5*), corticosteroid-binding globulin (*serpina6*), and thyroxine-binding globulin (*serpina7*), than to inhibitory serpins like the canonical lung- and liver-expressed *serpina1* gene (A1AT) in humans, which prevents the breakdown of connective protein elastin by the destructive enzyme elastase (Seixas 2015). Our phylogeny indicates that the serpinA1 subfamily diversified within dendrobatids. Consistently, dendrobatid ABGs show pronounced biochemical divergence at putative binding-site residues relative to other frogs, including substitutions predicted to alter polarity near the ligand-binding pocket (Fig. 2). Although the forces driving the initial divergence of BBS-like and *serpina1*-like at the base of modern frogs remain unclear, the expansion of these clades in dendrobatids may reflect repeated co-option of SerpinA proteins for the safe handling of diet-derived alkaloids.

Dendrobatid *serpina1*-like proteins (ABGs) appear to have evolved a novel function as transporters of dietary alkaloids. A proteomic study by O’Connell et al. (2021) showed that an ABG from *O. sylvatica*, then identified as the ortholog of the canonical human A1AT, was significantly more abundant in the livers of decahydroquinoline-fed frogs than in controls. Subsequent work by Alvarez-Buylla et al. (2023) identified this protein, OsABG1, as the highest-affinity plasma protein for a PTX-like photoprobe in *O. sylvatica*. Recombinant expression of putative orthologs from *D. tinctorius* (DtABG1) and *E. tricolor* (EtABG2) revealed variable binding among alkaloids from three classes: high-affinity binding by both to PTX **251D**, moderate binding by both to decahydroquinoline, and weak (DtABG1) or no (EtABG2) binding to epibatidine. In contrast, assays with OsABG1 demonstrated that it bound all three alkaloids and two additional ligands (a histrionicotoxin-like compound and indolizidine ring) with high affinity. However, the PTX-photoprobe-binding assay indicated that the primary alkaloid-binding proteins in *D. tinctorius* and *E. tricolor* are slightly larger than OsABG1, although those proteins were not identified at the time (genomes were unavailable for quantitative proteomics) (Alvarez-Buylla et al. 2023). This raised the possibility that distinct paralogs contribute to alkaloid transport or sequestration in different dendrobatid lineages.

Our serpinA reconstruction helps reinterpret these findings. Our results highlight the difficulty of assigning homology within ABGs, a rapidly diversified gene group, and suggest that the identity and binding properties of major alkaloid-binding proteins may differ among species and tissues. Although the binding results from Alvarez-Buylla et al. (2023) show that a single ABG can bind structurally diverse alkaloids, additional paralogs or variants may show higher affinity for specific alkaloid classes. For example, DtABG1 may not be the highest-affinity binding variant for epibatidine, which *D. tinctorius* sequesters efficiently (Waters et al. 2024). We recovered two major dendrobatid ABG clades, ABG1 and ABG2 (Figs. 2, 3; supplementary figs. S1, S2). The principal ABG identified in *O. sylvatica* (OsABG1) and its putative homolog in *D. tinctorius* (DtABG1) both belong to ABG1 and cluster with sequences present in liver and intestine transcriptomes, but not the skin transcriptome, of *O. sylvatica*. This pattern is consistent with that uncovered by Alvarez-Buylla et al. (2023), who reported highest expression of OsABG in the liver, moderate-to-low expression in the intestine, and no expression in the skin. In contrast, skin-expressed *serpina1*-like variants of *O. sylvatica* are nested within ABG2, together with skin-expressed variants from other sequestering dendrobatids, including the previously proposed *E. tricolor* homolog of OsABG1, which we interpret here as the paralog EtABG2.

Alvarez-Buylla et al. (2023) reported little or no plasma binding activity by the PTX-**251D**-like photoprobe in both *A. femoralis* and *Mantella aurantiaca* (Mantellidae), the latter representing a separate origin of alkaloid sequestration in frogs. Assuming plasma binding is a critical and conserved aspect of ABG-mediated alkaloid transport, this result is broadly consistent with the apparent absence or reduced diversity of *serpina1*-like variants in these taxa (supplementary figs. S1, S2). Although beyond the main scope of this study, *M. aurantiaca* also appeared to harbor relatively high diversity of *serpina6*-like genes, which encode corticosteroid-binding globulins, consistent with the proposal that mantellids may rely on a partially distinct sequestration mechanism involving a different ligand-transport serpin clade (Meyer et al. 2016; Alvarez-Buylla et al. 2023). In contrast, in *Pseudophryne corroborree*, which represents another anuran origin of alkaloid sequestration, we recovered only two *serpina1*-like variants and single variants of other serpinA genes: BBS, *serpina7*-like and *serpina10*-like. This pattern raises the possibility that sequestration in *Pseudophryne* may rely on mechanisms largely independent of serpinA diversification.

Members of the other highly diversified serpinA1 group in dendrobatids, BBS-like, are involved in physiological chlorosis in diverse arboreal frogs; chlorosis has evolved at least 41 times in 11 anuran families (Taboada et al. 2020). BBSs have been functionally studied only in hylids and centrolenids. Whereas most vertebrates rapidly excrete biliverdin, a toxic waste product of heme catabolism, BBSs enable chlorotic lineages to safely sequester biliverdin and alter its spectral properties through protein binding, rendering bones, lymph, and other tissues blue-green (Taboada et al. 2020). In most mammals, biliverdin is rapidly reduced to bilirubin before excretion; chlorotic frogs, however, reportedly tolerate biliverdin concentrations at least fourfold higher than those observed in severe human hyperbiliverdinemia and roughly 200-fold higher than those in non-chlorotic frog species (Greenberg et al. 1971; McDonagh 2001; Taboada et al. 2020).

BBS-like is expanded in Dendrobatidae and is more diverse than BBSs in chlorotic frogs, represented in our analysis by all sampled centrolenids (*Chimerella*, *Espadarana*, and *Nymphargus*) and several hylids (*Sphaenorhynchus*, *Pseudis*, *Boana*, *Hyloscirtus*, and *Aplastodiscus*; Figs. 2, 3). This result is surprising, because dendrobatids are not known to exhibit unusually high biliverdin concentrations among anurans (Taboada et al. 2020; Aguilar-Gómez et al. 2026) or to include species with blue-green bones (see Taboada et al. 2020 supplementary materials). We identified five dendrobatid BBS-like clades (A–E), each with species- and tissue-specific variants. Consistently, Taboada et al. (2020) found evidence for the occurrence of species- and tissue-specific BBS paralogs in chlorotic frogs.

The functional significance of BBS-like clades in dendrobatids is unclear. Some variants may retain biliverdin-binding roles. One possibility is that a restricted subset of BBS-like paralogs contributes to carotenoid uptake, trafficking, or deposition in certain brightly colored dendrobatids, since vertebrates must obtain their carotenoids, which often underlie yellow, orange, and red coloration, from the diet (Toomey et al. 2022). Although the genetic basis of carotenoid sequestration in dendrobatids remains poorly understood outside of *Oophaga pumilio* and *O. vicentei* (Aguilar-Gómez et al. 2026; Mantzana-Oikonomaki et al. 2026), BBS-like genes represent plausible lineage-specific candidates. Nonetheless, hypotheses centered on external coloration do not readily explain why the trace-accumulating, inconspicuously-colored *Anomaloglossus baeobatrachus* possesses paralogs from each BBS-like clade, including four variants of the BBS-like B paralog (supplementary figs. S1, S2). An alternative is that dendrobatid BBS-like proteins contribute to physiological tolerance of dietary alkaloids, analogous to the role of BBSs in permitting exceptionally high circulating biliverdin levels in chlorotic frogs. Both sequestering and trace-accumulating dendrobatids consume ants and mites, prey types associated with alkaloids, at moderate to high frequencies (Saporito et al. 2004, 2007b; Tarvin et al. 2024). Across dendrobatids, BBS-like proteins may therefore bind excess alkaloids or their metabolic byproducts and facilitate their clearance.

Some members of dendrobatid BBS-like clades may also retain other ancestral serpin functions. For example, hylaserpin-S1, which is nested within the hyloid biliverdin-binding serpin group (supplementary figs. S1, S2), inhibits trypsin and chymotrypsin and exhibits antimicrobial activity (Wu et al. 2011). If BBS-like proteins across neobatrachians, including dendrobatids, are found to possess antimicrobial activity or broad metabolite-binding properties, then biliverdin transport might represent one specialized manifestation of a more general multifunctional role. BBS-like could be involved variably in transport or sequestration of metabolites as well as in innate defense.

### The Rapid Diversification of ABGs and BBS-like Genes

Examples from mammals suggest rapid elaboration and functional diversification of serpinA genes may represent a recurrent route by which vertebrates evolve to deal with toxins. Within and among rodent lineages, *serpina1* and *serpina3* have undergone repeated expansion and show signatures of diversifying selection (e.g., Barbour et al. 2002; Ward et al. 2025). In ground squirrels (*Otospermophilus beecheyi*), accelerated evolution of a *serpina1* paralog is associated with resistance to snake venom serine proteases (Gibbs et al. 2020; Ochoa et al. 2023). Likewise, among the 12 *serpina3* paralogs in the big-eared woodrat (*Neotoma macrotis*), two inhibit venom serine proteases; one of these has neofunctionalized to inhibit both chymotrypsin-like and trypsin-like proteases (Ward et al. 2025).

Similar patterns of rapid duplication and diversification are common among genes involved in ligand binding and defense. For example, immunoglobulin genes generate antigen-binding diversity through somatic hypermutation in B cells (Yaari et al. 2013; Hoehn and Kleinstein 2024), while amphibian antimicrobial peptide precursors retain conserved signal regions but rapidly evolve mature peptide domains that differ in sequence and bioactivity (Vanhoye et al. 2003). The expansion of dendrobatid BBS-like and ABG sequences into multiple clades (with diverse variants among ABGs) may reflect a broader pattern in which conserved protein scaffolds are repeatedly modified to accommodate new ligand-binding functions. Considering dendrobatid ABGs, an ancestral serpinA1 may have been co-opted for alkaloid exposure and sequestration and subsequently diversified under selection imposed by geographically variable arthropod prey communities (e.g., Saporito et al. 2007c; Martin et al. 2025). Such selection could favor binding pockets that are broadly permissive but chemically tuned to ecologically relevant alkaloid classes (Alvarez-Buylla et al. 2023). Consistent with this possibility, ABG1 and ABG2 differ in polarity at homologous positions corresponding to predicted OsABG1 alkaloid-binding residues, suggesting that ABG paralogs may differ in ligand affinity or specificity.

Given the broad structural diversity of dietary alkaloids sequestered by poison frogs, some degree of ligand-binding promiscuity may be advantageous; evolving a unique binding protein for every alkaloid would likely be inefficient. At the same time, partial specialization among paralogs and variants could improve binding affinity for particular alkaloid classes, potentially explaining the polarity-influencing amino-acid variation observed at putative ABG1 and ABG2 binding sites. In addition, increased numbers of paralogs and variants could increase protein dosage, levels of bound alkaloids, and ultimately skin alkaloid concentrations. Consistent with a possible role for ABG diversification in sequestration, we found in *Epipedobates* and other sequestering dendrobatids sampled in our serpinA analysis (*Ranitomeya imitator*, *R. variabilis*, *Phyllobates terribilis*, *D. tinctorius*, and *O. sylvatica*) multiple ABG1 and/or ABG2 variants; on the other hand, we found in the two basal trace-accumulating dendrobatids we sampled reduced or undetected ABG diversity. We recovered in *Anomaloglossus baeobatrachus* only two ABG variants, both assigned to ABG1, whereas no ABG variants were detected in *Allobates femoralis*. The two ABG1 variants in *An. baeobatrachus* were invariant at the six putative amino-acid binding sites (Figs. 1a, 2, 3; supplementary figs. S1, S2; supplementary table S1). Although our serpin reconstruction included only these two representatives from an early-diverging trace-accumulating dendrobatid lineage (Aromobatinae), these findings raise the possibility that transitions to alkaloid sequestration in Dendrobatidae are associated with ABG sequence diversification and gene duplication. However, reliably assessing whether ABG variant number covaries with alkaloid load, either between sequestering and trace-accumulating lineages or among sequestering species, will require paired chemical and genomic data from diverse dendrobatids, including trace-accumulating species from more derived dendrobatid lineages, such as the two sampled in our gene-expression analysis (*S. flotator* and *H. awa*).

Rather than relying on a single generalized ABG to bind and transport all dietary alkaloids, dendrobatids may possess a suite of ABG-like paralogs and variants with complementary or partially overlapping binding preferences. Under this model, ABG diversity could generate a gradient of alkaloid affinities, facilitating the efficient sequestration of chemically diverse dietary alkaloids. The occurrence of tissue-specific ABG variants further raises the possibility that different paralogs act at distinct stages of the uptake pathway in sequestering frogs, including intestinal uptake, mobilization through circulation, and eventual deposition in the skin (Chen et al. 2025b).

Complementary proteomic and biochemical approaches across structurally distinct alkaloid classes will be needed to test whether ABG1 and ABG2 variants from sequestering and trace-accumulating species differ predictably in binding affinity or specificity. These approaches could also address broader questions about the phylogenetic distribution of alkaloid classes across Dendrobatidae; for example, whether *Phyllobates* possess ABGs specialized for batrachotoxins, highly toxic steroidal alkaloids known only from this genus among dendrobatids (Daly et al. 1987). One strategy would be to use plasma-binding assays with photoprobes that mimic distinct alkaloid classes, combined with gel-punch pulldown proteomics, as in the approach used to identify OsABG in *O. sylvatica* (Alvarez-Buylla et al. 2023). This could be paired with *in vitro* binding assays following recombinant expression and purification of full-length proteins from synthesized or cloned ABG coding sequences. Together, these assays could test whether trace-accumulating dendrobatids also use ABGs, perhaps with reduced variant number, expression, or binding affinity, as part of an incipient or low-efficiency ABG-mediated transport mechanism.

These approaches can also be used to test the alkaloid-binding activity of the expanded repertoires of serpinA1 genes observed in several non-dendrobatid amphibians. In *Hyperolius riggenbachi* we found four *serpina1*-like variants and six BBS-like variants; skin alkaloids were not detected in a study of four *Hyperolius* species (not including *H. riggenbachi*; Portik et al. 2015). Synthesizing and testing the affinity of these proteins for alkaloids could reveal whether low-level binding capacity is more phylogenetically widespread than currently recognized (supplementary figs. S1, S2). We also recovered six A1AT-like variants from five distinct phylogenetic lineages in the salamander *Lissotriton helveticus* (Figs. 2, 3). In the only chemical survey of this species, nine of ten sampled individuals contained the water-soluble alkaloid tetrodotoxin, often at high concentrations (Yotsu-Yamashita et al. 2007). In addition, reports of the highly toxic lipophilic alkaloid samandarone are widespread among profiled Salamandridae, including *Lissotriton boscai* (Vences et al. 2014). Although samandarone is suspected to be synthesized endogenously, tetrodotoxin in salamanders has been linked to dietary mites (Nakazawa et al. 2026) and symbiotic microbes (Vaelli et al. 2020). A1AT-like expansion in salamanders should be evaluated to determine if it is associated with binding, transport, or tolerance of tetrodotoxin.

### ABG Expression in the Most Recent Origin of Sequestration in Poison Frogs

Daly (1998) hypothesized that the high alkaloid levels observed in sequestering dendrobatids may have evolved through the “overexpression of a primitive alkaloid uptake system” present in the trace-accumulating ancestors, although no specific mechanism was proposed. The recent identification of alkaloid-binding activity by ABG proteins provided a plausible mechanism. Findings from our DGE analysis are consistent with this framework: *serpina1*-like was among the candidate DEGs, and both the gene and its associated co-expression module (tan) showed significantly higher expression in *Epipedobates* relative to trace-accumulating species. These findings suggest that overexpression of an ancestral transport mechanism could underlie sequestration, though sequencing data from more *Epipedobates* species are needed to assess whether comparatively high *serpina1*-like expression was the ancestral state for the genus.

Within *Epipedobates*, we did not detect a significant association between *serpina1*-like expression, or tan ME values, and alkaloid abundance (Fig. 6b; supplementary figs. S9, S16). Although limited sample size may have reduced statistical power, it is also possible that ABGs are constitutively expressed once sequestration is established, with among-individual variation in alkaloid load reflecting short-term responses to recent alkaloid intake or subtler effects of longer-term exposure. Notably, *serpina1*-like was the only candidate among the 28 DEGs in the tan module. This suggests that gene networks more directly involved in alkaloid handling may be partly distinct from broader transcriptomic changes associated with the sequestering phenotype. The tan module therefore represents a promising target for identifying regulators of ABG expression and other genes involved in intracellular transport or alkaloid trafficking (supplementary table S9). Last, although both *H. awa* and *S. flotator* accumulate low alkaloid loads relative to *Epipedobates*, the higher MEtan values we observed in *H. awa* could reflect higher expression of a serpin-associated co-expression pathway in this species compared to *S. flotator* (Fig. 6a). The expression of this pathway, while intermediate between that of *Epipedobates* and *S. flotator*, may still be expressed inadequately to result in efficient alkaloid sequestration, especially in tandem with a broader trace-accumulating genomic architecture (a lack of coordinated expression of other key modules, and possibly no expansion and/or poor alkaloid-binding of the ABG repertoire).

Tag-Seq does not reliably distinguish closely related isoforms or paralogs, so our mapping strategy collapsed ABG1 transcripts (no ABG2 transcripts were present in the *O. sylvatica* reference) under the shared annotation *serpina1*-like (see Methods). We thus have not determined in this study whether the higher *serpina1*-like counts observed in *Epipedobates* compared with the trace-accumulating species reflect (1) higher expression of particular broad- or high-affinity paralogs and variants; (2) similar per-paralog and per-variant expression but higher aggregate expression due to a greater number of ABG paralogs and/or variants; or (3) both higher expression and greater breadth of expression across ABG paralogs and variants. Disentangling ABG variant-number diversity from paralog-specific expression will require transcriptomic and proteomic approaches with finer isoform resolution across sequestering and trace-accumulating dendrobatids, including representatives of all three origins of sequestration.

Nonetheless, the broader differential-expression patterns recovered here, including numerous candidate DEGs and multiple co-expression modules associated with the sequestering phenotype, indicate that gene-expression evolution is important for sequestration. Considering the serpinA phylogenetic results, we suggest that ABG diversification, shifts in total expression dosage, and possibly the evolution of higher-affinity binding sites together contributed to the high alkaloid loads observed in sequestering dendrobatids.

### Gene Expression in Epipedobates

Although our phylogenetic analyses focused on serpins, alkaloid sequestration is expected to require coordinated changes across proteins involved in alkaloid metabolism and detoxification (Sanchez et al. 2019; Alvarez-Buylla et al. 2022) and resistance (Tarvin et al. 2016, 2017a; Yeager et al. 2024). We therefore complemented our targeted analysis of serpinA evolution with an exploratory gene-expression study aimed at identifying broader genes and pathways associated with sequestration. This analysis also implicated *serpina1*-like, as discussed above, while revealing expression patterns extending beyond the serpinA gene family.

Comparative transcriptomic studies increasingly enable the identification of both individual genes and co-expressed gene networks associated with complex physiological adaptations, providing insight into whether repeated trait evolution proceeds through shared or lineage-specific mechanisms. Changes in gene expression frequently underlie the rapid evolution of traits under selection (Gilad et al. 2006; Emerson and Li 2010). For example, expression of both distinct and shared kidney-function genes were found to underlie three independent origins of desert specialization in rodents (Bittner et al. 2022). In response to episodes of drought, two populations of field mustard (*Brassica rapa*) evolved changes in the expression of predominantly distinct genes, though overlapped in broader biological processes; expressed genes in both cases were related to stress response and flowering (Hamann et al. 2021).

Alkaloid sequestration in poison frogs likely involves a combination of shared physiological pathways and lineage-specific molecular solutions. Its evolution likely required substantial physiological and metabolic reorganization, including shifts in alkaloid clearance, transport, storage, and tolerance of high toxin burdens. Such changes likely depended on coordinated re-tuning of gene expression. We would expect genes directly involved in uptake, trafficking, and storage of alkaloids to be overexpressed in sequestering clades. In contrast, shifts in detoxification, stress-response, and immune pathways may be less predictable. Stress-response and immune pathways could facilitate tolerance to alkaloid exposure but impose cellular and inflammatory costs and interfere with alkaloid retention when chronically activated.

Detoxification pathways may evolve underexpression in sequestering clades to enhance alkaloid retention or evolve overexpression to detoxify dietary alkaloid loads in excess of what can be efficiently sequestered. With the exception of *serpina1*-like, candidate DEGs in our study are functionally uncharacterized in frogs and could influence sequestration directly or indirectly through effects on transport, detoxification, stress tolerance, or immune regulation (Table 1).

Candidate DEGs from several superfamilies showed uniform directions of expression in *Epipedobates* compared to the trace-accumulating lineages. For example, our results revealed uniform overexpression by *Epipedobates* of candidate DEGs from the glutathione S-transferase (GST) superfamily, which are central to xenobiotic detoxification across diverse taxa (Fig. 5; Tables 1, 2). Brassicales plants defend themselves against herbivores with glucosinolates that are hydrolyzed into toxic, sulfur-containing compounds called isothiocyanates. Over half of the 34 GSTs from the African cotton leafworm moth *Spodoptera littoralis* were able to detoxify isothiocyanates (Sun et al. 2025). Two GST proteins from the tick *Haemaphysalis longicornis* (HlGST and HlGST2) detoxify diverse organophosphate pesticides commonly used against ticks (Hernandez et al. 2018). *Epipedobates* may overexpress GSTs to deal with a high alkaloid burden. Conversely, *Epipedobates* underexpressed three candidate DEGs (*c3*, *c5*, and *c7*) critical to the terminal-complement cascade, which in turn activates the complement immune system in response to pathogens and microbes (Fig. 5; Tables 1, 2). Perhaps *Epipedobates* maintains a dampened complement response compared to trace-accumulating species partly because alkaloids hinder microbial and fungal growth. Skin alkaloids in *O. pumilio* are known to do this (Mina et al. 2015; Hovey et al. 2018). Consistently, the evolution of high alkaloid levels in the skin of sequestering dendrobatids is associated with restructuring of the microbial community (Caty et al. 2025). *Epipedobates* might also be minimizing complement-driven cellular damage and inflammation due to chronically high alkaloid levels.

Candidate DEGs from other superfamilies were divergent in their direction of expression within *Epipedobates*. Two solute carriers (SLCs) (*slc4a7* and *slc16a10*) were overexpressed in *Epipedobates*, while one (*slc16a6*) was underexpressed. SLCs transport a broad range of substrates, and these genes are plausible candidates for mediating multiple aspects of dendrobatid biology. The contrasting expression patterns are therefore unsurprising (Fig. 5; Tables 1, 2). Liver-expressed SLCs have previously been proposed as contributors to sequestration, including possible bile-acid-associated alkaloid transport (Clark et al. 2012; Dawson and Karpen 2015; Caty et al. 2019), and other SLCs have repeatedly emerged as candidates in affecting frog coloration, including melanosome transport, carotenoid uptake, and chromatophore differentiation (supplementary table S5; Rodríguez et al. 2020; Rubio et al. 2024; Monteiro et al. 2025; Mantzana-Oikonomaki et al. 2026). SLCs thus may contribute both to alkaloid handling and to the evolution of warning coloration.

Candidate DEGs from the heat shock protein (HSP) superfamily also showed divergent patterns; HSPs primarily function as molecular chaperones involved in protein folding, assembly, and immune or stress responses, but they can also bind small molecules (Tables 1, 2). Notably, alkaloids are known to interact with heat shock proteins and can either inhibit or promote their expression (Wisén and Gestwicki 2008). For example, isoquinoline alkaloids from *Monoon longifolium* (the false ashoka tree) appear to bind HSP70-x of the parasite *Plasmodium falciparum* and inhibit its function (Shrestha et al. 2025); nicotine suppresses *hsp70* expression and induces apoptosis in human cancer cells (Khalaf and Mohammed 2022); and sanguinarine induces multiple HSP genes in *Arabidopsis*, increasing heat tolerance (Hara and Kurita 2014). In poison frogs, three HSP genes were overexpressed in wild-caught relative to alkaloid-free laboratory *O. sylvatica*, including one of our candidate DEGs (*hspa2*), and a thermal proteome assay showed that another (*hsp90aa1*) binds the alkaloid decahydroquinoline (Caty et al. 2019). We therefore identify HSPs as strong candidates for direct alkaloid binding, toxin-induced stress responses, or both in poison frogs. Interestingly, we found contrasting expression patterns for two *hsp70* genes (*hspa2* and *hspa8*) and two *hsp40* genes (*dnajc19* and *dnajc4*), which was unexpected given that members of the HSP70 and HSP40 families commonly function jointly within chaperone networks (Tables 1 and 2), suggesting complex modulation of these pathways.

We recovered broad biological functions that overlapped with those found in experimental studies of the oldest of the three dendrobatid origins of sequestration (at the base of Dendrobatinae at ∼25 mya). Our GO analyses were highly congruent with transcriptomic gene sets previously implicated in sequestration-related processes, repeatedly highlighting small-molecule transport, metabolism, and immune pathways as central components of sequestration (Caty et al. 2019; O’Connell et al. 2021; Alvarez-Buylla et al. 2022). However, 20 of the 23 candidate DEGs recovered in our analysis were not recovered in these same studies. Although our study contrasted liver gene expression in sequestering versus trace-accumulating frogs from the wild, the experimental studies of Dendrobatinae have tended to contrast a single sequestering species on alkaloid-rich versus alkaloid-free diets (but see Sanchez et al. 2019).

Of the three candidate DEGs identified here and in previous studies of alkaloid metabolism in dendrobatid liver (*hspa2*, *c3*, and *serpina1*-like), there are differences in patterns of expression. As expected, *Epipedobates* overexpressed *serpina1*-like, consistent with overexpression of this gene in alkaloid-administered *O. sylvatica* (O’Connell et al. 2021; Alvarez-Buylla et al. 2023). In contrast, *O. sylvatica* overexpressed *hspa2* and *c3* following alkaloid consumption, whereas *Epipedobates* in our dataset showed reduced expression of these genes relative to trace-accumulating species (O’Connell et al. 2021) (Fig. 5; supplementary table S5). This contrast may indicate that different alkaloid-sequestering dendrobatids have evolved distinct strategies for balancing chemical defense with the costs of immune activation.

A challenge in studying natural frog populations is that measures of skin alkaloid load reflect cumulative lifetime exposure, whereas measures of other variables, such as stomach arthropod diversity (Martin et al. 2025) and constitutive gene expression (e.g., Caty et al. 2019), capture narrower time windows. This mismatch complicates efforts to directly link alkaloid abundance with these traits. Nonetheless, we found marginal support for associations between gene expression and finer-scale variation in alkaloid abundance within *Epipedobates*. Of the three key modules that were significantly associated with sequestration and potentially correlated with alkaloid abundance among individuals, two showed concordant directional relationships across these analyses: lightyellow was positively associated with sequestration and positively correlated with alkaloid abundance, whereas yellow was negatively associated with both. The yellow module followed the expected categorical pattern, differing significantly between trace-accumulating and sequestering species but not within either category. Conversely, lightyellow showed a more complex species-level pattern: although positively associated with sequestration overall, its highest ME values occurred in the trace-accumulating species *S. flotator* (Fig. 6a).

While the yellow module lacked significant GO enrichment, the two candidate DEGs in the yellow module (*apob* and *serpinc1*) contained counterparts from the same superfamilies in lightyellow (*apoh* and *serpinf2*) that contributed to blood clotting, an overrepresented function of lightyellow. Nonetheless, of the two candidate DEGs in yellow, only *serpinc1*, which encodes the clotting-related serpin Antithrombin III, is directly implicated in blood clotting (Table 1) (Whitfield et al. 2004; Pike et al. 2005). The yellow module may have clotting-adjacent functions and be involved only partly in the physiology of alkaloid sequestration; given that lightyellow had only moderate representation of DEGs (∼50%), it is plausible that the function of this module is dominated by lineage-specific physiological requirements.

In contrast, the blue module was strongly positively associated with sequestration but showed marginal support for a negative correlation with alkaloid abundance within *Epipedobates* (Table 2). One possible explanation is that these associations reflect physiological tradeoffs operating at different scales. Across species, elevated expression of blue-module genes, which were overrepresented for functions related to stress response, immunity, and xenobiotic handling, could facilitate alkaloid metabolism in sequestering frogs, for example, by detoxifying or clearing alkaloid loads that exceed sequestration capacity. Within *Epipedobates*, however, individuals or populations with especially large alkaloid loads may reduce expression of these pathways to limit the physiological costs of sustained stress or immune activation. This interpretation remains tentative given the marginal support for the within-*Epipedobates* correlation. Overall, these results suggest that gene-expression patterns associated with the presence of sequestration across species are not necessarily the same as those correlated with variation in alkaloid abundance within sequestering frogs.

Confirming precise roles for candidate DEGs and their pathways will require functional validation, for example experiments with standardized alkaloid administration coupled with molecular and proteomic approaches. As functional genomic tools (e.g., CRISPR-Cas9) become increasingly tractable in non-model frogs, targeted manipulation of candidates could establish causal roles of candidates in sequestration pathways.

## Conclusion

Diet-derived skin alkaloids function as antipredator defenses in dendrobatid poison frogs, with toxin profiles shaped by predator-mediated selection and ecological alkaloid availability. Our results indicate that repeated origins of dendrobatid alkaloid sequestration involve at least two complementary evolutionary processes: expansion of ligand-binding serpins and coordinated changes in gene expression. SerpinA analyses revealed diversification of alkaloid-binding proteins (ABG1 and ABG2) and BBS-like serpins, suggesting expansion and functional divergence of ligand-binding paralogs. Gene-expression analyses of *Epipedobates*, the youngest sequestering dendrobatid lineage, identified shifts involving small-molecule transport, metabolism, detoxification, and immune function. Selection across variable predator and prey communities likely produced a mosaic of convergent and divergent molecular mechanisms among dendrobatid origins. ABGs may have been repeatedly recruited within Dendrobatidae, whereas dietary toxin acquisition in other taxa may rely on distinct genes or gene families.

## Materials and Methods

### Phylogenetic Reconstruction and Analysis of the SerpinA Gene Family in Dendrobatids

Recent proteomic studies revealed that a member of the gene group *serpina1*-like encodes an alkaloid-binding globulin (ABG) in *O. sylvatica*, *D. tinctorius*, and *E. tricolor* (Alvarez-Buylla et al. 2023). To investigate the evolution of the broader serpinA gene family, we surveyed published amphibian genomes (N = 37 species), transcriptomes (N = 4), and PCR sequences (N = 10) for serpinA genes, including members of other serpinA subfamilies for phylogenetic context and *Latimeria chalumnae* as a taxonomic outgroup. We retrieved annotated sequences from NCBI using keyword searches and aligned them using a codon-informed method in DECIPHER version 3.0.0 (Wright 2016). Phylogenies were reconstructed in IQ-TREE version 3.0 (Wong et al. 2026) using ModelFinder for model and partition selection and 1,000 ultrafast bootstrap replicates with the --bnni optimization option. We conducted additional protein-similarity searches against the ClusteredNR database to address inconsistent gene nomenclature and subsequently harmonized names according to phylogenetic placement. Complete sequence metadata, alignments, classifications, and amino acid sites associated with PTX-**251D** binding are provided in supplementary table S1; additional details concerning sequence retrieval, nomenclature, and phylogenetic analyses are provided in the Supplementary Methods.

### SerpinA Transcript Isolation from de novo Assemblies and Targeted Read Baiting

Because ABGs from *O. sylvatica*, *D. tinctorius*, and *E. tricolor* and BBSs from hylids and centrolenids are encoded by liver-expressed transcripts (Taboada et al. 2020; Alvarez-Buylla et al. 2023; Aguilar-Gómez et al. 2026), we prioritized liver transcriptomes while also examining gut/intestine and skin transcriptomes from *O. sylvatica*, *E. anthonyi*, *E. tricolor*, and *Mantella aurantiaca*. We reannotated the existing *M. aurantiaca* transcriptome using genomic serpinA references and reassembled and reannotated published *O. sylvatica* transcriptomes from raw reads. We also generated transcriptomes from the liver, skin, and intestines of one *E. tricolor* collected in Chazojuan, Ecuador, and two *E. anthonyi* collected in Santa Isabel, Ecuador, under St. John’s University IACUC protocol AUP 1965. RNA extraction and library construction followed Ortiz et al. (2025), and 150-bp paired-end sequencing generated approximately 70–97 million read pairs per sample. Raw reads were processed using the Pincho version 0.1 pipeline (Ortiz et al. 2021), which integrates read cleaning and error correction, multiple *de novo* assemblies, consensus construction, quality assessment, and annotation. Because existing annotations and preliminary searches indicated unusually high paralog/variant diversity within the ABG and BBS-like clades, we used a modified Pincho workflow incorporating targeted read baiting to improve recovery of these transcripts from *Epipedobates*. Bait sequences were obtained from the *O. sylvatica* transcriptome, dendrobatid genomes, and previously published ABGs, and we evaluated the targeted-reconstruction procedure by applying it to 11 broadly expressed housekeeping genes (supplementary table S2a,c). Data sources, accession numbers, assembly-quality metrics, and complete processing, baiting, assembly, annotation, and validation procedures are provided in supplementary tables S1 and S2a,c and the Supplementary Methods.

### Sample Collection for Gene-Expression Analysis

Sequestering samples included multiple populations from two *Epipedobates* species (N = 27; all from Ecuador): *E. anthonyi* from Moromoro (MOR-A; N = 4), Zapotillo (ZAP-A; N = 5), San Rafael de Sharug (RAF-A; N = 4), and Santa Martha (MAR-A; N = 4), and *E.* aff. *espinosai* from Pedro Vicente Maldonado (PVM-E; N = 3), Pachijal (PCJ-E; N = 3), and Caimito (CTO-E; N = 4). Trace-accumulating samples included *H. awa* from near Pedro Vicente Maldonado, Ecuador (N = 4), and *S. flotator* from Soberanía National Forest near Pipeline Road, Gamboa, Panama (N = 3). The Panamanian *S. flotator* samples were collected in August 2022; all Ecuadorian samples were collected in May and June 2023 (Fig. 1b; supplementary table S3a).

Frogs were euthanized by applying 20% benzocaine to the venter. Livers were dissected in the field, stored immediately in RNAlater, and maintained under refrigerated conditions during fieldwork to minimize RNA degradation while avoiding freeze-thaw cycles. Samples were then transported at room temperature and transferred to −80°C storage upon return to the laboratory. Skins were also sampled for later alkaloid extraction, profiling, and quantification to confirm that alkaloid abundances accorded with our phenotypic categories (trace-accumulating and sequestering) (Fig. 1; Tarvin et al. 2024; Supplementary Methods and Results).

Total RNA was extracted using a dithiothreitol protocol with the PureLink RNA Mini Kit and on-column digestion with the PureLink DNase Set (Thermo Fisher Scientific), according to the manufacturer’s instructions. In place of the standard mortar-and-pestle or homogenizer procedure, tissue was lysed with acid-washed glass beads (G1277) on a bead beater for 40s. The RNA quality of seven randomly selected samples was verified on a Bioanalyzer 2100 (Agilent) at the Genomic Sequencing and Analysis Facility (GSAF) (University of Texas at Austin, TX, USA). All samples had RIN values of ∼9. Library preparation and sequencing with the 3’Tag-Seq (Tag-Seq) method were completed at the GSAF. Tag-Seq libraries use oligo-dT priming to capture polyadenylated transcripts, enriching for mRNA without a separate rRNA-depletion step.

### Reference Transcriptome Annotation

We used the *O. sylvatica* liver transcriptome assembled with Trinity by Caty et al. (2019) (file: *Osylvatica_transcriptome_Catyetal.fasta*). We annotated this transcriptome with Pincho, a modular *de novo* transcriptomics workflow that includes redundancy reduction (CD-HIT) and completeness assessment (BUSCO) (Ortiz et al. 2021). Functional annotation of the *O. sylvatica* reference was implemented in Pincho, performed by querying transcripts against UniProt (Swiss-Prot), TrEMBL (Amphibia), and KEGG databases to obtain best-hit identifications and associated functional information. We then applied an AT-split/ID-normalization step to (i) resolve A/T-rich chimeric contigs and (ii) standardize transcript and gene identifiers. For this step, we used the recently assembled and annotated genome of the dendrobatid *Ranitomeya imitator* (aRanImi1 (GCF_032444005.1)) (Rhie et al. 2021) as a reference.

After annotating the *O. sylvatica* reference transcriptome, we found that some serpinA isoforms (all *serpina1*-like and *serpina6*-like, many *serpina10*) had been assigned LOC identifiers in our pipeline, indicating that automated annotation had not confidently assigned them to a named gene family. However, the transcript identities had already been established as part of the comprehensive search performed for the serpinA phylogenetic analysis. To ensure that no serpinA fragments were missed during gene-level annotation, we also searched the annotation output for LOC-assigned transcripts that shared Trinity DN identifiers with phylogenetically identified serpinA sequences or that retained expected serpinA-related protein identifiers, such as A1AP/A1AT or ZPI, despite lacking informative gene names. Candidate LOC-assigned fragments were then validated by BLAST. We manually curated gene identifiers for confirmed serpinA transcripts in place of LOC identifiers before downstream expression analyses. The resulting annotated assembly for *O. sylvatica* was used as the reference transcriptome for downstream expression analyses.

Because dendrobatids possess multiple *serpina1* paralogs whose relationships among loci remain unresolved, we conservatively referred to sequences in the reference transcriptome as “*serpina1*-like” rather than assigning specific orthologous identities. Liver-expressed *serpina1*-like paralogs have not been identified for several focal species, including *E.* aff. *espinosai*, *S. flotator*, and *H. awa*, and the short 3′ reads generated by Tag-Seq have limited power to distinguish closely related paralogs or transcript isoforms; the recovered *serpina1*-like signal may therefore reflect gene-clade-level expression rather than any single paralog.

### Read Processing, Alignment, and Expression Quantification

After removing poly(A)-rich, short, duplicate, and low-quality reads, we assessed preprocessing quality and quantified each Tag-Seq library against the *O. sylvatica* reference transcriptome using Salmon version 1.10.2 (Patro et al. 2017). Transcript-level TPM and count estimates were imported into R version 4.3.3 (R Core Team 2025) using tximport version 1.30.0 (Soneson et al. 2015). Separately, reads were aligned to the reference transcriptome using BWA version 0.7.17-r1188 (Li and Durbin 2009); differences in mean alignment percentages among species were evaluated using ANOVA followed by Tukey’s HSD test. Complete preprocessing and quality-control procedures are provided in the Supplementary Methods. Unless otherwise specified, all subsequent analyses were performed in R.

### Filtering and Normalization

Genes with ≥1 NA values across individuals were removed to prevent conflating mapping errors with genuine zero counts. TPM-normalized transcript counts from the same gene were summed with tximport to consolidate multiple isoforms into single gene-level estimates. We removed all transcripts with LOC identifiers from the *O. sylvatica* annotation. We then retained only genes with non-zero expression in all samples. A design matrix was generated to classify which samples belonged to sequestering versus trace-accumulating species. The remaining genes and gene counts were run through the edgeR version 4.0.16 (Chen et al. 2025a) functions DGElist and filterByExpr to remove low-expression genes. filterByExpr applied a library-size adaptive count-per-million threshold (CPM) equivalent to 10 counts in the smallest library, sequestering or trace-accumulating, and retained genes that passed the CPM threshold in at least as many samples as the smaller contrast (trace-accumulating; N = 7). For normalization, Trimmed Mean of M-values (TMM) normalization factors were calculated using the edgeR function calcNormFactors. This function computes a scaling factor for each sample as a weighted mean of trimmed log fold-changes relative to the sample whose 75th percentile count is the closest to the mean upper quartile across all samples, correcting for compositional differences between libraries. The filtered gene counts were next converted to log_2_-CPM using the calculated TMM normalization factors and processed with voom to estimate the mean-variance relationship and generate observation-level precision weights for use in linear modeling (Law et al. 2014). The voom:mean-variance trend plot was inspected to verify that the CPM threshold removed noise.

### Differential Gene Expression Analysis

We used limma version 3.58.1 (Ritchie et al. 2015), a linear model-based approach, to identify genes significantly differentially expressed between sequestering and trace-accumulating species. Filtered, voom-normalized counts for the 940 genes retained after expression filtering, each with at least one mapped read in every sample, were used as inputs for limma. We specified a no-intercept design matrix in which samples were assigned to one of the two phenotypic categories (trace-accumulating or sequestering). To account for correlation structure within the dataset, we first estimated the within-sample correlation using the duplicateCorrelation function (Smyth et al. 2005). This consensus correlation estimate was then incorporated into the lmFit function, which fits a linear model separately for each gene (Phipson et al. 2016). We then used makeContrasts to define the pairwise contrast between sequestering and trace-accumulating species, and fitted this contrast to the linear model using contrasts.fit. Moderated t-statistics were calculated with eBayes, which applies empirical Bayesian shrinkage to gene-wise standard errors, stabilizing variance estimates across genes while accounting for mean-variance relationships in the data. The determination of DEGs was made using decideTests, with p-values adjusted using the BHFDR correction. Genes with an adjusted p-value ≤ 0.05 were considered significantly differentially expressed for all downstream analyses.

### WGCNA

To identify co-expression modules associated with phenotypic category, we conducted a WGCNA using the WGCNA R package version 1.73 (Langfelder and Horvath 2008). Pearson gene-gene correlations were transformed into an unsigned weighted network using the lowest soft-thresholding power between 1 and 20 that produced a scale-free topology fit of R² ≥ 0.85. Modules were identified from topological-overlap dissimilarity using the blockwiseModules function, with a minimum module size of 15 genes. We summarized each module using its ME and tested associations between MEs and phenotypic category using Pearson correlations and Student asymptotic p*-*values, followed by BHFDR correction across module-trait tests. For significant modules, we tested species-level differences in ME values using pairwise Welch two-sample t*-*tests without pooling standard deviations, with BHFDR correction applied within each module-trait combination. We used Fisher’s exact tests to identify modules enriched for limma-identified DEGs and selected four modules for further investigation because they were both significantly enriched for DEGs and significantly associated with phenotypic category (FDR ≤ 0.05). Within these modules, module membership (kME) was calculated as the correlation between each gene’s expression profile and its ME. Networks were visualized in Cytoscape (Shannon et al. 2003); we retained genes that were differentially expressed (adjusted *p* ≤ 0.05) or had high module membership (kME ≥ 0.6) and connections with topological-overlap values ≥ 0.02 (supplementary figs. S12–S15). Additional network-construction and visualization details are provided in the Supplementary Methods.

### ME Comparisons Among Species and Expression Correlations with Alkaloid Abundance

To evaluate whether ME values differed among species, we performed pairwise Welch two-sample t-tests using the base R function pairwise.t.test without pooling standard deviations. Pairwise p-values were corrected within each module-trait combination using BHFDR, with adjusted p ≤ 0.05 considered significant. Module eigengene values were plotted in faceted boxplots, with letters indicating significant pairwise differences (supplementary fig. S11).

To assess whether expression patterns distinguishing *Epipedobates* from trace-accumulating species also tracked finer-scale variation in alkaloid abundance, we modeled voom-normalized log_2_-expression of the 23 candidate DEGs and MEs of the 10 modules associated with phenotypic category as functions of ln-transformed alkaloid abundance using the base R function lm, within *Epipedobates* (we excluded the trace-accumulating species and two *E. anthonyi* individuals lacking paired gene-expression and alkaloid-profile data, one each from MOR and MAR). For each candidate DEG or module, we extracted the slope, unadjusted p-value, R², and Pearson correlation coefficient. Candidate-DEG p-values were adjusted across genes using BHFDR, with adjusted p ≤ 0.05 considered significant. Relationships were visualized using ordinary least-squares regression lines with 95% confidence intervals (supplementary figs. S9 and S16; supplementary table S14).

### Overrepresentation Analysis of GO Terms

To identify biological processes associated with variation between the sequestering and trace-accumulating categories, we performed gene set overrepresentation analyses using the ShinyGO web application version 0.85 (Ge et al. 2020). For each analysis, the foreground gene set comprised genes assigned to one of four key WGCNA modules. Key modules were defined as modules that (1) contained candidate genes; (2) were significantly enriched for DEGs between the sequestering and trace-accumulating phenotypic categories, identified by limma (FDR ≤ 0.05); and (3) were associated with phenotypic category through significant module eigengene– trait correlations (FDR ≤ 0.05). As the background gene set, we used the 3,261 genes with ≥1 read count across all samples. Overrepresentation was evaluated using the 2025 *Homo sapiens* GO biological process database. ShinyGO assessed overrepresentation using hypergeometric tests with BHFDR correction. Enriched terms were ranked by FDR, and the top 10 terms were visualized as bar plots (supplementary tables S12, S13; supplementary figs. S17, S18).

## Supporting information

Supplemental Methods and Results

## Supplementary Material

Supplementary material is available online.

## Data Availability

All raw read data are archived with the National Center for Biotechnology Information (accession number pending). Codes for all analyses, gene expression counts, and figures are available on Zenodo (submission pending). Raw chromatographic data are deposited on the Mass Spectrometry Interactive Virtual Environment (https://massive.ucsd.edu/; accession number pending) and newly generated sequence data are deposited in NCBI.

## Acknowledgments

We are enormously grateful to Gabriela Maldonado and Pablo Quinteros Astudillo for their assistance with fieldwork in Ecuador, and to Amalia Espinoza-Regalado for her counsel regarding field sites. We thank Gabriela Maldonado and Andrea Terán-Valdez (Centro Jambatu) for handling collection and export paperwork. J.L.C. thanks Luke Larter for his assistance with fieldwork in Panama, his extraordinary ingenuity in brainstorming and devising methods, and for saving him numerous times when he was in a pinch. J.L.C. is grateful to Tom Juenger, who for years has generously allowed him to use his bead beater for RNA extractions, including those for this manuscript. J.L.C. is also indebted to Rick Fitch, from whom he learned a great deal about alkaloid chemistry and the identification of challenging compounds. M.I.P. thanks Fundación de Conservación Jocotoco for authorizing data collection within Buenaventura Natural Reserve. We thank the UT Austin Freshman Research Initiative stream EvoDevOmics for assistance with data processing and analysis. We acknowledge the Genomic Sequencing and Analysis Facility at UT Austin, Center for Biomedical Research Support (RRID: SCR_021713), for providing library preparation and Tag-Seq services. We thank the team at UT Austin’s Mass Spectrometry Facility for their GC–MS instrumentation and software services. AI tools were used minimally to refine the text and flow of the article, but did not generate content; all scientific content, analyses, interpretations, and final text were reviewed and approved by the authors.

## Author Contributions

J.L.C.: conceptualization, data curation, formal analysis, funding acquisition, investigation, methodology, resources, validation, visualization, writing—original draft, writing—review and editing; V.L.: conceptualization, formal analysis, data curation, investigation, methodology, visualization, writing—original draft, writing—review and editing; G.Y.R.: formal analysis, methodology, visualization, writing—review and editing; M.A.B.S.: formal analysis, methodology, writing—review and editing; M.I.P.: methodology, project administration, resources, supervision, validation, writing—review and editing; D.S.: methodology, writing— review and editing; M.H.D.: methodology, resources, writing—review and editing; I.M.R.: methodology, resources, writing—review and editing; N.D.C.A.: investigation, methodology, writing—review and editing; M.B.: investigation, methodology, writing—review and editing; J.C.S.: conceptualization, data curation, formal analysis, investigation, methodology, resources, validation, visualization, writing—review and editing; R.L.Y.: conceptualization, funding acquisition, methodology, resources, validation, writing—review and editing; D.C.C.: conceptualization, funding acquisition, methodology, resources, validation, writing—original draft, writing—review and editing.

## Ethics approval

Collection in and export from Panama were performed under Ministerio de Ambiente Permiso de Colecta Científica (No. ARBG-0038-2022) and Permiso de Transferencia de Material Genético y/o Biológico No. PA-01-ARG-096-2022, respectively. Collection in and export from Ecuador were performed under Ministerio del Ambiente, Agua y Transición Ecológica (No. MAATE-DBI-CM-2022-0254) and Ministerio del Ambiente, Agua y Transición Ecológica (Nos. 018-2023-EXP-CM-FAU-DBI/MAATE and 23EC000018E), respectively. The animal use protocols were approved by St. John’s University IACUC (AUP 1965) and the Smithsonian Tropical Research Institute (SI-22017). Voucher specimens are deposited in the Herpetology Division of the UT Biodiversity Collections.

