## Supplementary figures and images for "From diet to defense: serpin duplication and gene-expression evolution as putative contributors to poison-frog alkaloid sequestration"

### Supplemental Figure 1

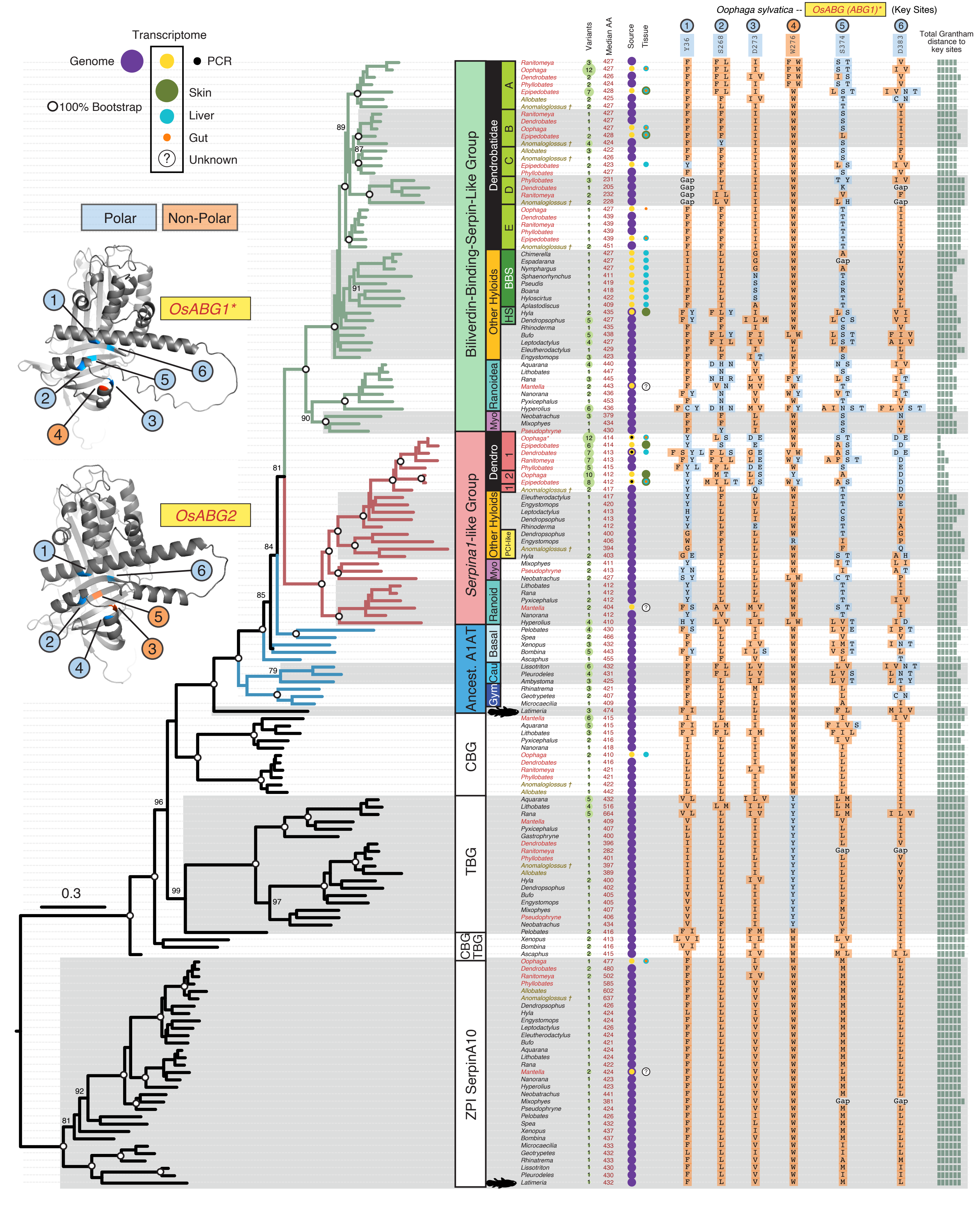

### Supplemental Figure 3

#### Bubble Map of Alkaloid Percent Abundances by Population of *Epipedobates*

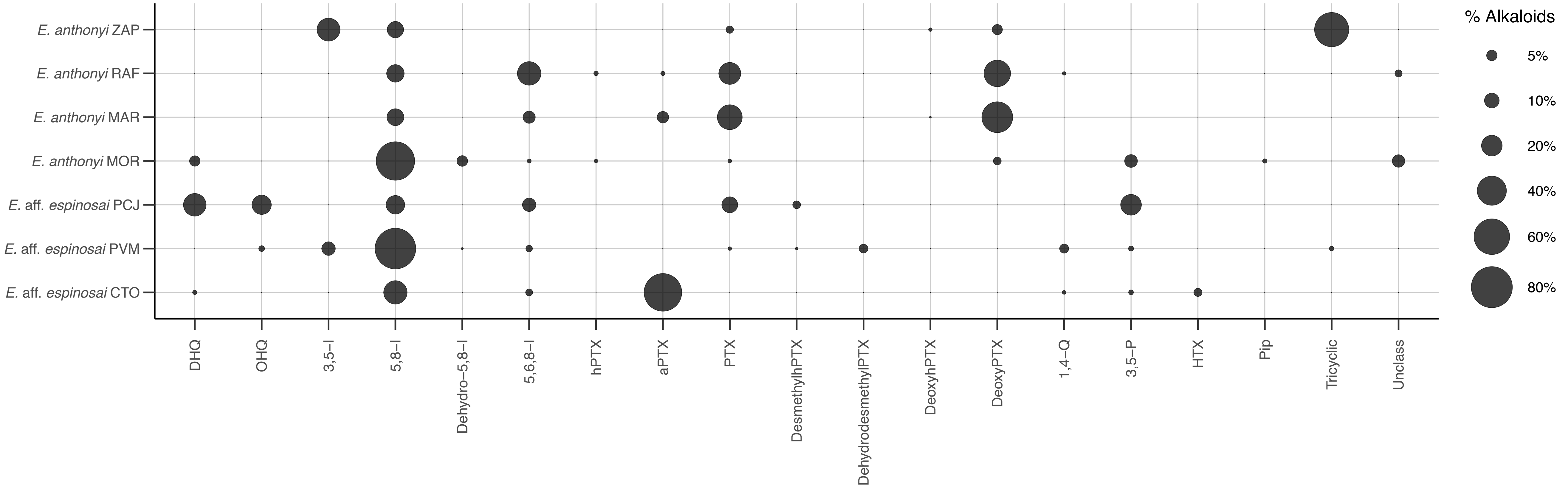

### Supplemental Figure 4

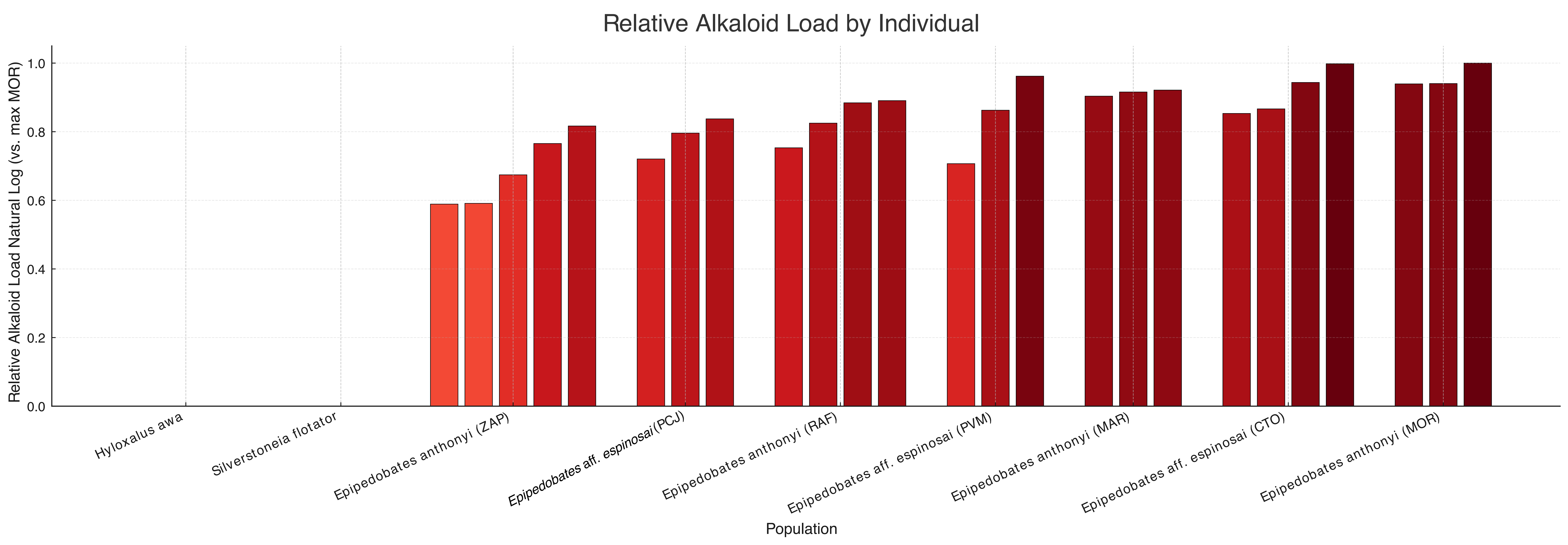

### Supplemental Figure 5

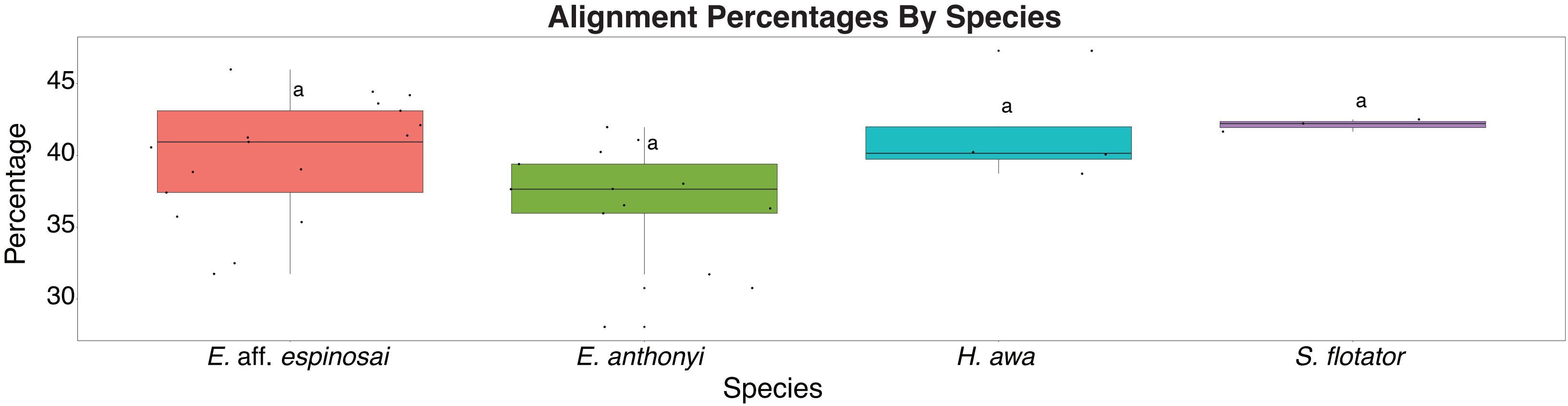


ANOVA Results

| | Df | Sum Sq | Mean Sq | F value | Pr(>F) | |
|---|---|---|---|---|---|---|
| Sample | 3 | 146.3 | 48.75 | 3.042 | 0.0425 | * |
| Residuals | 33 | 528.8 | 16.03 | | | |

### Supplemental Figure 6

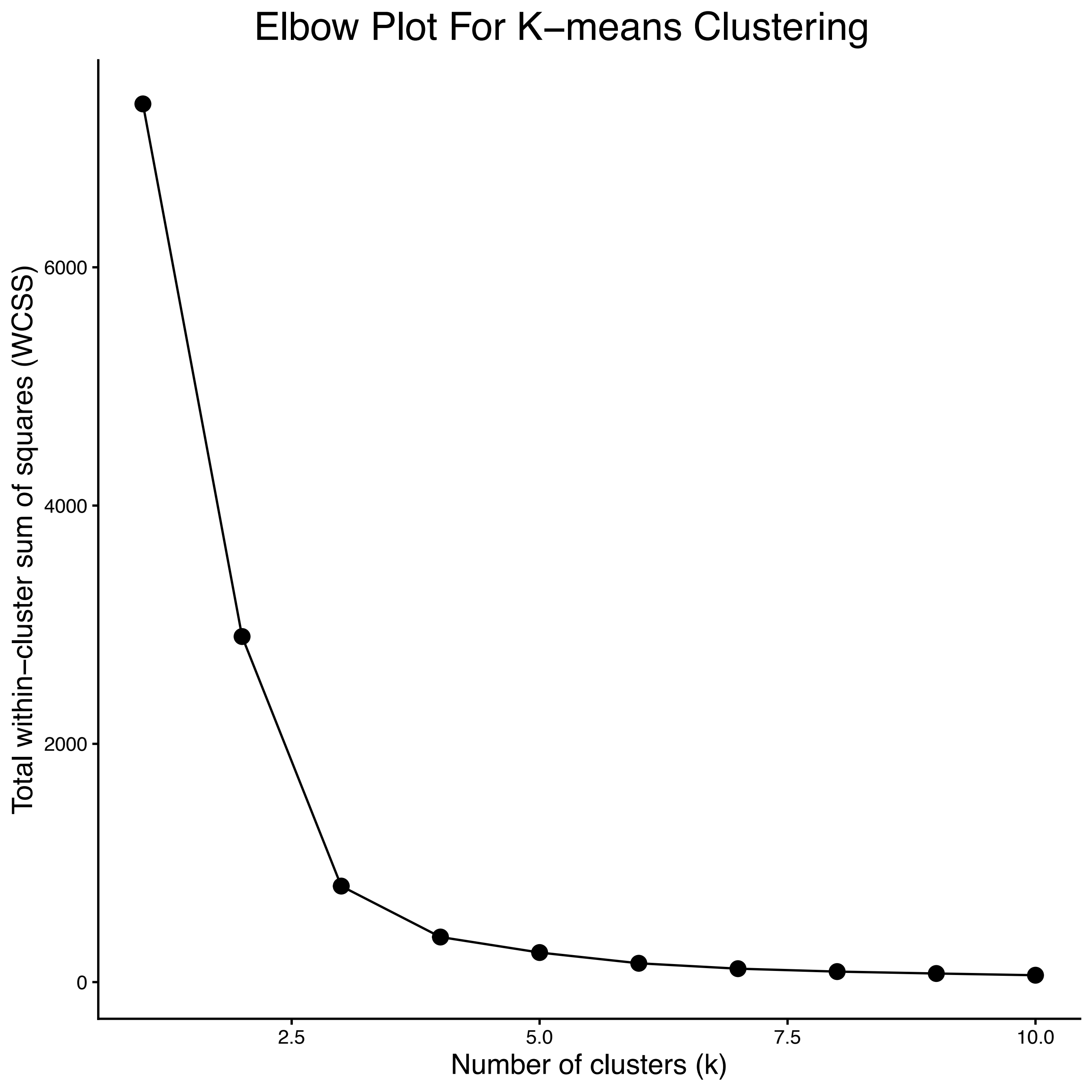

### Supplemental Figure 7

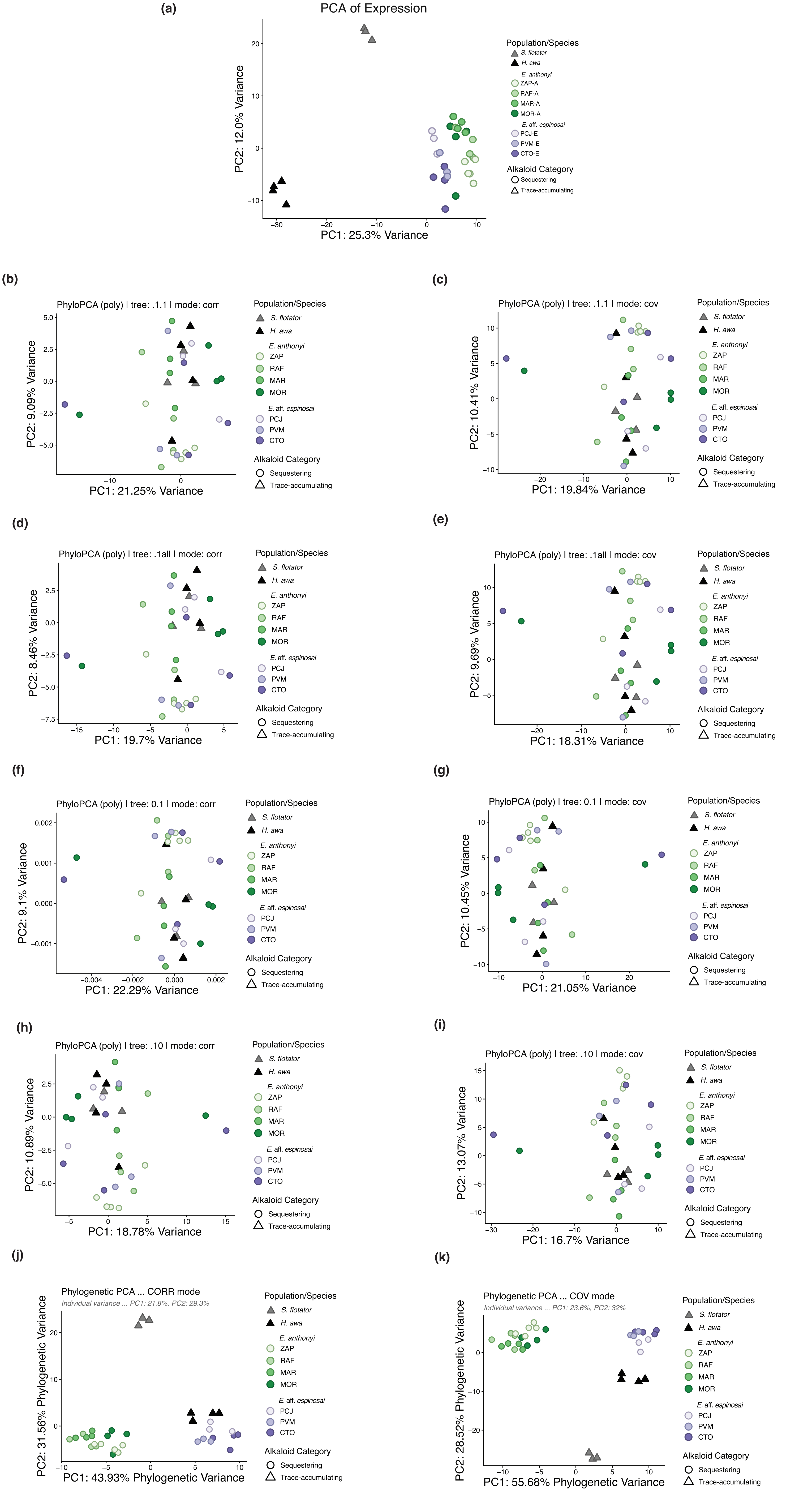

### Supplemental Figure 8

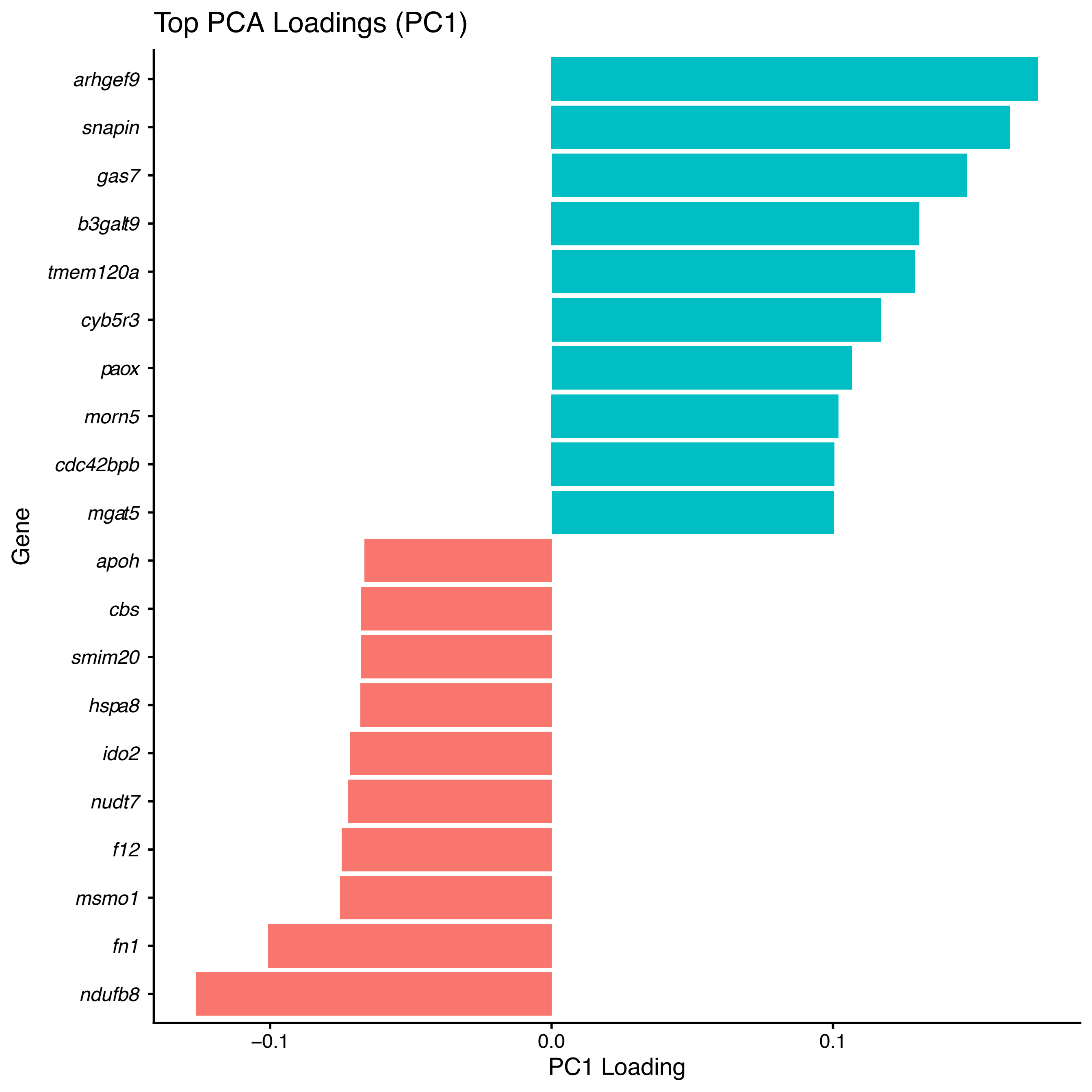

### Supplemental Figure 9

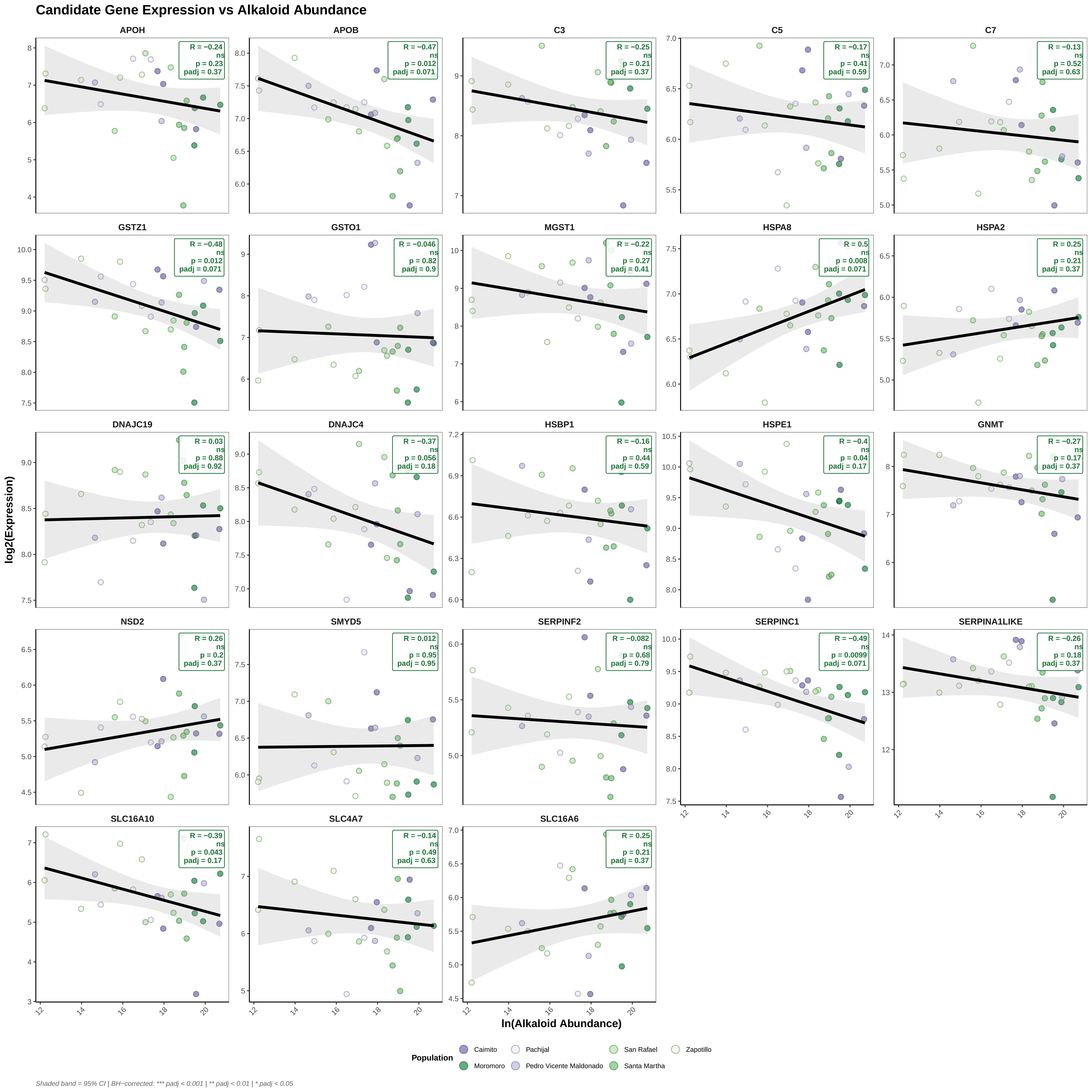

### Supplemental Figure 10

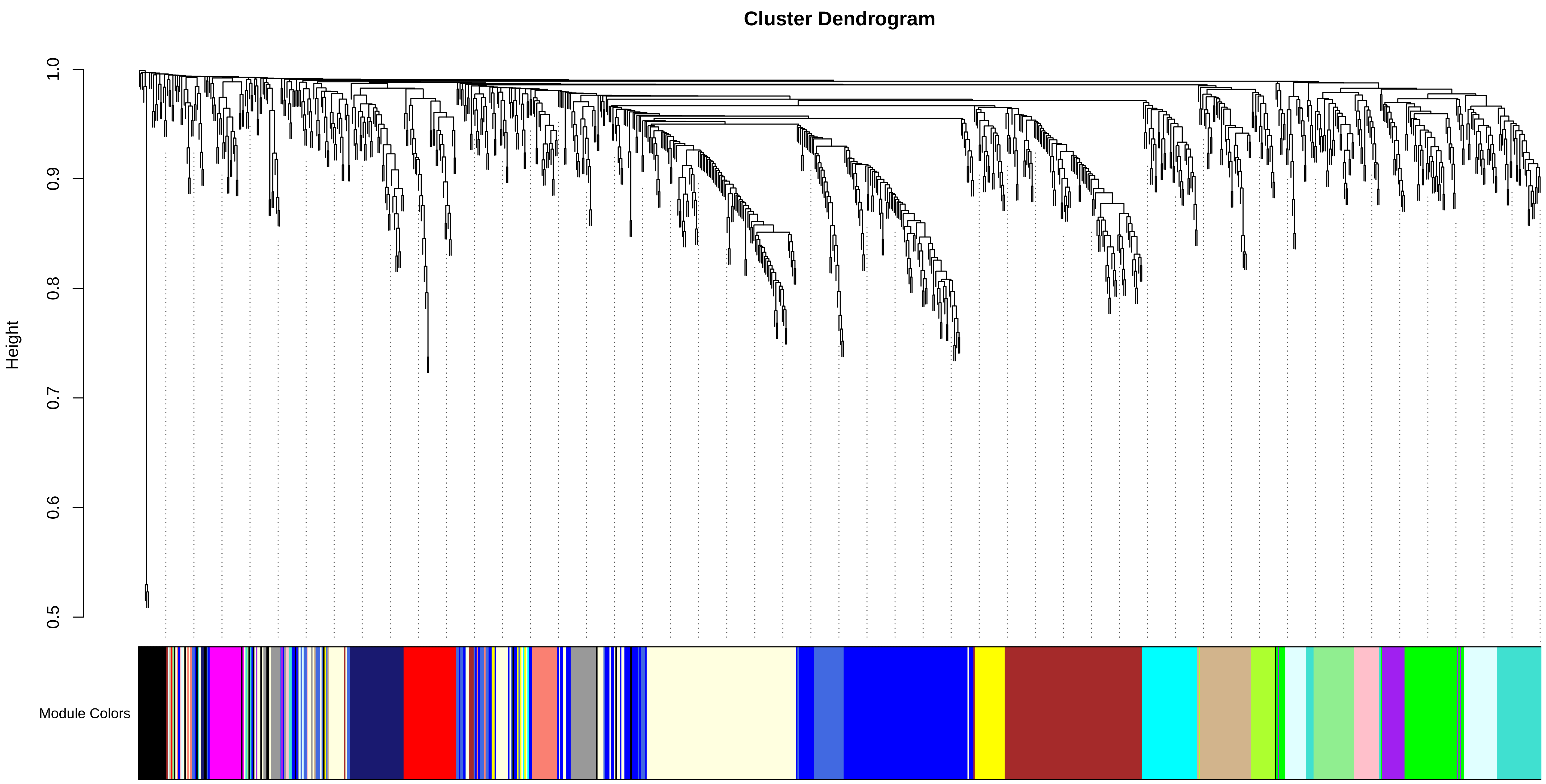

### Supplemental Figure 11

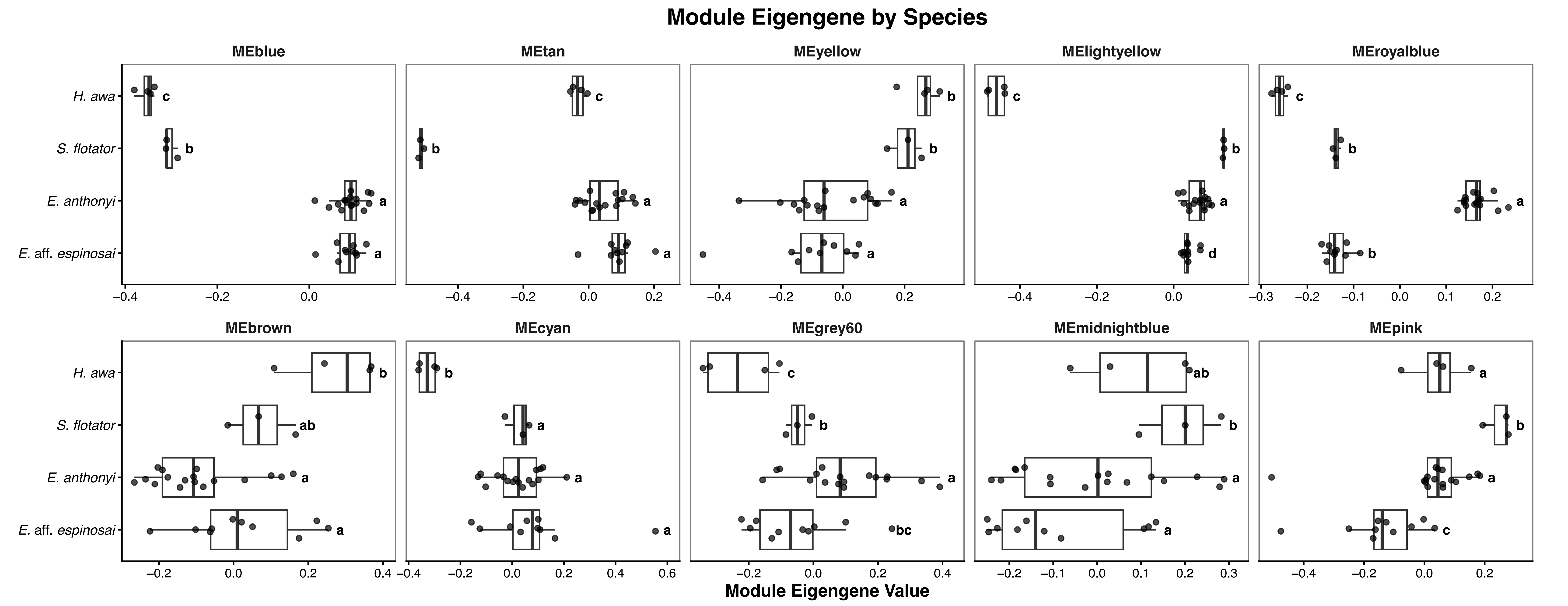

### Supplemental Figure 12

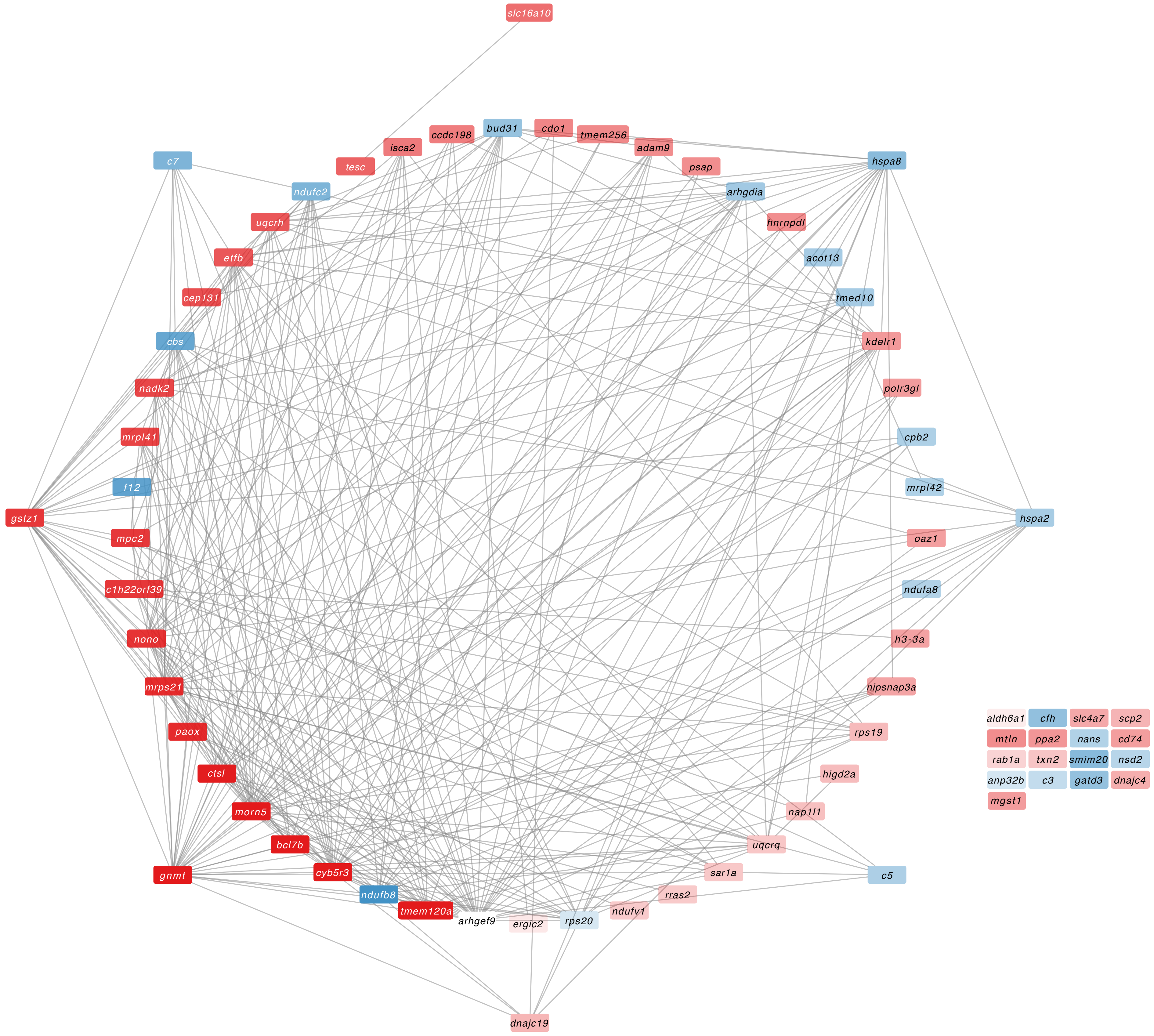

### Supplemental Figure 13

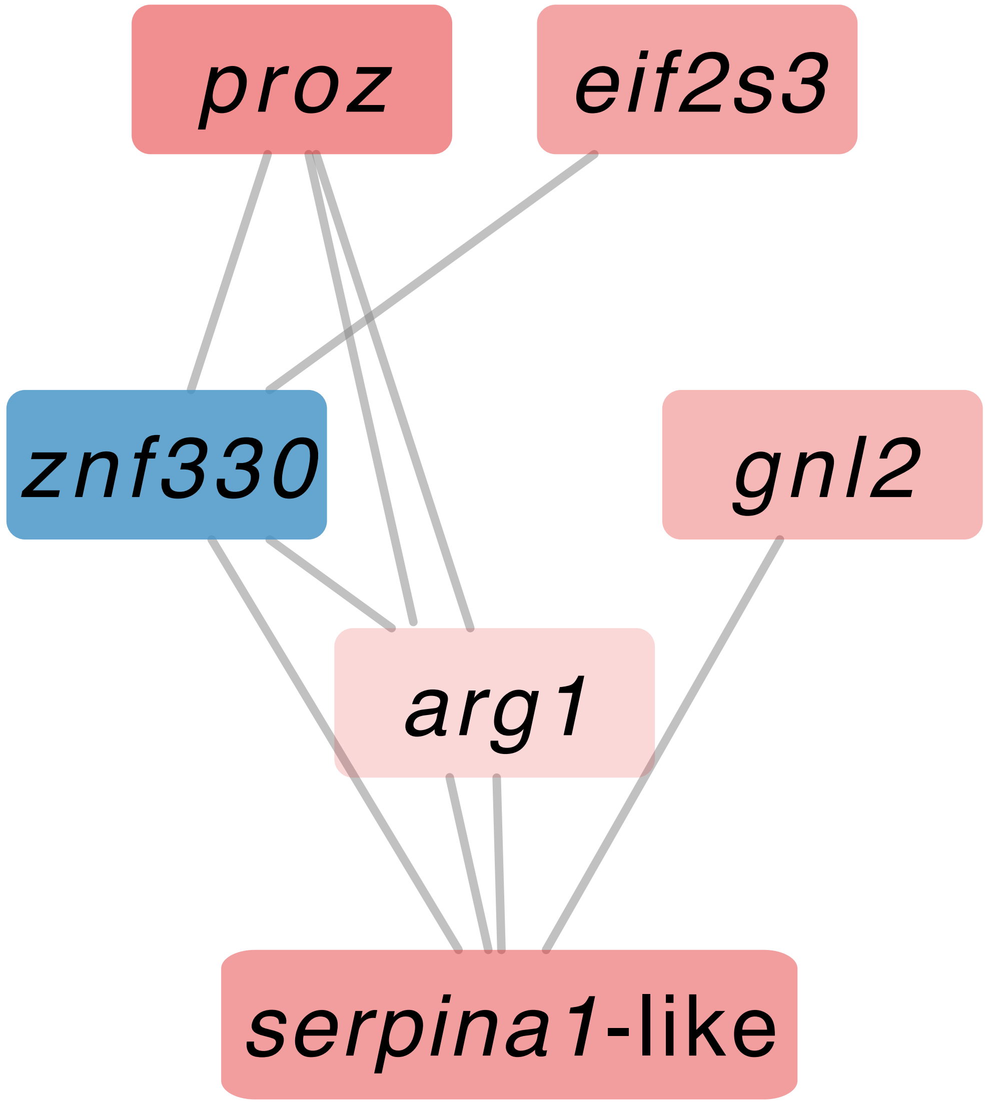

### Supplemental Figure 14

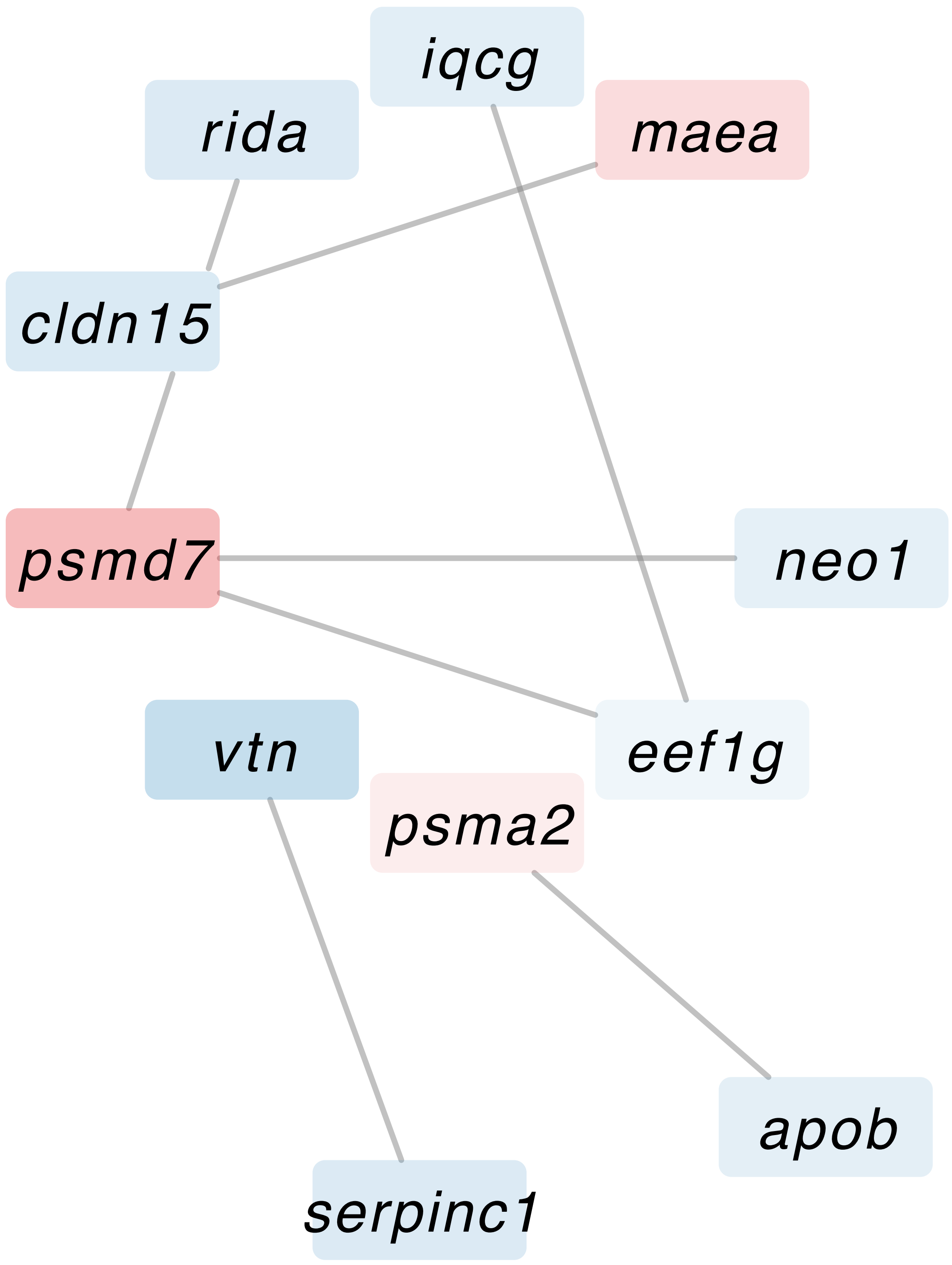

### Supplemental Figure 15

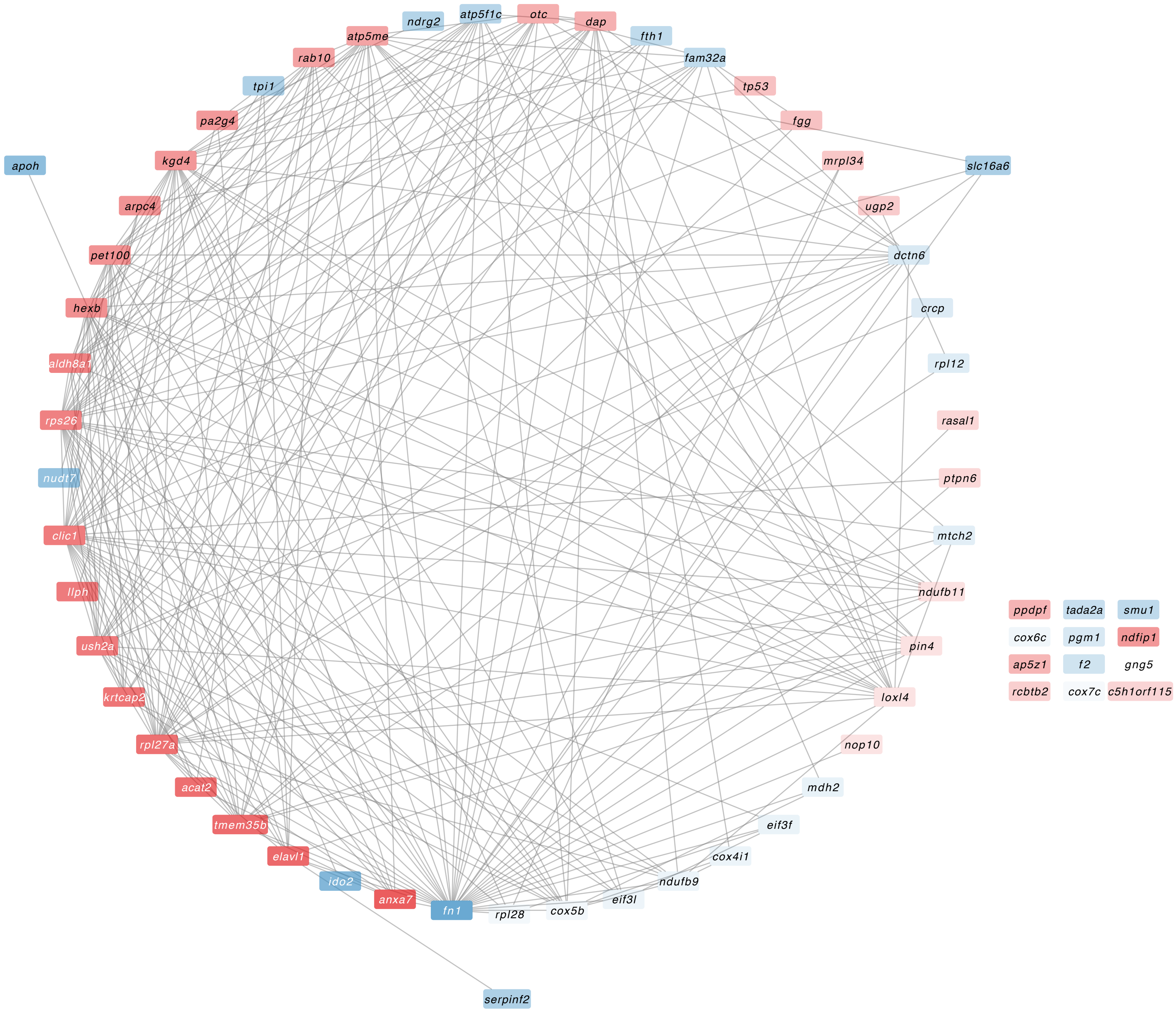

### Supplemental Figure 16

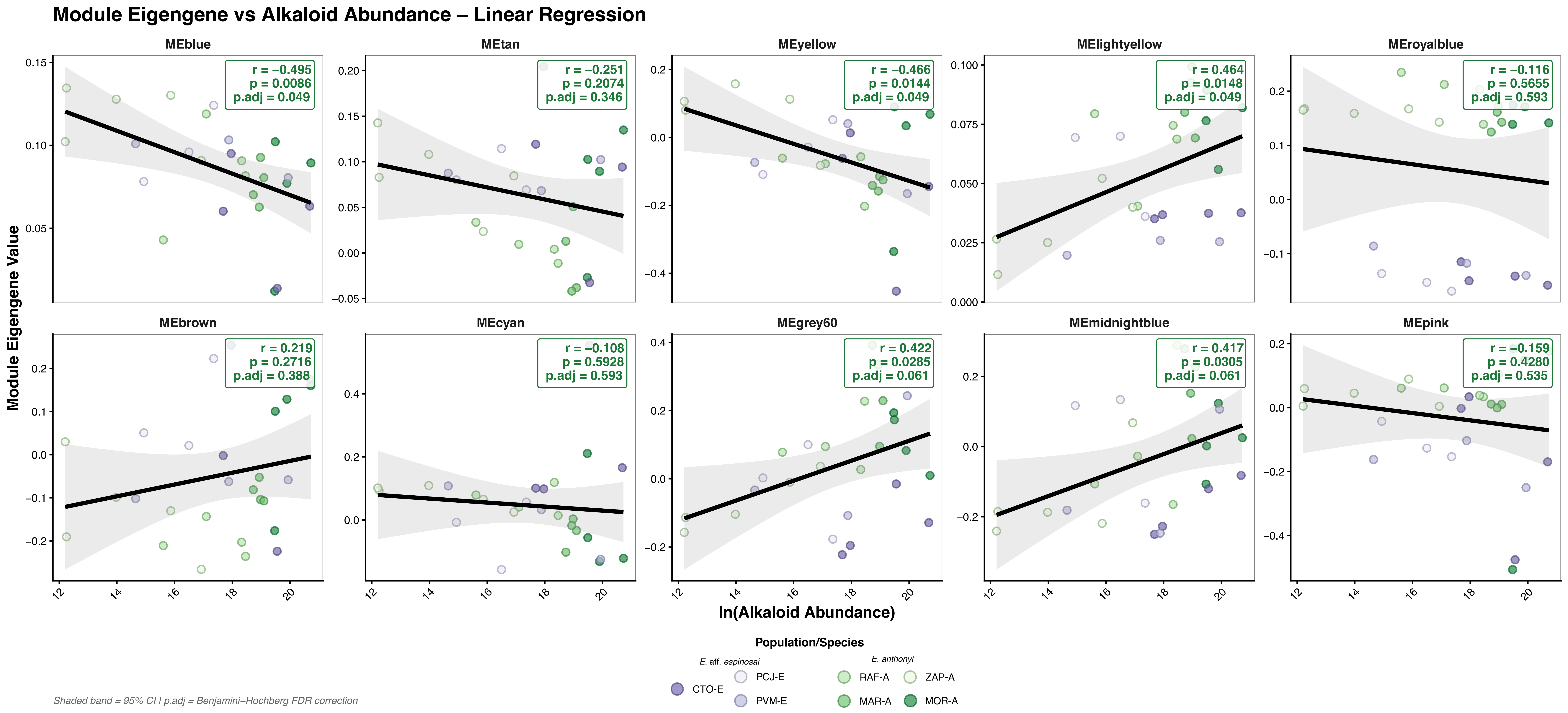

### Supplemental Figure 17

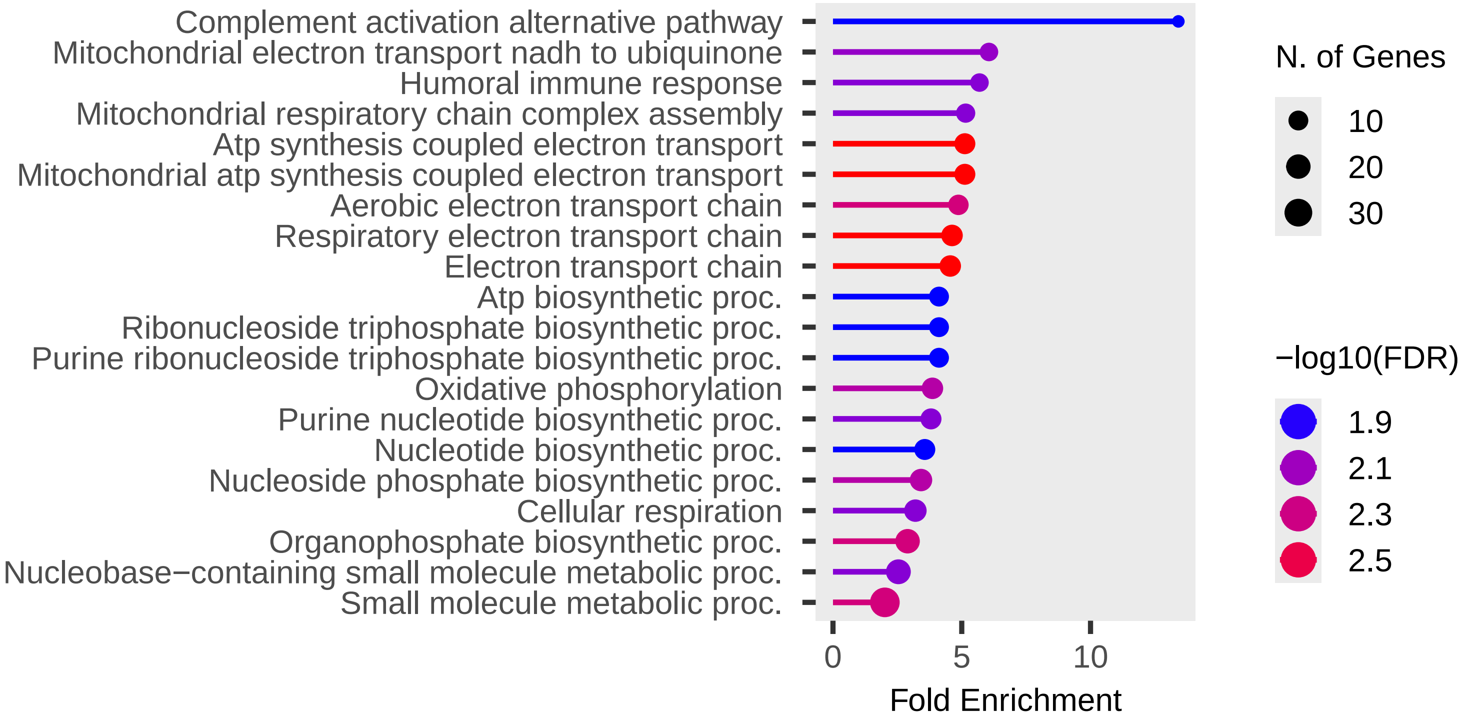

### Supplemental Figure 18

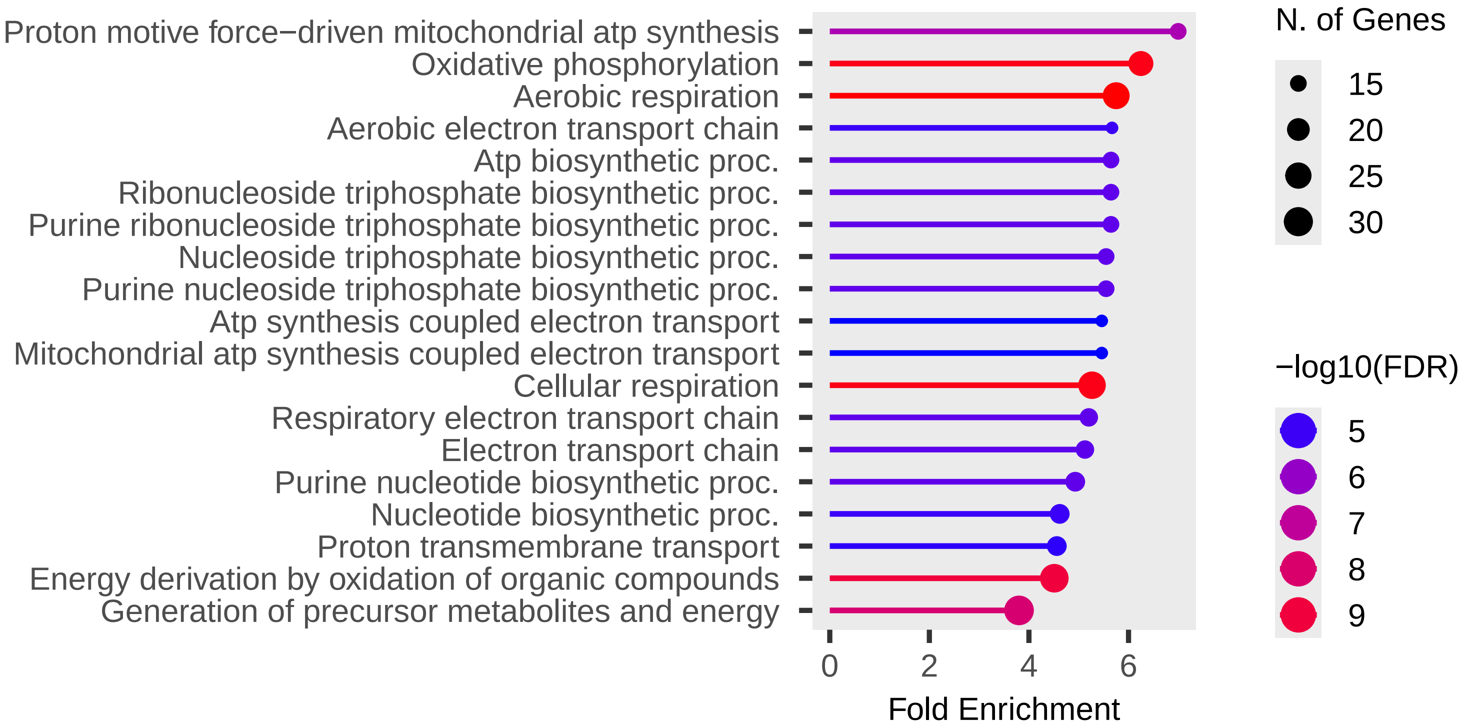
