## Supplemental Figure 2 for "From diet to defense: serpin duplication and gene-expression evolution as putative contributors to poison-frog alkaloid sequestration"

(a)

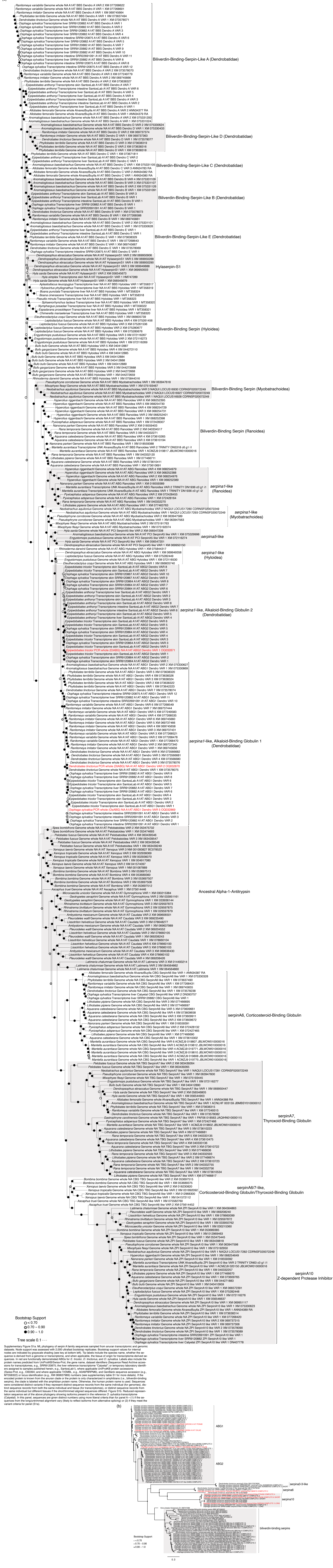


Supplementary Figure S1a. ML phylogeny of serpin A family sequences sampled from anuran transcriptomic and genomic datasets. Node support was assessed with 2,000 ultrafast bootstrap replicates. Bootstrap support values for internal nodes are indicated by grayscale shading (see key at bottom left). Tip labels include the species name, whether the sequence is derived from a genome or transcriptome, and when applicable, the tissue of origin for transcriptome-derived sequences. In red are functionally demonstrated ABGs for *E. tricolor*, *D. tinctorius*, and *O. sylvatica*. Labels also include the protein names predicted from UniProtKB/Swiss-Prot, the gene name, dataset identifiers (Sequence Read Archive accessions for transcriptomes, e.g., SRR9120875; the liver reference transcriptome “Catyetal”; or temporary laboratory identifiers assigned to samples published herein, e.g., SantosLab1), where applicable UniProtKB protein accessions (Swiss-Prot, e.g., O00394; and where applicable TrEMBL, e.g., A0A6P8RPM6), and GenBank sequence accession (e.g., MT358322) or locus identification (e.g., XM 069637989) numbers (see supplementary table S1 for more details). If the encoded protein is known from the anuran clade or the protein is only characterized in amphibians (i.e., biliverdin-binding serpins), the clade is labeled with the amphibian protein name. Otherwise, the human protein name is used. Sequences were considered distinct variants if they represent distinct sequence records from the same individual (for genomes), distinct sequence records from both the same individual and tissue (for transcriptomes), or distinct sequence records from the same individual but different tissues if the short/trimmed aligned sequences differed. Figure S1b. Reduced-representation sequence set of the above phylogeny showing isoforms present in the reference *O. sylvatica* transcriptome (Catyetal). In this panel, sequences are given distinct numbers using more liberal criteria than for panel A—(1) if the sequences from the long/untrimmed alignment vary (likely to reflect isoforms from alternative splicing) or (2) if they meet the variant criteria for panel (S1a).

(b)
