## Supplemental Table 5 for "From diet to defense: serpin duplication and gene-expression evolution as putative contributors to poison-frog alkaloid sequestration"

**Table S5.** Listed in table rows are genes determined to be up- or downregulated in a dendrobatid species after experimental administration of alkaloids. For each gene, columns list the name of the protein for which the gene codes; the gene/protein family and superfamily to which the gene belongs; the main molecular function of proteins in the superfamily; the dendrobatid tissue(s) in which the gene was differentially expressed; the conditions (e.g., the particular alkaloid administered, experimental or natural population) of the contrast between which differential gene expression was evaluated, note that PTX is pumiliotoxin, DHQ is decahydroquinoline; the direction of expression (up- or downregulated) in the livers of alkaloid-administered or higher-alkaloid condition relative to the alkaloid-free or lower-alkaloid condition; the type of methods, i.e., whether the method used assessed differences in protein quantities produced (proteomics) or differences in levels transcription (RNA-seq, aka transcriptomics); the experimental study referenced.

| Gene | Protein | Family, Superfamily | Superfamily Function | Tissue(s) | Contrast and Species | Direction | Evidence Type | Study |
|---|---|---|---|---|---|---|---|---|
| *abca8* | ATP-binding cassette sub-family A member 8 | Subfamily A, ATP-binding cassette (ABC) transporters | Bind xenobiotics | Liver | DHQ-administered vs. alkaloid-free lab *Oophaga sylvatica* | Up | Proteomics | O'Connell et al. (2021) |
| *abca9* | ATP-binding cassette sub-family A member 9 | Subfamily A, ATP-binding cassette (ABC) transporters | Bind xenobiotics | Liver | DHQ-administered vs. alkaloid-free lab *Oophaga sylvatica* | Up | Proteomics | O'Connell et al. (2021) |
| *apoa4* | Apolipoprotein A-IV | APOA1/C3/A4/A5, Apolipoprotein | Lipid transport/ metabolism | Liver | Wild-caught frogs vs. alkaloid-free lab *Oophaga sylvatica* | Up | RNA-seq | Caty et al. (2019) |
| *c3* | Complement component 3 | Complement component C3/C4/C5 | Adaptive and innate immunity | Liver, intestines, skin | Wild-caught frogs vs. alkaloid-free lab *Oophaga* | Up | RNA-seq, Proteomics | Caty et al. (2019); O'Connell |

| | | | | | | | | |
|---|---|---|---|---|---|---|---|---|
| | | Thioester-containing protein (TEP) | | | *sylvatica*; DHQ-administered vs. alkaloid-free lab *Oophaga sylvatica* | | | et al. (2021) |
| *c5ar1* | C5a anaphyla-toxin chemotactic receptor 1 | Receptor 1, G-protein coupled receptor | Binds to an inflammatory complement protein (C5a) to trigger inflammatory cascades | Liver | Wild-caught frogs vs. alkaloid-free lab *Oophaga sylvatica* | Up | RNA-seq | Caty et al. (2019) |
| *cyp1a5* | Cytochrome P450 1A5 | Family 1, Cytochrome p450 | Catalysis of xenobiotic metabolism | Liver | Wild-caught frogs vs. alkaloid-free lab *Oophaga sylvatica* | Up | RNA-seq | Caty et al. (2019) |
| *cyp2k1* | Cytochrome P450 2K1 | Family 2, Cytochrome p450 | Catalysis of xenobiotic metabolism | Liver, intestines | Wild-caught frogs vs. alkaloid-free lab *Oophaga sylvatica* | Up[a] | RNA-seq | Caty et al. (2019) |
| *cyp2k4* | Cytochrome P450 2K4 | Family 2, Cytochrome p450 | Catalysis of xenobiotic metabolism | Liver, intestines | Wild-caught frogs vs. alkaloid-free lab *Oophaga sylvatica* | Up[b] | RNA-seq | Caty et al. (2019) |

[a] Opposite trend in intestines (downregulated in wild-caught frogs compared to alkaloid-free lab frogs)

[b] Same trend in intestines (upregulated in in wild-caught frogs compared to alkaloid-free lab frogs)

| | | | | | | | | |
|---|---|---|---|---|---|---|---|---|
| *cyp4f22* | Cytochrome P450 4F22 | Family 4, Cytochrome p450 | Catalysis of xenobiotic metabolism | Liver | DHQ-administered vs. alkaloid-free lab *Oophaga sylvatica* | Down | Proteomics | O'Connell et al. (2021) |
| *gstk1* | Glutathione S-transferase kappa 1 | Kappa, Glutathione S-transferase (GSTs) | Detoxify xenobiotics by rendering them hydrophilic | Liver, intestines, skin | DHQ-administered vs. alkaloid-free lab *Oophaga sylvatica* | Up | Proteomics | O'Connell et al. (2021) |
| *hsp90aa1* | Heat shock protein 90 alpha | Heat shock protein 90, Heat shock proteins | Protein folding and stabilization | Liver | Wild-caught frogs vs. alkaloid-free lab *Oophaga sylvatica* | Up | RNA-seq, Proteomics | Caty et al. (2019) |
| *hspa1l* | Heat shock 70 kDa protein 1L | Heat shock protein 70, Heat shock proteins | Protein folding and stabilization | Liver | Wild-caught frogs vs. alkaloid-free lab *Oophaga sylvatica* | Up | RNA-seq | Caty et al. (2019) |
| *hspa2* | Heat shock 70 kDa protein 2 | Heat shock protein 70, Heat shock proteins | Protein folding and stabilization | Liver | Wild-caught frogs vs. alkaloid-free lab *Oophaga sylvatica* | Up | RNA-seq | Caty et al. (2019) |
| *mhcia* | Major histo-compatibility complex (MHC) class I alpha | NA, MHC class I-like antigen recognition | Adaptive immunity | Liver and intestines | PTX- and DHQ-administered frogs vs. DHQ- | Up[c] | RNA-seq | Alvarez-Buylla et al. (2022) |

[c] Same trend in intestines (upregulated in in wild-caught frogs compared to alkaloid-free lab frogs)

| | | | | | | | | |
|---|---|---|---|---|---|---|---|---|
| | | | | | administered *Dendrobates tinctorius* | | | |
| *nnmt* | Nicotinamide N-methyl-transferase | NNMT/ PNMT/ TEMT, SAM-dependent methyl-transferase | Methylate proteins or metabolites, which can alter biological activity | Liver, intestines, skin | DHQ-administered vs. alkaloid-free lab *Oophaga sylvatica* | Up[d] | Proteomics | O'Connell et al. (2021) |
| *pfkl* | ATP-dependent 6- phospho-fructokinase | ATP-dependent 6-phospho-fructokinase vertebrate type, Phospho-fructokinase | Catalyzes the first step of glycolysis (phosphorylation) | Liver, intestines, skin | DHQ-administered vs. alkaloid-free lab *Oophaga sylvatica* | Down | Proteomics | O'Connell et al. (2021) |
| *serpina1* | Alpha-1-antitrypsin[e] | Family A, Serpins | Inhibit enzymes that degrade proteins (proteases) | Liver, intestines, skin | DHQ-administered vs. alkaloid-free lab *Oophaga sylvatica* | Up | Proteomics | O'Connell et al. (2021); Alvarez-Buylla et al. (2023) |
| *slc26a3* | Solute carrier organic anion exchanger family member 26A3 | SLC Family 26, Solute carriers (SLC) | Facilitate transmembrane molecule transport | Liver | DHQ-administered vs. alkaloid-free lab *Oophaga sylvatica* | Down | Proteomics | O'Connell et al. (2021) |

[d] Overexpression in the intestines and skin, but not liver, was significant

[e] Encodes alkaloid-binding-globulin in *O. sylvatica* (Alvarez-Buylla et al. 2023)

| | | | | | | | | |
|---|---|---|---|---|---|---|---|---|
| *slc38a2* | Solute carrier family 38 member 2 | SLC Family 38, Solute carriers (SLC) | Facilitate transmembrane molecule transport | Liver | Wild-caught frogs vs. alkaloid-free lab *Oophaga sylvatica* | Up | RNA-seq | Caty et al. (2019) |
| *slc51a* | Organic solute transporter subunit alpha | SLC Family 51, Solute carriers (SLC) | Facilitate transmembrane molecule transport | Liver | DHQ-administered vs. alkaloid-free lab frogs, wild-caught frogs vs. alkaloid-free lab *Oophaga sylvatica* | Up | RNA-seq, Proteomics | Caty et al. (2019); O'Connell et al. (2021) |
| *slco1a2* (alias *oatp1a2*) | Solute carrier organic anion exchanger family member 1A2 | SLCO (former SLC 21), Solute carriers (SLC)—specifically Organic Anion Transporting Polypeptides (OATP) | Facilitate transmembrane molecule transport | Liver | DHQ-administered vs. alkaloid-free lab *Oophaga sylvatica* | Down | Proteomics | O'Connell et al. (2021) |
| *slco2b1* (alias *oatp2b1*) | Solute carrier organic anion exchanger family member 2B1 | SLCO (former SLC21), Solute carriers (SLC)—specifically Organic Anion Transporting Polypeptides (OATP) | Facilitate transmembrane molecule transport | Liver | DHQ-administered vs. alkaloid-free lab *Oophaga sylvatica* | Up | Proteomics | O'Connell et al. (2021) |

| | | | | | | | | |
|---|---|---|---|---|---|---|---|---|
| *sxph* | Saxiphilin | Transferrins, Transferrins | Known to bind saxitoxin in diverse vertebrates, including frogs | Liver, intestines, skin | DHQ-administered vs. alkaloid-free lab *Oophaga sylvatica* | Up[f] | RNA-seq, Proteomics | Caty et al. (2019); O'Connell et al. (2021) |
| *vco3*[g] | Cobra venom factor | Complement component C3/C4/C5, Thioester-containing protein (TEP) | Adaptive and innate immunity | Liver, intestines, skin | DHQ-administered vs. alkaloid-free lab *Oophaga sylvatica* | Up | Proteomics | O'Connell et al. (2021) |
| *vtg2* | Vitellogenin-2 | Vitellogenin, Large lipid transfer proteins | Yolk protein precursor crucial for embryonic development | Liver | PTX- and DHQ-administered frogs vs. DHQ-administered *Dendrobates tinctorius* | Down | RNA-seq | Alvarez-Buylla et al. (2022) |

### Literature Cited

[f] Overexpression in the intestines and skin, but not liver, was significant

[g] Functional analog of C3

Alvarez-Buylla A, Payne CY, Vidoudez C, Trauger SA, O'Connell LA. Molecular physiology of pumiliotoxin sequestration in a poison frog. PLoS One. 2022:17(3), e0264540. https://doi.org/10.1371/journal.pone.0264540.

Alvarez-Buylla A et al. Binding and sequestration of poison frog alkaloids by a plasma globulin. Elife. 2023:12, e85096. https://doi.org/10.7554/eLife.85096.

Caty SN, Alvarez-Buylla A, Vasek C, Tapia EE, Martin NA, McLaughlin T, Golde CL, Weber PK, Mayali X, Coloma LA, Morris MM, O'Connell LA. Alkaloids are associated with increased microbial diversity and metabolic function in poison frogs. Curr Biol. 2025:35(1), 187-197.e8. https://doi.org/10.1016/j.cub.2024.10.069.

O'Connell LA et al. Rapid toxin sequestration modifies poison frog physiology. J Exp Biol. 2021:224(Pt 3), jeb230342. https://doi.org/10.1242/jeb.230342.
