## Supplemental Table 6 for "From diet to defense: serpin duplication and gene-expression evolution as putative contributors to poison-frog alkaloid sequestration"

**Table S6.** This table lists the superfamilies derived from the literature on vertebrate metabolism, representing one of the two lists used to identify candidate DEGs; examples of proteins/protein families belonging to the superfamily; and the primary function in different stages of what the field of pharmacology considers to be the phases of xenobiotic metabolism.

| Protein Superfamily | Example Proteins/Protein Families | Primary Function |
|---|---|---|
| Cytochrome P450s (CYPs) | CYP1A1, CYP2D6, CYP3A4 | Phase I oxidation/hydroxylation of xenobiotics (including alkaloids), phase II (conjugation); metabolic activation or detoxification (Celander and Förlin 1995; Nebert and Dalton 2006) |
| Glutathione S-transferases (GSTs) | GSTA, GSTM, GSTP | Phase II conjugation of electrophilic toxins with glutathione; protects cells from oxidative/toxic stress (Prysyazhnyuk et al. 2021; Zhao et al. 2023) |
| UDP-glucuronosyltransferases (UGTs) | UGT1A1, UGT2B7 | Phase II glucuronidation; enhances solubility and excretion of alkaloids/xenobiotics (He et al. 2010; Kutsuno et al. 2014) |
| Sulfotransferases (SULTs) | SULT1A1, SULT2A1 | Phase II sulfation; conjugates xenobiotics, hormones, and alkaloids for excretion (Glatt and Meinl 2024) |
| ATP-binding cassette transporters (ABCs) | ABCB1 (MDR1), ABCC1 (MRP1), ABCG2 (BCRP) | Phase III efflux pumps; export xenobiotics/alkaloids across membranes; protect tissues from toxic buildup (Ghanem and Manautou 2022) |
| Solute carriers (SLCs) | SLC22 family, SLC15, SLC7 | Import/transport of cations, anions, |

| | | |
|---|---|---|
| | | peptides; some mediate alkaloid uptake into cells (Schumann et al. 2020) |
