## Supplemental Methods and Results for "From diet to defense: serpin duplication and gene-expression evolution as putative contributors to poison-frog alkaloid sequestration"

### Supplementary Methods

*Alkaloid Extraction and Quantification*

For the alkaloid analysis, we extracted skin tissues from two species from trace-accumulating dendrobatid lineages (*Hyloxalus awa* and *Silverstoneia flotator*) and two species from the sequestering clade *Epipedobates* (*E. anthonyi* and *E.* aff. *espinosai*). *Silverstoneia flotator* (*N* = 3) were collected in Soberanía National Forest close to Pipeline Rd., near Gamboa, Panama in August 2022. The remaining species were collected in Ecuador in May and June of 2023: *H. awa* (near Pedro Vicente Maldonado [*N* = 4]); *E. anthonyi* (Moromoro [*N* = 3], Zapotillo [*N* = 5], San Rafael de Sharug [*N* = 4], and Santa Martha [*N* = 3]); and *E.* aff. *espinosai* (Pedro Vicente Maldonado [*N* = 3], Pachijal [*N* = 3], and Caimito [*N* = 4]) (supplementary table S3b). Where possible, the same individuals used in the liver gene expression analysis were profiled for alkaloids, with the following exceptions: (1) skins from the four *H. awa* used for gene expression were not available for this study, so four other individuals collected from the same population on the same day were used instead; nonetheless, a prior, coarser GC–MS analysis of skins from the four *H. awa* whose liver gene expression was profiled was consistent, revealing no alkaloids in the *H. awa* but recovering alkaloids in sympatric *E.* aff. *espinosai* (*unpublished*); (2) skin from a Moromoro individual whose gene expression was profiled (TNHCFS08709) was unavailable, and only the three remaining individuals used for gene expression analysis were profiled for alkaloids; (3) skins from two Santa Martha individuals (TNHCFS08737 and TNHCFS08746) were unavailable, but an additional sample (TNHCFS08744) was included.

Partial dorsal skin samples from *Epipedobates* were collected in 90% EtOH in glass vials and stored in a –80ºC freezer. The extract was syringe filtered through a 0.45-µm PTFE filter (Thermo Scientific, 44504-NP) into a new glass vial, vortexed, and stored for 24h at −80ºC to precipitate lipids and proteins. Skins were removed, freeze-dried overnight in a lyophilizer, and weighed on an analytical balance. After precipitation, the supernatant was filtered again through a 0.45-µm PTFE syringe filter into a new glass vial. The extract was then evaporated completely under $N_2$ gas and resuspended in 50µL ethyl acetate. All resuspended extracts were next transferred to an autosampler vial with a 250µL limited volume insert (Thermo Scientific, 03-452-330) and analyzed by electron impact–mass spectrometry (EI–MS) over a short time range (24h), with several samples re-run to verify that ion abundance was stable among samples.

Ethyl acetate extracts were then re-evaporated at room temperature and resuspended in 10µL ethyl acetate, and re-run on the instrument with both EI–MS and chemical ionization–mass spectrometry (CI–MS). Due to a column replacement between the 50µL and 10µL runs, a cubic spline was calculated to match pre- and post-column change retention times and identify the same alkaloids in both runs. A Thermo AS 3000 autosampler was used to inject 1µL of extracts into a Thermo Scientific TSQ Quantum mass-spectrometry instrument interfaced with a Trace GC Ultra gas chromatograph, with a polar Stabilwax GC Capillary MS Column, 30 m, 0.25 mm ID, 0.25 µm (Restek 10623). The separation of alkaloids was achieved with helium as the carrier gas (flow rate: 1 mL/min) using a temperature program increasing from 60 to 260°C at a rate of 10°C/min.

To ensure consistent depth (unbiased) searching for alkaloids across samples, a batch-processing pipeline was implemented for all samples run in EI at both concentrations (50µL and 10µL) in which spectra from total ion current (TIC) peaks identified by the Thermo Xcalibur software were searched against the NIST main reference library database and top-10 hits were retained from all searches. Then, for all EI spectra for which anuran alkaloids were returned within the top-10 hits, both EI and corresponding CI spectra were deconvoluted and inspected manually and compared against published (e.g., Daly et al. 2005, 2009) and archival data to confirm the identification as an alkaloid and determine class, obtain molecular weights, and determine the precise names. When an anuran alkaloid was identified during batch processing, all other samples from the same population were checked for the same alkaloid, and these alkaloids too were retained for further analysis.

Extractions for both *S. flotator* and *H. awa* had more starting material (whole dorsal skin samples and whole dorsal and ventral skins from *H. awa*), stronger extraction solvents (*S. flotator* were stored in 100% MeOH and *H. awa* in 98% EtOH) and in the case of *S. flotator*, a more sensitive extraction into 90% MeOH intended for a higher-throughput spectrometry method—ultra-high-performance liquid-chromatography heated electrospray-ionization tandem mass spectrometry (UHPLC–HESI–MSMS) (Tarvin et al. 2024; Coleman et al. 2025). Despite this and the use of bench and analytical methods identical to those for *Epipedobates* (evaporation and resuspension in 10µL ethyl acetate, injection of 1µL of extracts into the Thermo Scientific TSQ Quantum mass spectrometry instrument), the batch-processing pipeline recovered no poison-frog alkaloids within the top-10 hits for spectra from TIC peaks for either species. Thus, skin extracts from the two species were not subject to further instrument runs.

A system with a nonpolar column—necessary for retention time and Kovats retention index determination—was also used for further confirmation of identities of alkaloids from *Epipedobates*. An Agilent 7693A GC autoinjector (G4513A) was used to inject 1μL of the 10μL extracts into an Agilent Technologies 5977E Single Quadrupole mass spectrometry instrument interfaced with an Agilent Technologies 7820A GC system, with a DB-5ms Low Bleed GC Column, 60 m, 0.25 mm, 0.25 μm, 7 inch cage (Agilent 122-5562). Confidently-identified alkaloids (>80% library match and already confirmed manually based on EI and CI spectra) were normalized to literature data using linear regression. Alkaloids were then matched between Agilent and Thermo systems where obvious based on relative abundance and mass spectra for retention-time determination. Kovats retention indexes (semi-standard nonpolar) were also determined for the same spectra based on a run of alkanes 9–30 (in hexane).

Integrated areas of the base peak chromatograms of all alkaloids recovered from batch processing of the dilute batch (50μL) were calculated. Then, alkaloids batch-processed from the concentrated batch but not batch-processed in the dilute batch were recovered from the chromatograms of the dilute batch and integrated areas calculated for these compounds. One compound batch-processed in the concentrated batch did not have a peak present in the dilute batch (allopumiliotoxin 323B [aPTX 323B]). For concentrated samples in which this alkaloid was recovered, linear regression equations correlating integrated areas of alkaloids in the concentrated and dilute sample in which aPTX 323B was present were determined, and concentrations of aPTX 323B for the dilute samples were back-predicted. The abundance values (ion counts) from the dilute sample chromatograms were normalized by dividing by the dry skin weight. Finally, all chromatograms were exhaustively searched manually for alkaloids by inspecting all TIC peaks and extracting an array of common alkaloid base peaks, according to Daly et al. (2005) (supplementary table S15).

*Phylogenetic Reconstruction and Analysis of the SerpinA Gene Family in Dendrobatids*

Recent proteomic studies revealed that members of the serpinA gene group *serpina1*-like encode ABGs in *O. sylvatica*, *D. tinctorius*, and *E. tricolor* (Alvarez-Buylla et al. 2023). To elucidate the evolution of the *serpina1*-like gene group, we carried out a phylogenetic study of this group and included in our analysis other serpinA gene subfamilies (e.g., *serpina5*, *serpina6*, *serpina7*, and *serpina10*) for completeness. We surveyed published amphibian genomes (N = 37 species), transcriptomes (N = 4), and PCR sequences (N = 10) for serpinA genes, and we used *Latimeria chalumnae* (GCF_037176945.1) as a taxonomic outgroup. All relevant information on these sequences is provided in supplementary table S1, which includes taxonomic classification, source (i.e., genomic, transcriptomic, PCR), tissue (i.e., gut/intestine, liver or skin), reclassification within BBS/BBS-like and ancestral A1AT/*serpina1*-like/ABG groups, number of variants, NCBI/SRA accession numbers, metadata (e.g., collection site), nucleotide/amino acid sequence alignments, and key AA sites states associated with PTX-**251D** binding by OsABG1 (Alvarez-Buylla et al. 2023), with relevant physicochemical properties.

Our surveyed genomes included 31 anurans (including 25 neobatrachians), three salamanders, and three caecilians. To extract serpinA genes from the genomes, we retrieved all annotated predicted transcripts generated as individual sequences or as Coding DNA Sequences (CDS) within chromosome scaffolds by the *NCBI Eukaryotic Genome Annotation Pipeline* (NCBI EGAP) version 10.4, which uses a combination of Gnomon, cmsearch/Infernal, and tRNAscan-SE software (Nawrocki and Eddy 2013; Thibaud-Nissen et al. 2013; Chan et al. 2021). We searched using key names including "A1AT" or "A1AP" (alpha-1 antitrypsin), "serpinA", "serpin family A" , "serpina-like," "BBS," "hylaserpin," and variants of these terms. We found widespread inconsistency in the naming which included sequences from the same serpin group such as BBS-like genes that have been assigned multiple generic names, including "A1AT" or "serpina1-like." Likewise, some SerpinA members have been characterized with variants of their common nomenclature such as for *serpina5*-like (protein C inhibitor, PCI), *serpina6*-like (corticosteroid-binding globulin, CBG), *serpina7*-like (thyroxine-binding globulin, TBG), and *serpina10*-like (Protein Z-dependent protease inhibitor, ZPI).

We aligned the retrieved sequences with a codon-informed method using DECIPHER version 3.0.0 (Wright 2016) with AlignTranslation, followed by phylogenetic reconstruction using IQ-TREE version 3.0 (Wong et al. 2026). The best-fit substitution models and partitioning scheme were selected using ModelFinder with the MFP + MERGE option. Nodal support was also estimated in IQ-TREE using the ultrafast bootstrap approximation (UFBoot) with 1,000 replicates, followed by optimization with the --bnni parameter to reduce support overestimation. We rooted all phylogenies with *serpina10*-like (ZPI) as the outgroup, and we renamed all recovered serpinA members according to their corresponding phylogenetic groups for further analyses and discussion. Due to the aforementioned challenges with nomenclature, we harmonized gene names for the full serpinA sequence set by performing protein similarity searches against the human NCBI ClusteredNR database, a reduced-redundancy version of the NCBI non-redundant protein database generated using MMSeqs2 by clustering proteins at >90% sequence identity and within 90% of the length of the longest cluster member. Each cluster is represented by a selected well-annotated protein sequence, reducing redundancy and improving taxonomic breadth in BLAST results. We repeated the phylogenetic analyses with the extended dataset derived from transcriptomes, as indicated below.

*SerpinA Transcript Isolation from* de novo *Assemblies and Targeted Read Baiting*

We focused on transcriptomes from *O. sylvatica*, *E. anthonyi*, *E. tricolor*, and *Mantella aurantiaca*. First, we reannotated the *M. aurantiaca* transcriptome (Dryad doi:10.5061/dryad.mkkwh7143) with the genomic references for serpinA as obtained from the genome repositories. Second, we reassembled and reannotated published transcriptomes of *O. sylvatica* from raw reads found in the NCBI-SRA with their corresponding SRR numbers (supplementary table S1) and Dryad repositories (i.e., doi:10.5061/dryad.4cs9573 and doi:10.5061/dryad.mkkwh7143). Third, *Epipedobates* data were generated *de novo*, but focused on a targeted reconstruction of only ABGs and BBSs transcripts using a sequence bait, *de novo* assembly, and their annotation using PCR-based dendrobatid ABGs (i.e., NCBI OQ032869–71), and *O. sylvatica* transcriptome and genomic references.

For the SRA data, raw RNA sequence reads were processed using the Pincho version 0.1 modular pipeline (Ortiz et al. 2021), which includes an integrated cleaning, multi-assembly construction, consensus assembly quality assessment, and automated annotation. Briefly, Pincho automated *de novo* assembly construction with the following steps: (1) raw data download from SRA followed by removal of Illumina adapter sequences with Trimmomatic version 0.39 (Bolger et al. 2014); (2) sequence insert error correction with Rcorrector version 1.0.4 (Song and Florea 2015); (3) three independent assembly reconstructions with trans-ABySS version 2.0.1 (Robertson et al. 2010) using five k-mer sizes (i.e., 21, 39, 59, 79, and 99), rnaSPAdes version 3.14.1 (Bushmanova et al. 2019), and TransLig version 1.3 with kmer = 31 (Liu et al. 2019); (4) combined consensus transcriptome assembly using TransRate version 1.0.3 (Smith-Unna et al.

2016); (5) removal of identical transcripts with CD-HIT v4.8.1 (Fu et al. 2012); (6) consensus transcriptome quality assessment with BUSCO version 6.0.0 (Tegenfeldt et al. 2025) for the eukaryota_odb12 dataset; for *Oophaga*, the resulting number of raw reads ranged from 58–309 M with BUSCO scores from 91.0–99.7% (supplementary table S2a); and (7) transcript annotation against both 2026 updated TrEMBL:Amphibia and UniProt: Swiss-Prot with BLASTx v2.10.0+ (Altschul et al. 1990) with acceptance e-value of 1e-10. For each transcript with a sequence database match, annotation was assigned based on the top reference hit, prioritizing the longest aligned sequence with the lowest e-value. We then applied the same keyword-based data-mining procedure used for genomes, retaining only transcripts whose annotations matched our serpinA search terms for further analysis. From these transcripts, we extracted the longest coding sequence using TransDecoder version 6.0.0 (Haas and Papanicolaou 2019) with the TransDecoder.LongOrfs module and generated nucleotide CDS sequences from the resulting longest_orfs.cds output.

We collected one individual of *E. tricolor* (Ecuador: Bolivar: Chazojuan) and two *E. anthonyi* (Ecuador: Azuay: Santa Isabel) following the St. John's University IACUC (AUP 1965) protocol for animal handling and tissue harvest (i.e., skin, liver, and intestines) in RNAlater (Invitrogen/Thermo Fisher Scientific, Waltham, MA, USA). Total RNA was extracted using the TRIzol protocol after tissue homogenization, followed by RNA integrity evaluation, mRNA isolation, and directional library construction using a published protocol (Ortiz et al. 2025). These *Epipedobates* samples were then sequenced (150 bp paired-end), producing approximately 70–97 M Illumina read pairs per sample. Transcriptomes were assembled using Pincho, and assemblies had BUSCO scores spanning 85.4–99.2% (supplementary table S2a). Raw RNA-seq reads from *Epipedobates* samples were processed for targeted reconstruction of ABG and BBS transcripts using a modified Pincho pipeline (Ortiz et al. 2021) that incorporated read mapping and baiting prior to transcriptome assembly. We focused targeted reconstruction on ABG- and BBS-like serpinA transcripts because existing annotations and preliminary searches indicated unusually high paralog/variant diversity in these clades. Briefly, the workflow consisted of the following steps: (1) raw Illumina reads were cleaned and error-corrected using Pincho*;* (2) reads were baited using ABG and BBS sequences identified from the *O. sylvatica* transcriptome, dendrobatid genome references, and previously published ABGs. Read baiting was performed with BBDuk from BBTools/BBMap version 35.85 (Bushnell 2014) using k-mer filtering parameters selected for high specificity and sensitivity: k = 31, hdist = 0, qhdist = 0, and minkmerhits = 2; (3) baited reads were assembled using the Pincho transcript-assembly workflow, followed by redundancy reduction, consensus construction, and BLASTn annotation against SerpinA genes isolated from *Oophaga* and other dendrobatid genomes. To validate the baiting pipeline, we also reconstructed 11 broadly expressed housekeeping genes as references (Vandesompele et al. 2002; Bustin et al. 2009; Kozera and Rapacz 2013; Molina et al. 2018): *actb*, *eef1a1*, *gapdh*, *hmbs*, *hprt1*, *ipo8*, *polr2a*, *rpl13a*, *rplp0*, *tbp*, and *ubc*. Sequence, NCBI, and SRR accession numbers are provided in supplementary table S2c.

*Quality Evaluation of SerpinA Transcript Reconstructions*

After transcript reconstruction, we evaluated the quality of each serpinA sequence recovered from *Oophaga* and *Epipedobates* using Bowtie2 version 2.5.5 (Langmead and Salzberg 2012), SAMtools version 1.23.1 (Li et al. 2009), mosdepth version 0.3.11 (Pedersen and Quinlan 2018), and Salmon version 2.1.1 (Patro et al. 2017). Read-depth and transcript-completeness analyses did not include transcriptomic data from *Mantella aurantiaca*, because the tissue of origin was uncertain and likely mixed. Baited paired-end reads from BBDuk were mapped back to their corresponding serpinA transcript reconstructions to assess coverage depth, mapping ambiguity, and transcript completeness. First, we generated a Bowtie2 index for each serpinA reconstruction and mapped baited reads to produce SAM files. These files were converted to sorted and indexed BAM files, and mapping statistics were summarized with SAMtools, including reference length (startpos, endpos), number of aligned reads (numreads), covered bases (covbases), percent coverage (coverage), mean depth (meandepth), mean base quality (meanbaseq), and mean mapping quality (meanmapq). Second, we used mosdepth to calculate regional coverage across each reference transcript using --no-per-base and --by reference BED intervals derived from SAMtools. Transcript completeness was evaluated as the percentage of transcript length covered at depth thresholds of ≥1x, ≥5x, ≥10x, and ≥20×. Third, we estimated transcript abundance with salmon quant, using the baited paired-end reads from BBDuk for each species-specific serpinA transcript set with --validateMappings to enable selective-alignment validation (Srivastava et al. 2020). This produced estimates of effective length, transcripts per million (TPM), and estimated read counts (NumReads). We repeated the Salmon quantification using the full collection of reads for each SRA and *Epipedobates* sample; because the results were comparable, only the baited-read mapping results are presented in supplementary table S2a.

We interpreted serpinA transcript-reconstruction quality using the following criteria. For coverage depth and completeness, we considered reconstructions well supported when mosdepth coverage at ≥10× exceeded 90% completeness, SAMtools percent coverage was 100%, mean depth exceeded 10, and mean base quality exceeded 35. To evaluate mapping uniqueness to reconstructed transcripts, we classified SAMtools mean mapping quality into the following confidence bins: <5, ambiguous or multi-mapping; 5–9, weak confidence; 10–19, medium confidence; 20–29, high confidence; and ≥30, excellent confidence consistent with unique mapping. From the Salmon results, we used TPM and estimated read counts to compare read support for serpinA reconstructions against the 11 reference housekeeping genes and to assess whether each transcript was supported in a given tissue-derived dataset. We did not use these values to infer differential gene expression; rather, they were used only to evaluate transcript recovery, read-mapping support, and tissue-level evidence for transcript presence.

*Additional Phylogenetic Reconstruction*

We performed an additional phylogenetic analysis in which we partitioned serpinA model inference by codon position, to confirm the robustness of the analysis we present in the main text. We aligned trimmed sequences (1,173 bp, see main text) using MUSCLE version 3.8.31 (Edgar 2004), applying default settings, and we corrected small errors in alignment and determined codon position under the standard genetic code in Mesquite version 4.02 (Maddison and Maddison 2025). We constructed multiple ML trees with combinations of parameters (1,000 or 2,000 ultrafast bootstrap replicates, 1, 5, and 20 runs, radius of 60 or 150) using IQ-TREE version 3.0.1 (Wong et al. 2026), implementing ModelFinder (Kalyaanamoorthy et al. 2017) to find the preferred model of molecular evolution for each data partition (codon position) under the Bayesian Information Criterion, with subsequent merging (Chernomor et al. 2016). Supplementary trees were cosmetically curated using the Interactive Tree of Life version 7 (Letunic and Bork 2024) and presented as unrooted.

*ABG Structure Prediction and Putative Binding Site Computation*

Predicted structures for OsABG1 and OsABG2 were generated from full-length amino-acid sequences using AlphaFold Server with default settings, with each sequence modeled as a single protein chain. To summarize amino-acid divergence at putative alkaloid-binding sites, we extracted residues corresponding to the six OsABG1 binding-pocket positions identified by Alvarez-Buylla et al. (2023) from the serpinA amino-acid alignment. For each sequence, Grantham distances were calculated between the observed residue and the OsABG1 reference residue at each position and then summed across the six sites to generate the total Grantham distance shown in Fig. 2. A gap at any of these positions was assigned the maximum Grantham distance for that site.

*Read Processing, Alignment, and Expression Quantification*

Tag-Seq raw reads were preprocessed prior to alignment using custom Perl scripts (Meyer et al. 2011), FASTX-Toolkit version 0.0.14 (Gordon and Hannon 2010), and Cutadapt version 2.8 (Martin 2011). Steps were as follows: (1) reads with homopolymer A runs greater than eight bases were removed; (2) reads were retained only if they had at least 20 bases after trimming; (3) PCR duplicates were removed when sequences shared the same degenerate header and first 20 bases; and (4) reads were filtered for quality by retaining reads with Phred quality scores greater than 20 across at least 90% of nucleotides. FastQC version 0.12.1 (Andrews 2010) was used for quality control to confirm the absence of low-quality bases and sequence biases, and that we had successfully removed any contaminating adapters during pre-processing. MultiQC version 1.19 (Ewels et al. 2016) was used to compile the FastQC reports into a summary for visualization. We built a Salmon index from the *O. sylvatica* reference transcriptome and quantified each Tag-Seq library using Salmon version 1.10.2 (Patro et al. 2017) to obtain transcript-level TPM and count estimates. Quantifications were imported into R version 4.3.3 (R Core Team 2025) using the tximport package (version 1.30.0; Soneson et al. 2015) and merged into count and abundance matrices using a sample-level metadata table. Separately, reads were aligned to the *O. sylvatica* reference transcriptome using BWA version 0.7.17-r1188 (Li and Durbin 2009). Mapping percentages were then evaluated among species. An ANOVA followed by a TukeyHSD test was implemented to statistically evaluate differences in mean mapping percentages among species. Unless otherwise specified, all subsequent analyses were performed in R.

*Principal Component Analysis (PCA) and Phylogenetic PCA (pPCA)*

PCA reduces the dimensionality of high-dimensional datasets by constructing new uncorrelated variables (principal components, or PCs) that successively maximize variance explained. To explore global patterns in gene expression, we performed a PCA on variance-stabilized data for the 940 shared genes that survived filtering, implementing the prcomp function (R Core Team 2025) using the correlation matrix (supplementary fig. S7a). Filtered and voom-normalized gene expression data were used as input. We extracted PC scores for each sample and calculated the percentage of variance explained by each PC by dividing the squared standard deviation of each PC by the total sum of squared standard deviations across PCs. We plotted samples along PC1 and PC2. We also inspected genes with the largest absolute PC1 loadings to determine which genes were most strongly associated with PC1. To determine the optimal number of clusters in PC space, we performed k-means clustering on PC1 and PC2 coordinates across $k = 1$ to 10 clusters and evaluated the total within-cluster sum of squares using an elbow plot (supplementary fig. S6). Statistical differences in PC1 scores between sequestering and trace-accumulating categories were assessed using a Welch two-sample t-test, with significance determined at $\alpha = 0.05$.

We ran pPCAs using the phyl.pca function in the R package phytools version 2.1-1 (Revell 2024) under a Brownian motion model of trait evolution. This approach accounts for expected covariance among species arising from shared evolutionary history. Under this model, the covariance structure of traits is transformed according to the phylogenetic variance-covariance matrix before PCs are calculated. We conducted the pPCA using both covariance (cov) and correlation (corr) matrices to confirm consistency between results, because PCA on the correlation matrix is identical to PCA on standardized variables. Covariance-based PCA emphasizes absolute variance and is therefore more sensitive to strongly expressed outlier genes, whereas correlation-based PCA emphasizes relative patterns of variation among variables (Revell 2024).

We first evaluated whether phylogenetic structure at the individual or population level could disproportionately influence variance in overall gene expression patterns. Population-level structure was only relevant for *Epipedobates*, from which multiple populations were sampled for both species. To assess these effects, we repeated the pPCA across three topologically equivalent phylogenetic trees that differed only in branch lengths. We conducted two pairs of correlation-based and covariance-based pPCAs in which in the underlying tree, both populations and individuals were given phylogenetic structure, but for which we varied branch lengths for species from the true median species divergence time (estimated from a posterior distribution; López-Hervas et al. 2024) to the same arbitrary value given to populations and individuals (0.1). This was to verify that both pairs of plots were concordant, i.e., that phylogenetic structure among species, and not divergence times among populations or expression variance among individuals, account for any phylogenetic signal. To generate the pPCA, we used (1) two trees retaining the real estimates of posterior median ages for species but with reduced branch lengths (0.1) for populations and individuals (supplementary fig. S7b, c); and (2) two trees with uniform branch lengths of 0.1 applied to all branches, including internal nodes (supplementary fig. S7d,e).

We also constructed two pairs of covariance- and correlation-based trees in which (a) populations, but not individuals and (b) individuals, but not populations, were given structure (treated as evolutionary units): in these two tree pairs, we retained real median posterior age estimates among species, assigned reduced branch lengths (0.1) to populations (first pair) or individuals (second pair), and assigned near-zero terminal branches to individuals (first pair) or populations (second pair) to approximate a star polytomy, i.e., assuming shared mean expression among individuals within a population (supplementary fig. S7f–i). Median posterior ages were 1.99 mya (between *Epipedobates*), 13.91 mya (between *Epipedobates* and *Silverstoneia*), and 33.04 mya (between Colostethinae [*Epipedobates* + *Silverstoneia*] and *Hyloxalus*) (López-Hervas et al. 2024). For the above pPCAs, individual samples were treated as tips, with population membership encoded by appending sequential integers to population labels to create unique identifiers. Phylogenetic PC scores were extracted from the S component of the phyl.pca output, and the percentage of phylogenetic variance explained by each PC was calculated as the proportion of each eigenvalue relative to the total sum of eigenvalues, multiplied by 100. Eigenvalues and eigenvectors were coerced to their real PCs prior to analysis to handle residual imaginary values arising from numerical operations.

Last, we conducted correlation-based and covariance-based species mean-expression pPCAs, in which phylogenetic structure was imposed only on species, with expression values averaged across individuals within each species. Individual samples were then projected onto the species-level pPCA axes using the scores function to obtain individual-level PC projections. The percentage of variance explained at the individual level was calculated from these projected scores and reported alongside the phylogenetic variance explained by each axis. Biplots for both the correlation- and covariance-based pPCAs are displayed as supplementary figures (supplementary fig. S7j,k).

#### *Candidate Gene Selection and Heatmap Visualization*

Candidate DEGs were defined as genes that (1) were differentially expressed between sequestering and trace-accumulating species in the limma analysis and (2) belonged to literature-curated gene superfamilies whose members have been directly implicated in dendrobatid alkaloid metabolism (supplementary table S5) or are known or hypothesized to participate in vertebrate liver toxin handling, including detoxification, clearance, absorption, biotransformation, and immune response (supplementary table S6). To visualize relative expression patterns of candidate DEGs across samples and among alkaloid categories, we generated hierarchically clustered heatmaps using the R package pheatmap version 1.0.13 (Kolde 2025). Expression matrices were constructed from filtered, voom-normalized, log-transformed counts. For heatmap visualization, gene expression values were row-scaled (Z-score transformed) to facilitate comparison across samples. Clustering was performed on both rows (genes) and columns (samples) using Euclidean distance and complete linkage (Fig. 5).

#### *WGCNA*

We conducted a WGCNA using the WGCNA R package version 1.73 (Langfelder and Horvath 2008). WGCNA identifies clusters of genes with highly correlated expression patterns (modules) by transforming gene-gene correlation matrices into adjacency matrices using a soft-thresholding approach, which preserves strong biological correlations while suppressing weak or noisy correlations. To determine the optimal soft-thresholding power ($\beta$), we evaluated powers ranging from 1 to 20 and selected the lowest power that achieved a scale-free topology model fit ($R^2 \geq 0.85$). This soft-thresholding power was then applied to transform the Pearson correlation matrix into a weighted adjacency matrix. We subsequently calculated a topological overlap matrix (TOM) from the adjacency matrix, which incorporates both direct and indirect gene-gene relationships to capture higher-order connectivity patterns within the network. Genes were hierarchically clustered based on TOM-based dissimilarity (1 - TOM), and modules were identified using the blockwiseModules function with a minimum module size of 15 genes and an unsigned network type. Modules are distinguished by colors for visualization and downstream analysis.

To assess the biological relevance of identified modules, we calculated MEs, defined as the first principal component of each module's expression profile, which summarizes the expression pattern of all genes within a module. We tested associations between MEs and alkaloid-phenotype category (sequestering or trace-accumulating) using a Pearson correlation and assessed statistical significance using Student asymptotic p-values, with BHFDR correction applied across all module-trait tests. Modules significantly associated with sequestration were subsequently tested for species-level differences in ME values using pairwise Welch two-sample t-tests, to evaluate whether species identity contributed additional structure to ME variation beyond phenotypic category. We also performed Fisher's exact tests to assess whether modules were significantly enriched for DEGs previously identified by limma. Four modules were selected for further investigation because they were significantly enriched for DEGs (FDR $\leq$ 0.05) and displayed significant ME-phenotypic-category correlations (FDR $\leq 0.05$). For genes within these modules, we calculated module membership (kME), defined as the correlation between each gene's expression profile and its module eigengene, to identify highly connected intramodular hub genes. Genes were retained for network visualization if they were differentially expressed (adjusted $p \leq 0.05$ from limma) or exhibited high module membership (kME $\geq 0.6$).

For network construction and visualization, we exported module-specific gene networks to Cytoscape (Shannon et al. 2003) by calculating TOM-based edge weights between genes within each prioritized module. Edges with TOM values below 0.02 were excluded before export to Cytoscape using the exportNetworkToCytoscape function in WGCNA, after visual inspection confirmed that this threshold excluded gene pairs with uninformative (noisy) connections. Edges falling below the 50th percentile of remaining edge weights were hidden for reduced visual clutter while still remaining in the underlying network file. Node attributes included module assignment, log-fold change from the differential expression analysis, and module membership values (supplementary figs. S12–S15).

### Supplementary Results

*Alkaloid Extraction and Quantification*

A diverse profile of poison-frog alkaloids was recovered in *Epipedobates* (supplementary tables S4a–g, S15; supplementary figs. S3, S4), while we failed to detect alkaloids in *S. flotator* and *H. awa*. Across *Epipedobates*, the number of alkaloids varied among populations, ranging from eight in ZAP to forty in MAR, with a mean of 24.86 ± 12.64 SD per population. In total, the batch-processing pipeline recovered 117 distinct alkaloids, including stereoisomers, representing eighteen anuran structural classes: decahydroquinolines (four), octahydroquinolines (three), 3,5-disubstituted indolizidines (three), 5,8-disubstituted indolizidines (thirty-two), dehydro-5,8-indolizidines (two), 5,6,8-trisubstituted indolizidines (fourteen), homopumiliotoxins (two), desmethylhomopumiliotoxins (one), allopumiliotoxins (three), pumiliotoxins (nine), dehydrodesmethylpumiliotoxins (two), deoxyhomopumiliotoxin (one), deoxypumiliotoxins (thirteen), 1,4-disubstituted quinolizidines (four), 3,5-disubstituted pyrrolizidines (eight), histrionicotoxin (one), piperidine (one), and tricyclics (three).

Alkaloid composition also varied among *Epipedobates* populations. Some classes, such as 5,8-disubstituted indolizidines, were widespread across populations, whereas others appeared enriched in particular species or populations, including deoxypumiliotoxins in *E. anthonyi*, allopumiliotoxins in *E.* aff. *espinosai* CTO, and tricyclics in *E. anthonyi* ZAP (supplementary fig. S3). For each individual, we calculated log-transformed total alkaloid abundance as the natural log of the summed integrated peak areas for identified alkaloids (Fig. 4). Total alkaloid abundance differed markedly between the sequestering *Epipedobates* and the trace-accumulating *Silverstoneia* and *Hyloxalus* (Fig. 4).

Our findings are congruent with previous surveys showing that *Epipedobates* is a uniformly alkaloid-rich clade (e.g., Cipriani and Rivera 2009; Tarvin et al. 2024; Caty et al. 2025), whereas members of *Hyloxalus* and *Silverstoneia* typically contain only trace concentrations of few alkaloids, if any (e.g., Gonzalez et al. 2021; Martin et al. 2025). For example, Coleman et al. (2025) characterized alkaloid profiles of 12 individuals of *S. flotator*, which included all three individuals sampled here, and detected only 0–1 putative poison-frog alkaloids per individual (precise identification is not possible with LCMS given lack of MS2 spectral references) and only four of these alkaloids were detected in total, despite use of a more sensitive UHPLC–HESI–MS/MS pipeline (Coleman et al. 2025). Likewise, a previous sensitive GC–MS workflow based on concentrated crude methanol extracts followed by manual characterization and quantification found 0–12 trace alkaloids per individual in seven *H. awa* (Tarvin et al. 2024).

*Quality Evaluation of SerpinA Transcript Reconstructions*

Results from these analyses supported the recovery of most reconstructed serpinA transcripts, with many targets showing high coverage breadth, high read depth, and high base quality. The reference genes showed consistently strong coverage and high mapping quality, indicating that the read-mapping workflow recovered broadly expressed transcripts cleanly. In contrast, ABG- and BBS-like serpinA transcripts frequently showed low mean mapping quality despite strong coverage support, consistent with multi-mapping of reads among closely related paralogs and transcript variants.

*Phylogenetic Reconstruction*

Findings from the additional phylogenetic analyses were concordant with each other and almost entirely concordant (with major clades well supported in both cases) with the phylogenetic results presented in the main text, except for the location of *Anomaloglossus baeobatrachus* ABGs; these two sequences were resolved in the main-text analysis with low certainty as sister to the ABG1 + ABG2 clade, whereas we find that, when we partition by codon position, these sequences come out as basal within the ABG1 clade, an expected phylogenetic position. We present the best tree from one analysis as supplementary fig. S2 (-m TESTMERGE -mfreq FO -mrate G --runs 1 -B 1000 --radius 60). Main text figures (Figs. 2, 3) and supplementary fig. S1 are based on the phylogenetic methods/results described in the main text, for which *A. baeobatrachus* ABG1 is non-monophyletic.

Codon partition 1 corresponded to the first codon position in the open reading frame. Maximum-likelihood values of best trees converged around –170485 log-likelihood. For this run, the following were the best-fit models per partition for positions/partitions 1–3, respectively: TIM3+FO+G4, GTR+FO+G4, and GTR+FO+G4.

*ABG Structure Prediction and Putative Binding Site Computation*

AlphaFold Server returned five predicted models per sequence; model 0 was visualized in PyMOL version 3.1 (Schrödinger LLC 2025). Residues corresponding to the six OsABG1 binding-pocket sites identified by Alvarez-Buylla et al. (2023) were highlighted on each predicted structure. See the main text (Fig. 2) for Grantham results and interpretation.

*PCA and pPCA: Overall Differences in Gene Expression*

Variation among individuals in gene expression clustered into three groups corresponding to the three sampled genera. An elbow plot of k-means clustering supported three clusters based on PC1 and PC2 scores from the PCA of gene expression data (supplementary figs. S6, S7a). The primary axis of transcriptomic variation (PC1, 25.3% variance explained) captured broad expression differences among genera, with *Epipedobates*, *H. awa*, and *S. flotator* widely separated, as well as subtler differences between *E. anthonyi* and *E.* aff. *espinosai* (supplementary fig. S7a). PC2 (12.0%) separated *H. awa* from the genera within Colostethinae (*Epipedobates* and *Silverstoneia*). Incidentally, although the per-population sample sizes were too low to support strong inference, *E. anthonyi* RAF did not cluster more closely with the geographically proximate MAR population than with other *E. anthonyi* populations along PC2, despite little evidence of neutral genetic differentiation between RAF and MAR (Fig. 1b; Páez-Vacas et al. 2022).

Plots of phylogenetic PCAs (pPCAs) conferring phylogenetic structure to individuals and populations jointly (supplementary fig. S7b–e), to populations only (supplementary fig. S7f,g), or to individuals only (supplementary fig. S7h,i) showed strong concordance, with samples largely overlapping rather than forming discrete clusters (Supplementary Methods). These results suggest that population- or individual-level sampling did not contribute disproportionate gene expression variation; individuals did not differ from one another more than expected given genealogical relatedness within populations or evolutionary divergence among populations. We therefore collapsed population- and individual-level variation and conducted a species mean-expression pPCA (supplementary fig. S7j,k). Expression similarity did not track phylogenetic relatedness. For example, PC1 for the correlation-based species-mean PCA (43.93% variance explained) markedly separated the two *Epipedobates* species despite their recent divergence; along this axis, *H. awa* was closer to *E.* aff. *espinosai*, whereas *S. flotator* was intermediate between the two *Epipedobates* species. PC2 (31.56%) separated *S. flotator* from the remaining species, while *H. awa* and both *Epipedobates* species had more similar PC2 scores. Results from the species-mean pPCA indicate substantial species-level expression structure that is not explained solely by phylogenetic relatedness.
